# DNMT1-Mediated Regulation of Somatostatin-positive Interneuron Migration Impacts Cortical Architecture and Function

**DOI:** 10.1101/2024.09.04.611268

**Authors:** Julia Reichard, Philip Wolff, Song Xie, Ke Zuo, Camila L. Fullio, Jian Du, Severin Graff, Jenice Linde, Can Bora Yildiz, Georg Pitschelatow, Gerion Nabbefeld, Lilli Dorp, Johanna Vollmer, Linda Biemans, Shirley Kempf, Minali Singh, K. Naga Mohan, Chao-Chung Kuo, Tanja Vogel, Paolo Carloni, Simon Musall, Geraldine Zimmer-Bensch

**Affiliations:** RWTH Aachen University, Division of Neuroepigenetics, Institute of Zoology (Biology 2), Worringerweg 3, 52074 Aachen, Germany; Research Training Group 2416 MultiSenses – MultiScales, RWTH Aachen University, 52074 Aachen, Germany; Institute of Neuroscience and Medicine (INM-9) Computational Biomedicine, Forschungszentrum Jülich GmbH, 52428 Jülich, Germany; RWTH Aachen University, Department of Physics, 52074 Aachen, Germany; College of Pharmacy (International Academy of Targeted Therapeutics and Innovation), Chongqing University of Arts and Sciences, 402160 Chongqing, PR China; Department of Physics, University of Cagliari, I-09042 Cagliari, Italy; Institute for Anatomy and Cell Biology, Department of Molecular Embryology, Faculty of Medicine, Albert-Ludwigs-University Freiburg, 79104 Freiburg, Germany; Faculty of Biology, Albert-Ludwigs-University Freiburg, 79104 Freiburg, Germany; Institute of Biological Information Processing (IBI-3) Bioelectronics, Forschungszentrum Jülich, Germany; RWTH Aachen University, Division of Neurophysiology, Institute of Zoology (Biology 2), Worringerweg 3, 52074 Aachen, Germany; Molecular Biology and Genetics Laboratory, Department of Biological Sciences, BITS Pilani, Hyderabad Campus, Hyderabad, India; Genomics Facility, Interdisciplinary Center for Clinical Research (IZKF), RWTH Aachen University, 52074 Aachen, Germany

## Abstract

The fine-tuned establishment of neuronal circuits during the formation of the cerebral cortex is pivotal for its functionality. Developmental abnormalities affecting the composition of cortical circuits, which consist of excitatory neurons and inhibitory cortical interneurons (cINs), are linked to a spectrum of neuropsychiatric disorders. Excitatory neurons originate in cortical proliferative zones, while inhibitory interneurons migrate from discrete domains of the basal telencephalon into the cortex. This migration is intricately governed by intrinsic genetic programs and extrinsic cues. Our current study reveals the role of the DNA methyltransferase 1 (DNMT1) in regulating the expression of key genes implicated in mouse cIN development and in guiding the migration of somatostatin (SST)-expressing interneurons at postmitotic level within the developing cortex. *Dnmt1* deletion causes SST^+^ cINs to exit prematurely from the superficial migratory stream. In addition to the perturbed migration pattern and altered gene expression signatures, *Dnmt1*-deficient SST^+^ cINs had a discernible non-cell autonomous effect on cortical progenitors, which culminated in nuanced alterations of layer thicknesses in the adult cortex. Our study uncovers that DNMT1 governs the migration of SST^+^ cINs and through this, their instructive role in sculpting the intricate cortical layer architecture by signaling to cortical progenitors, with pronounced effects on neuronal network function.

## Introduction

The intricate neuronal circuitry of the mammalian cerebral cortex, composed of excitatory and inhibitory neurons, empowers its enormous cognitive capabilities. The circuit formation involves coordinated developmental steps: proliferation, differentiation, migration, morphological maturation, and synapse formation^1^.

Excitatory cortical projection neurons originate from radial glial cells (RGCs) and intermediate progenitors (IPCs) located in the cortical proliferative zones, through direct and indirect neurogenesis. Symmetric divisions of RGCs initially expand the progenitor pool, later shifting to asymmetric divisions to produce neurons and RGCs or IPCs^2;3^. EOMES-expressing IPCs localize to the subventricular zone (SVZ) and generate neurons for all cortical layers after few self-renewal divisions^3^. Cortical excitatory neurons are produced in a time-dependent manner, with deep-layer neurons born first and later-born neurons forming superficial layers, coming with unique functional features and connectivity^4^. Cortical neuron generation is modulated by both intrinsic and extrinsic factors^2;3^. Intrinsic mechanisms include (epi)genetic factors that dictate the developmental potential of progenitors. Extrinsic factors encompass signaling molecules from neighboring cells, postmitotic neurons in the cortical plate (CP), embryonic cerebrospinal fluid secreted by the choroid plexus, and incoming thalamic afferents^5–9^. These signals influence the timed self-renewal and neurogenic potential of cortical progenitors. Additionally, invading cortical inhibitory interneurons (cINs) have been proposed to influence EOMES^+^ IPCs, thereby impacting the proper formation of cortical layers^10^.

Despite composing only 10-20% of the neuronal population, cINs, including Parvalbumin (PV) and somatostatin (SST) interneuron subtypes, are pivotal for precise information processing by inhibiting both excitatory and other inhibitory neurons^11^. Disturbing their development impairs the balance between excitation and inhibition, leading to abnormal cortical activity and cognitive impairments associated with various neurological and neuropsychiatric disorders, including schizophrenia and epilepsy^12–15^. The different types of cINs originate in three subpallial structures: the medial ganglionic eminence (MGE), the caudal ganglionic eminence (CGE), and the preoptic area (POA)^11^. The MGE gives rise to PV^+^ and SST^+^ interneurons, whereas the CGE generates Serotonin 3A receptor (5HTR3a) subtypes^16,17^. The POA produces mostly neurogliaform cells and Neuropeptide Y (NPY) expressing multipolar interneurons^18,19^. Postmitotic cINs migrate tangentially through the basal telencephalon into the cortex^13^. They invade the developing cortex along the marginal zone (MZ) and SVZ/intermediate zone (IZ), where they spread tangentially before switching to radial migration to enter the CP. This migration is guided by a diverse array of intrinsic and extrinsic cues^13,20–23^. For example, the transcription factors ARX and LHX6, expressed in MGE-derived cINs, directly and indirectly regulate the expression of cytokine receptors that mediate the attraction to CXCL12, which is expressed in the MZ migratory route of the developing cortex^24,25^. After tangentially spreading over the cortical areas, cINs invade the cortical plate. The cortical plate expresses NRG2, which exerts attractive effects on ERBB4-expressing interneurons^26^.

Epigenetic mechanisms, including histone modifications and DNA methylation, modulate intrinsic transcriptional programs^15,27^. It was previously reported that the DNA methyltransferase 1 (DNMT1) regulates gene expression in post-mitotic POA-derived cINs, promoting their proper migration and survival through non-canonical interactions with histone modifications^28^. Post-mitotic MGE-derived cINs, which give rise to SST^+^ and PV^+^ interneuron subtypes, also express DNMT1^28^. However, DNMT1 function in these subsets of immature, migrating cINs is unknown as of yet.

SST-expressing cells constitute about 30% of all cINs and play vital roles in inhibitory control, network synchronization, circuit plasticity, and cognitive functions^29^. SST^+^ cINs encompass two main subtypes: Martinotti cells, which migrate along the marginal zone (MZ) during development and form axon collaterals in layer I, and non-Martinotti cells, which disperse via the SVZ/IZ and have different layer distributions as well as targeting properties^13,29^.

The distinct migratory paths of SST^+^ interneurons are regulated by ARX and LHX6, as well as MAFB^24,30,31^. Specifically, MAFB marks Martinotti cells and, along with ARX and LHX6, directs their migration along the MZ^24,30,31^. This migration is closely associated with the subsequent formation of axonal arbors in layer I^31^.

Recognizing the functional relevance of SST^+^ cINs in cortical circuitry and the necessity of their precise development for cortical function, we investigated the role of DNMT1 in this process. Addressing this gap in knowledge advances the understanding of cell type-specific epigenetic regulation in cortical network formation.

Our findings show that reduced *Dnmt1* expression in SST^+^ cINs disrupts the transcriptional and DNA methylation profiles of key genes for cIN development, such as *Arx*, altering their migration along the MZ. This misregulation impacts excitatory cortical progenitor dynamics, affecting neurogenesis, and cortical layering, ultimately leading to functional and behavioral abnormalities in adulthood.

## Results

### Molecular dynamics simulations propose DNMT1’s catalytic domain binding to unmethylated DNA

DNMT1 is described as a maintenance methyltransferase, preserving DNA methylation during replication. This view is largely based on the comparison between a cryo-EM structure, showing a binding of DNMT1’s catalytic domain to the hemimethylated oligo 5′-ApCpTpTpApCpGpGpApApGpGp-3’ (HDNA)^32^, and the X-ray structure of the unmethylated oligo 5’- TpCpCpCpGpTpGpApGpCpCpTpCpCpGpCpApGpGp-3’ (UMDNA), which reveals weaker binding^33^. The latter observation is in apparent contrast to the reported dissociation constant (Kd) of the DNMT1/UMDNA complex in aqueous solution in the nM range^34^, comparable to that of DNMT1/HDNA and other high-affinity DNA/protein complexes (**Extended Data Table E1**). Together with recent findings reporting DNMT1 expression and functions in postmitotic and even adult neurons, such as POA and MGE-derived cortical GABAergic interneurons^28,35–37^, this proposes DNMT1 to have enzymatic functions in neurons other than the methylation of HDNA.

Visual inspection of the X-ray structure^33^ suggests that the weak interactions between the UMDNA and the catalytic domain of DNMT1 are caused by additional intermolecular contacts of UMDNA with two surrounding DNMT1 molecules in the crystal lattice (**Fig. 1a, Extended Figure E1a-f**). Consistently, in three independent 500-ns-long molecular dynamics (MD) simulations in solution, in conditions similar to those used for the K_d_ measurements^34^ (details are provided in **Extended data Tables 1-5**, and the **Supplementary methods**), the two moieties undergo a significant rearrangement at the end of the dynamics. Almost all of the interactions of the UMDNA’s 5’ region are formed with the catalytic domain, which performs the enzymatic reaction (**Fig. 1b-g, Extended Data Fig. E1g and h**). A CpG moiety interacts with Arg1234 and Asn1236 (**Fig. 1c**), as seen also in the cryo-EM structure of the HDNA/DNMT1 complex^32^ (PDB ID: 7XI9, **Extended Data Table E2**). The number of contacts between the protein and UMDNA in solution, which lacks the additional DNMT1 molecules that are present in the crystal (**Fig. 1a, b**), increased about 60% compared to the crystal structure (**Fig. 1d-g, Extended Data Fig. E1g, h**). Bioinformatic analysis showed that the solution MD structure aligns with other DNA/protein complexes with affinities in the nM range: the number of contacts for A^2^ of contact surfaces is 0.17, which is comparable to other complexes ranging from 0.17 to 0.21 (**Extended Data Table E1**). Thus, DNMT1 binds to UMDNA in solution stronger than in the crystal phase because of packing forces, consistent with K_d_ measurements in solution^34^.

**Figure 1.**
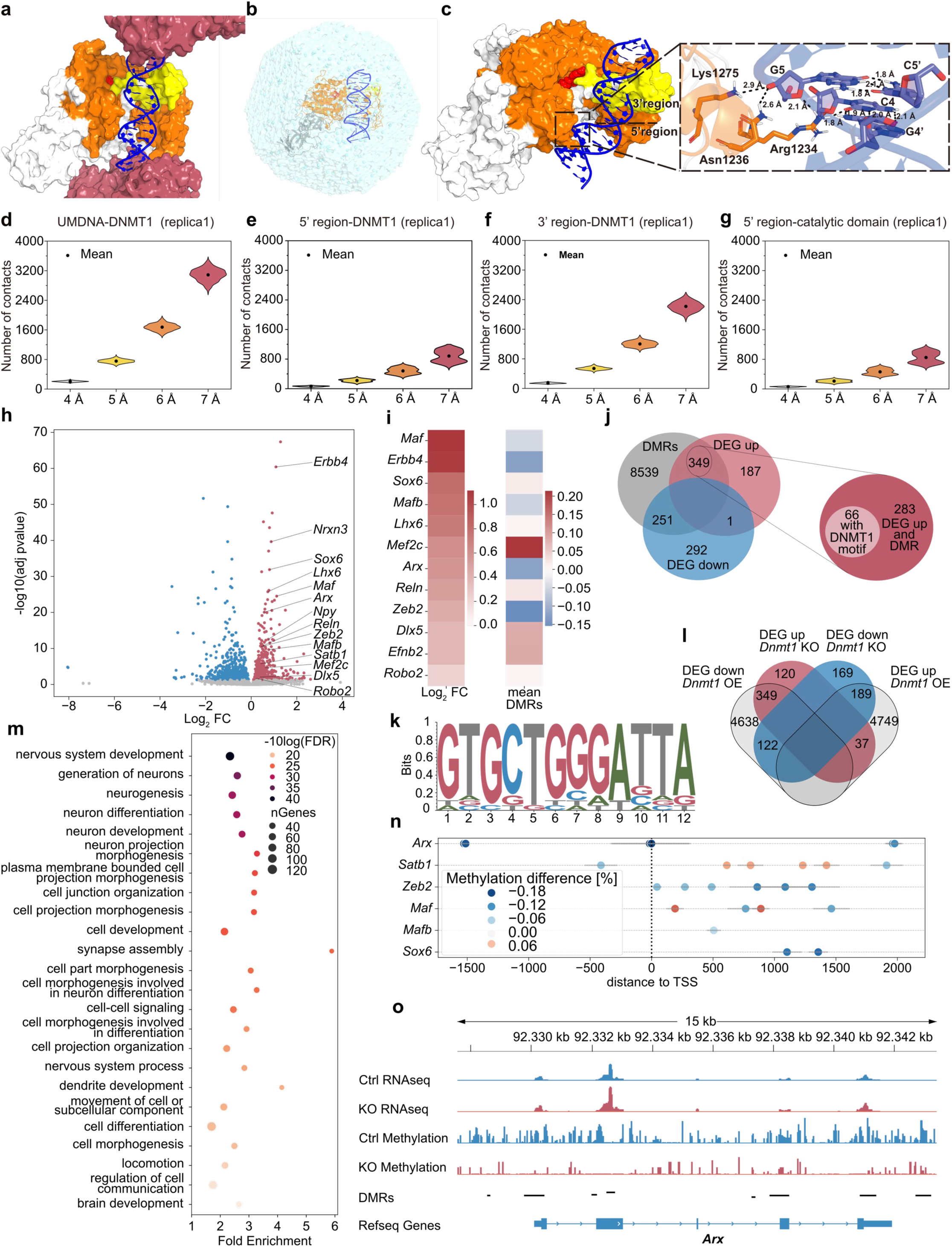
– DNMT1 regulates the expression of hub genes of cIN development in migrating E14.5 SST^+^ interneurons. **(a-g)** Molecular dynamic simulations of DNMT1 binding to unmethylated DNA (UMDNA). **(a)** Inter-unit DNMT1 proteins interacting with the DNMT1/UMDNA/SAH complex unit in the X-ray structure (PDB ID: 3PTA). Intra-unit DNMT1 regions shown include the autoinhibitory linker BAH1/BAH2 (white surface), catalytic domain (orange), SAH (red spheres), and CXXC domain (yellow), which interacts most with UMDNA (blue). The neighboring inter-unit DNMT1 proteins are illustrated in rose-pink. The inter-unit DNMT1 regions the UMDNA interacts with are mostly positively charged (**Extended Data Fig. E1b**). **(b-g)** Simulations of DNMT1/UMDNA interactions in aqueous solution. **(b)** Illustration of the DNMT1/UMDNA complex in solution. **(c)** The complex at the end of one of our three MD simulations (same coloring scheme as in **(a)**), with residues at the CpG site interacting with the catalytic domain highlighted by sticks. Only hydrogen atoms bound to polar groups are shown (as black dash lines). **(d-g)** Number of contacts between UMDNA **(d)**, the 5’ **(e)** and 3’ regions **(f)** with DNMT1, as well as between the 5’ region and the catalytic domain of DNMT1 **(g)** the during last-100-ns simulation (being consistent across replicas (**Extended Fig. E1g, h**). **(h-o)** Differential gene expression and methylation analysis of *Sst-Cre/tdTomato* control (ctrl) and *Sst-Cre/tdTomato/Dnmt1 loxP^2^* (*knockout;* KO) mice based on total RNA and enzymatic methyl-sequencing of FAC-sorted *tdTomato^+^* cells isolated from the E14.5 basal telencephalon. For RNA- and methyl-sequencing, we separately processed two samples per genotype, each consisting of cells pooled from multiple embryos (RNA sequencing: Ctrl *n* = 23 embryos; KO *n* = 14 embryos; methyl-sequencing: Ctrl *n* = 15 embryos; KO n = 11 embryos). **(h)** Volcano plot depicting the differentially expressed genes (DEGs) between control and KO samples. Genes annotated as significantly changed (adjusted *p*-value < 0.05) are colored with red depicting increased expression in KO. **(i)** Heatmap of the 50 genes with the lowest adjusted *p*-values, color-coded based on relative expression. **(j)** Venn diagram illustrating the overlap between up- or downregulated genes and genes associated with or containing a differentially methylated region (DMR). The intersection of upregulated and differentially methylated genes after knockout was tested for enrichment of the DNMT1 binding motif identified by DNMT1-ChIP-sequencing, shown in k. **(l)** DEGs identified in *Dnmt1* KO samples were compared to DEGs determined in embryonic stem cell-derived murine neurons overexpressing DNMT1. **(m)** Selection of brain development-related gene ontology (GO) terms enriched in genes that were both upregulated and differentially methylated in E14.5 *Sst-Cre/tdTomato/Dnmt1* KO cells. **(k)** DNA motif enriched in DNMT1-interacting chromatin, detected by ChIP-sequencing (*N* = 2 biological replicates). **(n)** Position of differentially methylated regions relative to the transcription start sites of genes related to cIN development. Horizontal bars indicate the region’s size, and color-coded is the mean methylation change for the respective region. **(o)** RNA sequencing tracks combined with the methylation profile of the *Arx* gene locus obtained from ctrl and KO samples.

Our analysis supports DNMT1’s broader functional role beyond maintenance methylation including potential DNA methylation-dependent regulation in postmitotic, non-replicating neurons, if the MD simulations reproduce the *in vivo* conditions.

### DNMT1 regulates key genes governing cIN development in embryonic SST⁺ cells

To examine the role of DNMT1 in SST^+^ cIN development, we generated a conditional knockout (KO) mouse model, in which *Dnmt1* deletion is induced by *Cre*-expression in postmitotic SST^+^ cells (*Sst-Cre/tdTomato/Dnmt1 loxP^2^*). *Sst-Cre/tdTomato* mice served as controls (**Extended Data Fig. E2a**). The *Sst-Cre* deleter strain is a well-established model, with *Cre*-dependent tdTomato expression being evident as cells leave the MGE (**Fig. 2b, c**), consistent with other studies^38–40^. Validation of the mouse models is shown in **Extended Data Fig. E2b-n**.

**Figure 2:**
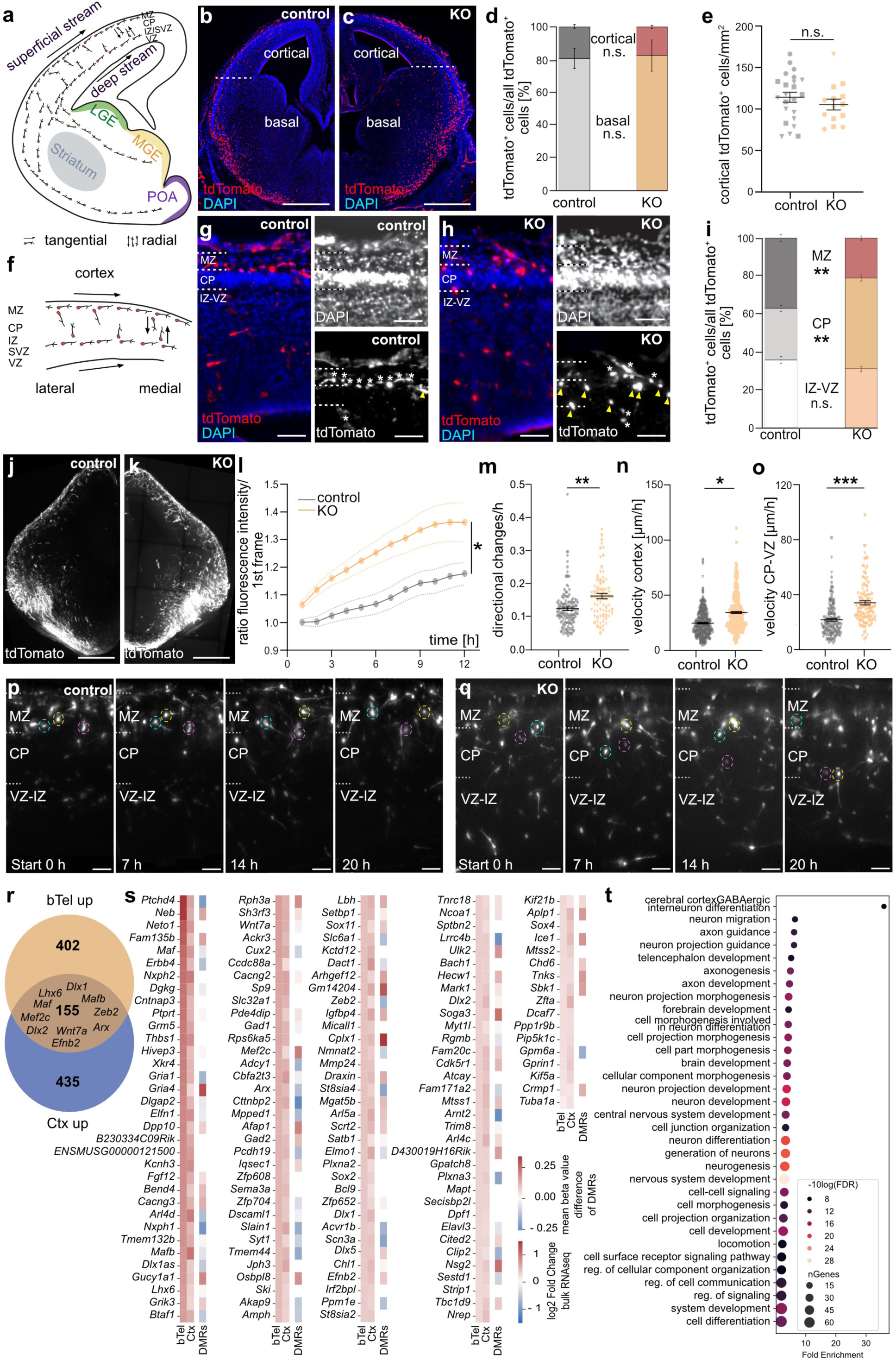
DNMT1 regulates the migration of SST^+^ interneurons in the developing cortex. **(a)** Schematic illustration of a coronally sectioned E14.5 brain hemisphere, depicting the migration routes of cINs. **(b, c)** Exemplary microphotographs of tdTomato^+^ interneurons in coronally sectioned (50 µm) hemispheres of *Som-Cre/tdTomato* (control) **(b)** and of *Som-Cre/tdTomato/Dnmt1 loxP^2^* (KO) embryos **(c)**. tdTomato-positive cells are labeled in red and DAPI in blue. Scale bars: 500 µm. **(d)** Quantification of the proportional distribution of tdTomato^+^ cells in the basal telencephalon and the cortical wall. Analyses were conducted in E14.5 coronal sections using a nested two-way ANOVA with *n =* 12 sections for control, *n =* 8 sections for KO, and *N =* 3 embryos per genotype. **(e)** Quantification of tdTomato^+^ cell number within the cortex of control and KO embryos normalized to the area, analyzed in 50 µm coronal cryosections (nested two-way ANOVA with *n =* 14 sections for control, *n =* 8 sections for KO, and *N =* 3 embryos per genotype). **(f)** Schematic illustration of migration trajectories within the E14.5 cortex. **(g, h)** Magnified microphotographs of the cortices of control and KO embryos (E14.5) to illustrate the distribution of tdTomato^+^ cells within the cortical zones. White asterisks point to cells migrating along the MZ (superficial migratory stream) or deep stream (SVZ/IZ). Yellow arrows indicate tdTomato^+^ cells located in the CP. Scale bars: 100 µm. **(i)** Quantitative analysis of the proportional distribution of tdTomato^+^ cells within the cortical zones normalized to the overall tdTomato^+^ cell count within the cortex of control and KO embryos at E14.5 analyzed in 50 µm coronal cryosections (nested two-way ANOVA; *n =* 14 sections for control, *n =* 8 sections for KO, and *N =* 3 embryos per genotype). **(j, k)** Temporal z-projections of migration tracks of tdTomato^+^ cells in organotypic brain slice cultures (350 µm thickness, coronal) from E14.5 *Som-Cre/tdTomato* (control) and *Som-Cre/tdTomato/Dnmt1 loxP^2^* (KO) embryos, captured over 20 h by live-cell imaging. Scale bars: 500 µm. **(l-o)** Live-cell imaging quantifications. Quantitative analysis of the increase in fluorescence intensity in the CP reflecting the invasion of tdTomato^+^ cells (first 12 h, nested two-way ANOVA with *n =* 4 slices for control, *n =* 6 slices for KO embryos, and *N =* 3 embryos per genotype) is shown in **(l)**, the average number of directional changes (*n =* 134 cells across 5 slices for control (*N =* 4 embryos), *n =* 74 cells across 3 slices for KO (*N =* 3 embryos), nested two-way ANOVA) is depicted in **(m)**. The velocity of all cells migrating within the cortex (*n =* 377 cells across 6 slices for control (*N =* 5 embryos), *n =* 323 cells across 5 slices for KO (*N =* 3 embryos)) is shown in **(n),** while the speed of cells within the area between CP and VZ is illustrated in **(o)** for control (*n =* 193 cells across 6 slices, *N =* 5 embryos) and KO (*n =* 114 cells across 3 slices, *N =* 3 embryos); nested two-way ANOVA. **(p, q)** Selected frames extracted from live-cell imaging recordings of organotypic brain slices (350 µm) derived from E14.5 control and KO embryos. Depicted are tdTomato^+^ cells migrating through the cortex. For each genotype, selected cells (dotted circles) are highlighted. Scale bars: 50 µm. **(r-t)** Differential gene expression analysis of FAC-sorted *Sst-Cre/tdTomato* and *Sst-Cre/tdTomato/Dnmt1 loxP^2^* cells isolated from the E14.5 cortex. Three samples per genotype were separately processed, consisting of cells pooled from multiple embryos (Ctrl *n* = 7 embryos; KO *n* = 8 embryos). **(r)** Venn diagram illustrating the overlap between upregulated genes in *Sst-Cre/tdTomato/Dnmt1 loxP^2^* cells from the basal telencephalon and the cortex (E14.5). **(s)** Heatmap illustrating the log2-fold change for the genes which were upregulated in *Sst-Cre/tdTomato/Dnmt1 loxP^2^* cells from the basal telencephalon and the cortex. Moreover, changes in DNA methylation (from the basal telencephalon dataset) are depicted. **(t)** Gene ontology (GO) terms enriched in genes that were upregulated in E14.5 *Sst-Cre/tdTomato/Dnmt1* KO cells from both the basal telencephalon and the cortex. Error bars represent the standard error of the mean (SEM). *p* < 0.05*\**, *p* < 0.01*\*\**, *p* < 0.001***; n.s.: not significant. CP: cortical plate, VZ: ventricular zone, SVZ: subventricular zone, MZ: marginal zone, IZ: intermediate zone, l: lateral, m: medial, Ctx: cortex, bTel: basal telencephalon. FC: fold change. Data points with the same symbol depicted in (e) and (l-n) belong to one embryo.

We focused on embryonic day (E) 14.5, a critical stage characterized by extensive SST^+^ cIN migration through the basal telencephalon and the onset of tangential migration within the cerebral cortex^41^. To identify putative DNMT1 target genes, we conducted RNA sequencing and methyl-sequencing of FACS-enriched E14.5 *Sst-Cre/tdTomato/Dnmt1 loxP^2^* (*Dnmt1* KO) and *Sst-Cre/tdTomato* (control) cells from the basal telencephalon. We revealed 1,160 differentially expressed genes (DEGs) and 13,080 differentially methylated regions (DMRs) overlapping annotated genomic regions (**Fig. 1h-j, Supplementary Table 1, 2**). Correlating DMRs with the DEGs, we identified 600 genes exhibiting both transcriptional alterations and DNA methylation changes in *Dnmt1* KO cells (**Fig. 1j, Supplementary Table 1-3**). Consistent with previous findings^28,36^, we observed up- or downregulated genes, as well as DMRs showing gain and loss in DNA methylation upon *Dnmt1* deletion. This aligns with findings from others, demonstrating that reduced *Dnmt1* and *Dnmt3a* expression leads to both gene activation and repression^38,42^. Downregulated genes are often interpreted as indirect effects, for instance, resulting from *Dnmt1* deletion-induced upregulation of a transcriptional repressor or adaptive transcriptional responses to physiological changes^36,38,42^. Given the partially redundant functions of DNMT1 and DNMT3A^43^, compensatory DNMT3A activity may contribute to the increased DNA methylation levels observed after *Dnmt1* deletion (**Supplementary Table 2**).

To identify potential targets of DNMT1-dependent repressive DNA methylation, we focused on the 349 of the 600 genes, which were upregulated in *Dnmt1* KO samples and displayed DMRs (**Fig. 1j**). Notably, 66 of these 349 genes contained DNMT1 binding motifs that we identified through additional ChIP-sequencing experiments (**Fig. 1j, k, Supplementary Table 4**).

Gene ontology (GO) enrichment analysis of upregulated genes with altered DNA methylation in *Dnmt1* KO cells revealed an overrepresentation of genes associated with nervous system development, neuron differentiation, cell-cell signaling, locomotion, neurogenesis, and cell differentiation (**Fig. 1m, Extended Data Fig. E3a; Supplementary Table 5)**. A similar enrichment was observed for the entire set of significantly upregulated genes in *Dnmt1* KO cells, regardless of concurrent DNA methylation changes (**Extended Data Fig. E3d, Supplementary Table 6**). Notably, many genes upregulated in E14.5 *Dnmt1* KO samples were downregulated in embryonic stem cell (ESC)-derived *Dnmt1^tet/tet^* neurons overexpressing DNMT1^44^, highlighting DNMT1’s repressive role in neurodevelopmental gene regulation (**Fig. 1l, Supplementary Table 7, 8**). Moreover, DNMT1-regulated genes in SST^+^ cINs exhibited significantly longer coding sequences, transcript lengths, genome spans, and extended 3′ and 5′ UTRs compared to the entire set of genes detected in SST⁺ cINs (*Chi-squared test*; **Extended Data Fig. E3b, c, e, f and E4b, c**). This aligns with the observation that *Mecp2* deletion, a reader of DNA methylation^45^, similarly leads to the upregulation of long genes expressed during neurodevelopment^46^.

Among DEGs with increased expression and DMRs in *Dnmt1* KO cINs, we identified genes coding for key cortical interneuron differentiation and maturation regulators, including *Arx*, *Zeb2*, *Mef2c*, *Dlx5*, *Sox6*, *Maf*, *Mafb*, and *Lhx6* (**Fig. 1h, i**). In addition to these well-characterized transcription factors governing cIN fate and migration^13,41,47^, DNMT1 also regulated genes impacting locomotion, and cell-cell signaling (**Fig. 1h, i**). These included *Erbb4*, *Reln*, *Robo2*, and members of the Eph/ephrin family such as *Efnb2*, all of which encode proteins critical for directional guidance and neuronal positioning^26,48,49,50^.

From these upregulated genes, *Erbb4*, *Arx*, *Zeb2*, *Maf*, and *Mafb* exhibited reduced DNA methylation, with *Erbb4*, *Arx*, and *Zeb2* showing the most pronounced reduction in DNA methylation averaged across all DMRs per gene (**Fig. 1i**). Of note, *Dnmt1* deletion in E14.5 SST^+^ cINs resulted in DMRs near transcriptional start sites (TSS) as well as within intragenic regions and intergenic loci (**Supplementary Table 2**). Given that promoter DNA methylation typically correlates with transcriptional repression^51^, we further examined methylation changes at the TSS of key cIN genes dysregulated in both *Dnmt1* KO and in *Dnmt1* overexpressing neurons (**Fig. 1n**; **Supplementary Table 7**). For *Arx*, *Zeb2*, *Mafb*, and *Sox6* reduced DMRs appeared at or near TSSs (**Fig. 1n, o**), aligning with their elevated expression in E14.5 *Dnmt1* KO cells (**Fig. 1h, i**). *Erbb4* exhibited intragenic DMRs more distant from the TSS (**Supplementary Table 2**). Intragenic DNA methylation has diverse regulatory functions, but is particularly implicated in neuronal differentiation and migration^52,53^. Consistently, many dysregulated genes in E14.5 *Dnmt1* KO samples encode proteins essential for cIN development and migration, such as ARX^24^, ZEB2^54,55^ and ERBB4^26^. In sum, our findings suggest that DNMT1 establishes proper gene regulatory networks essential for proper migration and differentiation of cortical SST^+^ interneurons.

### *Dnmt1* deletion in postmitotic SST^+^ interneurons causes a premature exit from the superficial migratory stream in the embryonic cortex

To investigate the potential impact of DNMT1 on SST^+^ cIN migration, we next analyzed brain sections of *Sst-Cre/tdTomato/Dnmt1 loxP^2^* (*Dnmt1* KO) and *Sst-Cre/tdTomato* (control) embryos. We initially focused on E14.5, when first SST^+^ interneurons have reached the cerebral cortex (**Fig. 2a-c**). Cortical dimensions were analyzed to confirm that both genotypes were at the same developmental stage (**Extended Data Fig. E5a, b**). *In situ* examination of *tdTomato*-expressing cells in E14.5 coronal brain sections revealed overall comparable numbers of SST^+^ cINs in the cortex and the basal telencephalon in both genotypes (**Fig. 2d, e**), indicating that the migration to the cortex was not impaired. However, within the cortex, we observed a reduced proportion of *tdTomato* cells migrating along the MZ and a significant increase of cells within the cortical plate (CP) in *Dnmt1* KO brain sections (**Fig. 2f-i**). Live-cell imaging of organotypic brain slices at E14.5 for 20 hours confirmed an elevated fraction of *Dnmt1* KO cells that deviated from the superficial migratory stream and entered the CP (**Fig. 2j-q, Supplementary Movies 1 and 2**). Notably, *Dnmt1* KO cells displayed a significantly increased frequency of directional changes compared to control *Sst-Cre/tdTomato* cells (**Fig. 2m**), suggesting a disruption in their ability to remain confined along the superficial migratory stream within the MZ. Moreover, their migratory pace was enhanced, which was particularly evident for radially migrating cells, and cells migrating through the CP (**Fig. 2n, o; Extended Data Fig. E6a**). This aligns with an increased path length seen for the migrating *Dnmt1* KO cells in living brain slices (**Extended Data Fig. E6b**). Of note, in contrast to *Dnmt1*-deficient POA-derived *Hmx3*-expressing interneurons^28^, we did not observe morphological abnormalities for migrating *Sst-Cre/tdTomato/Dnmt1 loxP^2^* cells (**Extended Data Fig. E6c**). Consistently, *in vitro* experiments aiming to assess the morphology of *Dnmt1* siRNA-treated MGE cells (E14.5 + 1DIV) did not reveal detectable morphological differences compared to control conditions (**Extended Data Fig. E6d-g**).

Given the impaired migration within the cerebral cortex, we profiled the transcriptional changes underlying the premature exit of SST^+^ interneurons from the marginal zone and their increased invasion of the CP in E14.5 *Dnmt1* KO embryos. To this end, we conducted RNA sequencing on FACS-enriched *Sst-Cre/tdTomato* cells isolated from the E14.5 cortex of both genotypes (**Fig. 2r-t, Supplementary Table 9**). Notably, we obtained a prominent overlap with the upregulated genes identified in *Dnmt1* KO cells from the basal telencephalon, with master regulators of GABAergic interneuron differentiation and migration such as *Dlx1*, *Dlx2*, *Dlx5*, *Arx*, *Lhx6*, *Maf*, *Mafb*, *Mef2c*, *Satb1*, and *Zeb2* being altered in expression in both datasets (**Fig. 2r, s, t**). Of note, *Mef2c,* known to be regulated by *Maf* and *Mafb,* is a key factor in specifying PV^+^ cINs^56^. Similarly, *Dlx5* is implicated in PV^+^ cIN development^57^.

Also, genes related to cell-cell signaling appeared significantly overrepresented among the commonly upregulated genes in *Dnmt1* KO cells from both compartments (**Fig. 2t, Supplementary Table 9**). This included *Erbb4* (**Fig. 2r, s, Extended Data Fig. E6h, i**), known to facilitate cIN invasion into the cortical plate upon activation by its ligand Neuregulin 3 (NRG3)^26^. Its increased expression could thus mediate the premature exit of *Dnmt1* KO cells from the superficial migratory stream.

Additionally, *Efnb2* expression was persistently elevated in *Dnmt1* KO cells upon entering the cortex. Since EPHB1 is strongly expressed in the cortical MZ^20^ and acts as a repulsive cue for migrating neurons from the basal telencephalon via reverse signaling^58^, the increased *Efnb2* levels could promote further the premature exit from the MZ.

Together, these findings emphasize that DNMT1 is critical for regulating the proper development and migration of SST^+^ cINs within the cortex at E14.5, acting through control of key genes involved in cIN migration and differentiation.

Although *Dnmt1*-deficient SST^+^ interneurons prematurely invaded the CP by E14.5 within 20 hours (**Fig. 2j-l**), we did not observe significant differences at later developmental stages (E16.5 and E18.5). This was neither the case for the proportion of cells that reached the cortex nor for the absolute number, density, or distribution of *Dnmt1* KO cells within the cortical regions quantified (**Extended Data Fig. E7**). The cortical dimensions were likewise similar between knockout and wildtype embryos at E16.5 and E18.5 (**Extended Data Fig. E5c-f**), confirming that the embryos analyzed were of comparable developmental stages. This suggests that the impact of *Dnmt1* deletion on the directional migration of SST^+^ cINs within the MZ may be transient.

### DNMT1-dependent regulation of SST^+^ interneuron migration affects cortical progenitors non-cell-autonomously

The development of cINs and cortical excitatory neurons is intricately linked^10,59^, and supernumerary cINs in the IZ elicit altered IPC numbers impacting the generation of upper-layer excitatory neurons^47^. Observing an impaired migration pattern of SST^+^ cINs prematurely invading the CP and proliferative zones prompted us to investigate its influence on excitatory cortical progenitors and neurogenesis. Moreover, we found changed expression of genes involved in cell-cell signaling, known to modulate cortical neurogenesis, such as *EfnB2*^60^, in *Dnmt1* KO cells. EFNB2 binds to EPHA4^61^, which is expressed in RGCs^5^. Activation of EPHA4 receptors by EFNA5, imported by invading thalamic afferents, has been shown to regulate RGC division, impacting IPC generation and cortical layer formation^5^.

To investigate whether the altered migration pattern and/or expression of signaling molecules in *Dnmt1* KO embryos affects the generation of cortical excitatory neurons, we performed immunostaining for TBR1, labeling early-born neurons of the preplate and layer VI^62,63^. We found a significant increase in TBR1^+^ cell numbers within the CP in E14.5 *Dnmt1* KO brains (**Fig. 3a-c**), which became even more evident at E16.5 (**Fig. 3f-h**). At E18.5 the radial extension of both layer VI (TBR1^+^ cells) and the remaining CP were significantly expanded in the conditional *Dnmt1* KO embryos compared to age-matched controls (**Fig. 3k-m, p and q**). We next investigated potential changes in EOMES^+^ IPCs in *Dnmt1* KO embryos, which represent an intermediate progenitor stage from RGCs to neurons, proposed to give rise to most cortical projection neurons of all cortical layers^64;65,66^, and had already been reported to be impacted by cINs^10^. Consistently, we observed an increase in EOMES^+^ IPCs at E14.5 (**Fig. 3d, e**) and E16.5 (**Fig. 3i, j**) in *Dnmt1*-deficient embryos, coinciding with the timing of the premature exit of *Dnmt1*-deficient SST^+^ interneurons from the MZ at E14.5 and the elevated numbers of TBR1^+^ neurons. As no differences in the pool of IPCs were detected at E18.5 (**Fig. 3n, o**), the deletion of *Dnmt1* therefore seems to elicit a temporally restricted non-cell autonomous effect on cortical IPCs and the production of deep-layer neurons.

**Figure 3:**
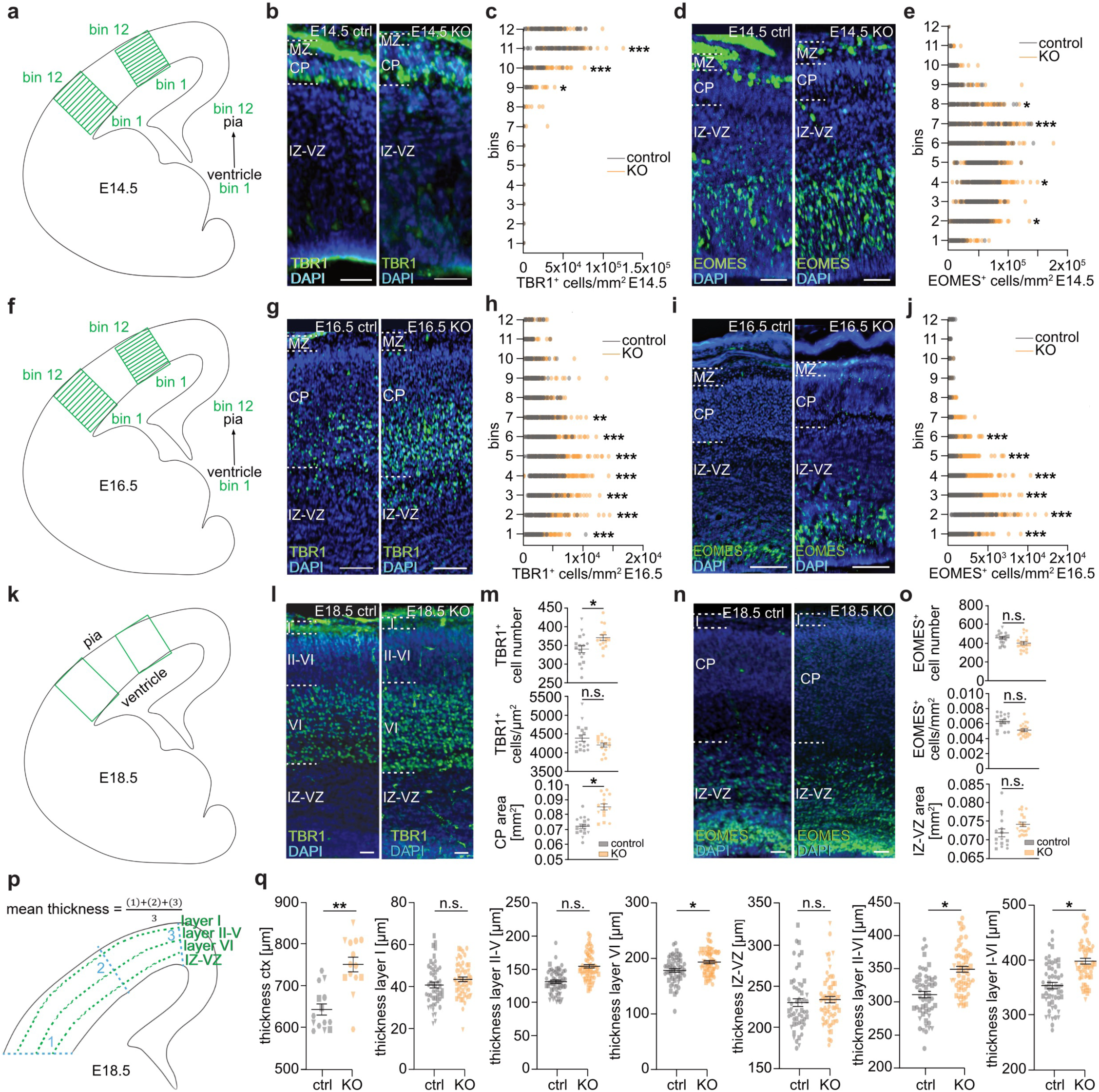
Conditional deletion of *Dnmt1* in SST interneurons affects IPCs and the generation of TBR1^+^ cells non-cell autonomously. **(a, f)** Schematic illustrations of a coronally sectioned embryonic brain hemisphere, depicting the lateral and dorsal localization of the two bin selections, used to quantify the density and distribution of EOMES^+^- and TBR1^+^- cells in E14.5 and E16.5 brain sections. **(b-e)** TBR1 and EOMES immunostaining in coronally sectioned (50 µm) brains of E14.5 *Sst-Cre/tdTomato* (control) and *Sst-Cre/tdTomato/Dnmt1 loxP^2^* (KO) embryos. **(b** and **d)** Exemplary microphotographs of (**b**) TBR1 immunostaining (green) and **(d)** EOMES immunostaining (green); scale bars: 50 µm (DAPI is shown in blue). **(c** and **e)** Quantitative analysis of the TBR1^+^ postmitotic neuron numbers **(c)** and EOMES^+^ intermediate progenitor cell (IPC) numbers **(e)** per bin in E14.5 cortices of control and KO embryos, analyzed in 50 µm coronal cryosections (two-way ANOVA with subsequent Bonferroni correction*, n =* 13 control and *n =* 9 KO sections from *N =* 3 embryos for both genotypes). **(g)** TBR1 immunostaining (green) in coronally sectioned (50 µm) brains of E16.5 control and KO embryos, scale bars: 50 µm. DAPI is depicted in blue. **(h)** Quantitative analysis of the TBR1^+^ postmitotic neuron numbers per bin in the E16.5 cortex of control and KO embryos, analyzed in 50 µm coronal cryosections (two-way ANOVA with subsequent Bonferroni correction*, n =* 8 for control and KO from *N =* 3 embryos for both genotypes). **(i)** EOMES immunostaining (green) in coronally sectioned (50 µm) brains of E16.5 control and KO embryos, scale bars: 50 µm. DAPI is depicted in blue. **(j)** Quantitative analysis of the EOMES^+^ postmitotic neuron numbers per bin in the E16.5 cortex of control and KO embryos, analyzed in 50 µm coronal cryosections (two-way ANOVA with subsequent Bonferroni correction*, n =* 15 control and *n*= 13 KO sections from *N =* 3 embryos for both genotypes). **(k)** Schematic illustration of a coronally sectioned E18.5 embryonic brain hemisphere, depicting the lateral and dorsal localizations used to quantify the density and distribution of EOMES^+^- and TBR1^+^-cells. **(l** and **n)** Exemplary microphotographs of (**l**) EOMES immunostaining (green) and **(n)** TBR1 immunostaining (green) in coronal sections of E18.5 control and KO brains; scale bars: 50 µm (DAPI is shown in blue). **(m** and **o)** Quantitative analyses of the total numbers and densities of EOMES^+^IPCs **(m)** and TBR1^+^postmitotic neurons **(o)** in the E18.5 cortex of control and KO embryos, analyzed in 50 µm coronal cryosections. Additionally, the area of the IZ-VZ and the CP was analyzed in **(m)** and **(o)**, respectively (nested two-way ANOVA with *n =* 11 sections for both genotypes from *N =* 4 embryos, respectively). **(p)** Schematic illustration of the cortex of a coronally sectioned embryonic brain (E18.5), depicting the parameters analyzed for the quantification shown in (**q**). Three regions (lateral (1), dorsolateral (2), and dorsal (3)) were averaged to obtain the mean thickness respectively for each layer (nested two-way ANOVA with *n =* 12 control and *n =* 9 KO sections. *N =* 4 brains for both genotypes). *p* < 0.05 *, *p* < 0.01 **, *p* < 0.001 ***. MZ: marginal zone, CP: cortical plate, IZ-VZ: intermediate zone-ventricular zone, ctrl.: control, n.s.: not significant. Data points depicted in (m), (o), and (p) with the same symbol arise from the same embryo.

### DNMT1 function in SST^+^ interneurons modulates cortical progenitor and neurogenesis

The non-cell-autonomous effects of *Dnmt1* deletion in SST^+^ cINs on cortical progenitors and the increased generation of deep-layer neurons suggest that SST^+^ interneurons provide signaling cues influencing progenitor dynamics, similar to thalamic afferents importing EFNA5^5^. Notably, altered expression of signaling molecules known to modulate cortical neurogenesis and progenitor division, including *Wnt7a* and *EfnB2* (**Figure 2s, t, Supplementary Table 1, 9**)^6, 60,69^ were detected in *Dnmt1* KO cells.

To further investigate potential cIN-progenitor interactions, we employed CellChat analysis of scRNA-sequencing data from E14.5 dorsal telencephalon, predicting ligand-receptor interactions between SST+ cINs and cortical progenitor types: RGCs and IPCs (**Fig. 4a, b, Extended Data Fig. E8a-c**).

**Figure 4:**
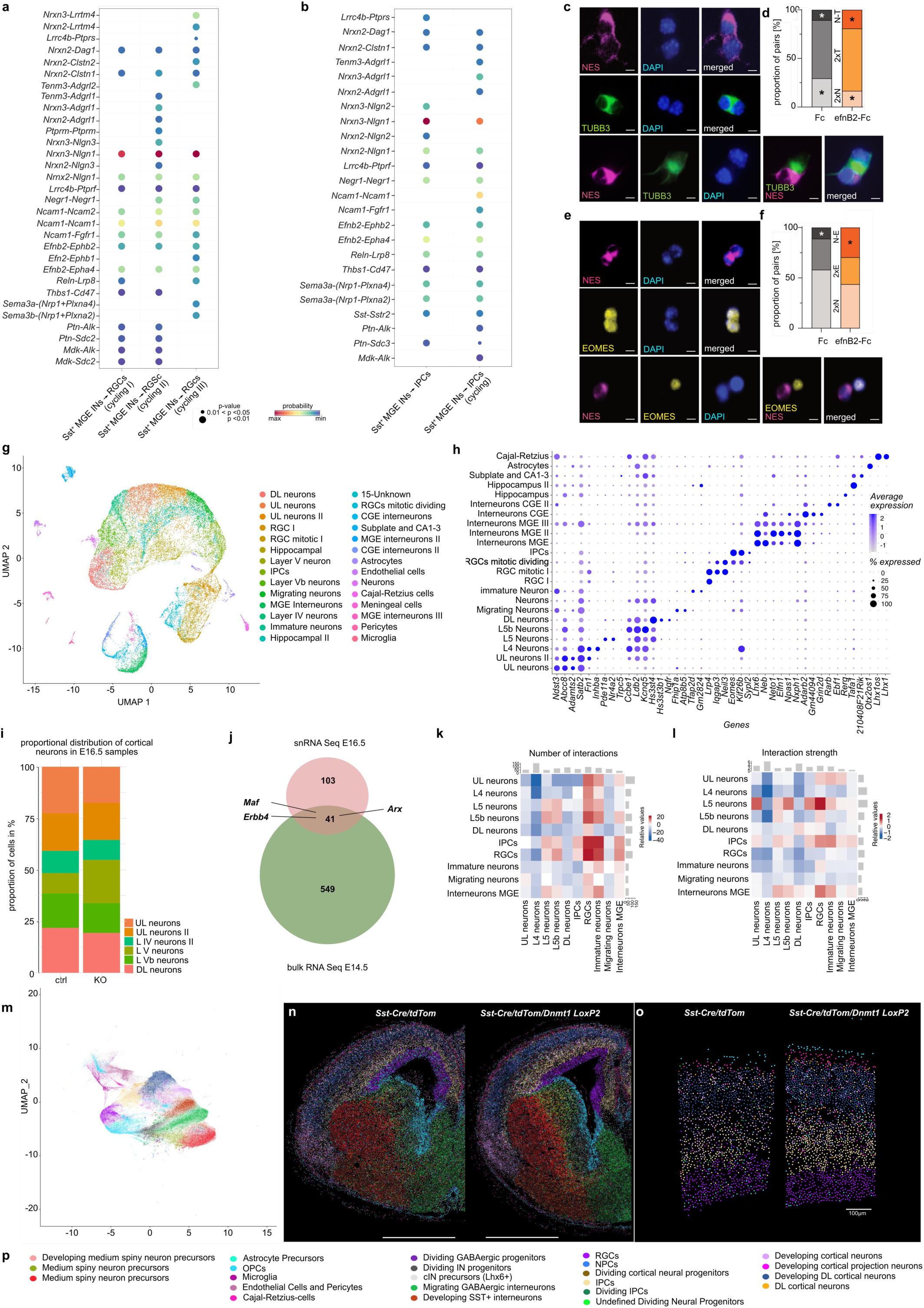
Single-cell resolution approaches reveal non-cell-autonomous effects of SST+ interneurons on cortical progenitors. **a, b)** Single-cell RNA sequencing (scRNA-seq) of dorsal telencephalons from E14.5 C57BL/6J mice identifies significant ligand-receptor interactions mediating communication from SST+ cINs to **(a)** different apical cortical progenitor populations (APs) and **(b)** to intermediate progenitors (IPCs), filtered by differentially expressed genes (DEG). Communication probability is represented by dot color, and the corresponding *p*-value by dot size, with *p*-values being computed using a one-sided permutation test. **(c-f)** Pair cell assay performed with cortical neurons isolated from E14.5 C57BL6/J embryos, plated at clonal density. Cell clones were either treated with control (ctrl)-or recombinant efnB2-Fc for 24 h, before immunocytochemical staining for nestin (NES, magenta), β-III-tubulin (TUBB3, green), EOMES (yellow) and DAPI (blue). Scale bars: 5 µm. **(d** and **f)** Quantification of the proportion of nestin/nestin (2xN), β-III-tubulin/β-III-tubulin (2xT), nestin/β-III-tubulin (N-T), EOMES-EOMES (2xE), and nestin-EOMES (N-E) positive cell pairs normalized to the overall number of these pairs. Unpaired Welch’s *t*-test with control-Fc: *n* = 90 cell pairs, efnB2-Fc: *n* = 91 cell pairs, from *N =* 4 experiments in **d**; and control-Fc: *n* = 90 cell pairs, efnB2-Fc: *n* = 94 cell pairs, from *N =* 4 experiments in **f**. **(g-l)** Single nuclear RNA (snRNA) sequencing of cortical cells prepared from E16.5 *Sst-Cre/tdTomato* and *Sst-Cre/tdTomato/Dnmt1 loxP^2^* embryos (*n* = 2 brains from *N* =2 mice per genotype). **(g)** UMAP illustrating cell clusters determined by snRNA sequencing across all samples. **(h)** Dot plot depicting marker gene expression in the different cell clusters. **(i)** Bar plot showing the proportional distribution of the cell numbers determined for all postmitotic excitatory cortical neurons. **(j)** Venn diagram illustrating the overlay of the upregulated genes in E14.5 cortical FAC-sorted *Sst-Cre/tdTomato/Dnmt1 loxP^2^* cells (bulk RNA sequencing) compared to *Sst-Cre/tdTomato* samples, with the upregulated genes found in E16.5 cortical *Sst^+^/Gad^+^* cells form *Sst-Cre/tdTomato/Dnmt1 loxP^2^* compared to *Sst-Cre/tdTomato* samples (snRNA sequencing). **(j, k, l)** Interaction maps extracted with CellChat from the snRNA sequencing dataset of E16.5 cortices from *Sst-Cre/tdTomato/Dnmt1 loxP^2^* and *Sst-Cre/tdTomato* embryos, depicting number of interactions **(k)** and interaction strength **(l)**. **(m-p)** Multiplexed spatial transcriptomics analysis of E16.5 *Sst-Cre/tdTomato/Dnmt1 loxP^2^* and *Sst-Cre/tdTomato* brain sections using MERFISH. **(m)** UMAP visualization of distinct cell clusters identified based on gene expression profiles. **(n)** Representative tissue sections with spatially resolved cell clusters mapped onto anatomical regions, scale bar 1000 µm. **(o)** Magnified sections from the cortex Scale bar = 100 µm. **(p)** Cluster annotation for (m-o). *p* < 0.05 *, *p* < 0.01 \*\**, p* < 0.001 ***. n.s.: not significant.

NRXN3-NLGN1 signaling had the highest probability for interactions between SST^+^ cINs with both RGCs and IPCs. However, *Nrxn3* was not detected among the upregulated genes in *Sst-Cre/tdTomato/Dnmt1 loxP^2^* cINs at E14.5 (**Supplementary Table 9**). In addition, EFNB2-EPHA4 signaling was revealed as a potential mediator of these interactions, with *EfnB2* expressing SST^+^ MGE-interneurons interaction with RGCs as well as IPCs at E14.5 (**Fig. 4a, b**). E16.5 scRNA-sequencing data from the dorsal telencephalon retrieved similar results (**Extended Data Fig. 8d, e**). Supporting this, *Efnb2* was upregulated in *Dnmt1*-deficient SST^+^ cINs and altered in *Dnmt1*-overexpressing neurons (**Fig. 1h**, **Fig. 2s, t; Supplementary Table 1, 7, 9**). Given its known role in promoting neurogenesis at mid-corticogenesis^60^ and its known binding to EPHA4^69^ being expressed by RGCs^5^, we hypothesized that increased *Efnb2* expression in *Dnmt1*-deficient SST^+^ cINs contributes to enhanced RGC differentiation into IPCs and deep-layer neurons. Pair-cell assays confirmed that ephrinB2-Fc stimulation promotes neurogenic RGC divisions and IPC generation (**Fig. 4c-f**).

To examine altered cIN-progenitor interactions in *Dnmt1* KO embryos, we performed single-nucleus (sn) RNA-seq on E16.5 cortical cells from both genotypes. This approach allowed sample freezing for simultaneous processing and captured potential differences in cIN-progenitor interactions but also non-cell-autonomous effects on postmitotic neuron generation, which might become more pronounced at this developmental stage.

Aligning with the histological data, *Dnmt1* KO samples at E16.5 showed an increased proportion of deep-layer cortical neurons and a reduced fraction of upper-layer neurons (**Fig. 4g-I; Extended Data Fig. 8f, h, i**). Additionally, *Erbb4*, *Arx*, and *Maf* - key regulators of cIN development - along with numerous other genes previously identified as upregulated through bulk RNA sequencing of E14.5 cortical *Dnmt1* KO cells, displayed significantly elevated expression in *Sst*^+^/*Gad2^+^* cells from the E16.5 *Dnmt1* KO samples (**Extended Data Fig. 8g**). Applying *CellChat* analysis, genotype-related differences in interactions between MGE interneurons (senders) and RGCs were revealed, while this was not evident for the interaction between MGE interneurons and IPCs (numbers and interaction strength; **Fig. 4k, l**). This supports the instructive role of cINs on RGCs, and that this influence was increased in *Dnmt1* KO embryos. We further conducted MERFISH analysis in E16.5 brain sections of *Sst-Cre/tdTomato/Dnmt1 loxP^2^* and *Sst-Cre/tdTomato* embryos. Here, we also confirmed the elevated proportion of deep-layer neurons in *Dnmt1* KO brains and an increase in IPCs (**Fig. 4 m, n, o; Extended Data Figure 9)**.

In summary, DNMT1 is crucial for migration and function of SST^+^ cINs, with its loss disrupting migration trajectories, altering signaling, and promoting RGC differentiation into IPCs and deep-layer neurons.

### *Dnmt1* deletion-induced changes at embryonic stages manifest in an altered cortical architecture at adult stages

Finally, we investigated whether the *Dnmt1* deficiency-related embryonic defects results in long-lasting structural changes in the adult cortex. We conducted a comparative analysis of the distribution, density, and fate of *Sst-Cre/tdTomato* control and *Dnmt1* KO interneurons in cortices of adult mice. Immunohistochemistry confirmed SST expression *Sst-Cre/tdTomato* cells, as well as the reported co-expression of NPY and CALB2 (Calretinin)^70^ in both genotypes (**Extended Data Fig. E10a-e**). However, the proportion of SST^+^ cINs co-expressing PV^71^ was significantly increased in *Dnmt1* KO mice compared to controls (**Extended Data Fig. E10e-j**). This is in line with the elevated transcription of *Dlx5* and *Mef2c* found in E14.5 *Dnmt1* KO cells (**Fig 1h, i**; **Fig. 2t**), both coding for drivers of PV^+^ cINs development^57,72,73^.

*Sst-Cre/tdTomato* cIN numbers were comparable across all layers between genotypes (**Fig. 5a-c**), but cell density in the superficial layers was significantly increased in *Dnmt1* KO mice (**Fig. 5c)**. This correlated with a reduced radial extension of CUX1-positive layers II-IV and an expansion of deep layers V-VI, while overall cortical thickness remained unchanged (**Fig. 5a, b, d).**

**Figure 5:**
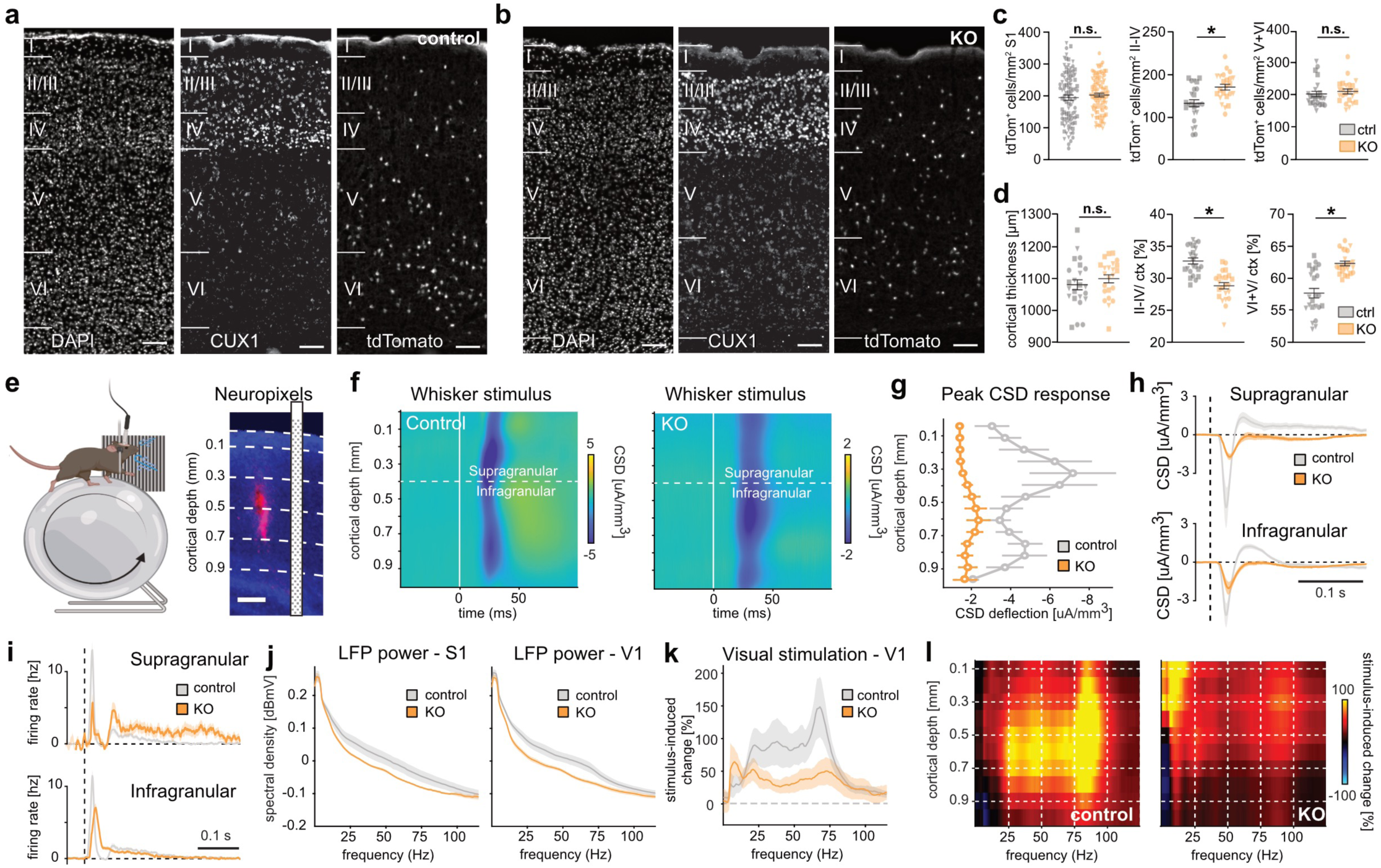
Adult *Dnmt1*-deficient mice show aberrant cortical architecture and functionality. **(a-d)** *Dnmt1* KO mice display changes in cortical layer thickness alongside alterations in *Sst-Cre/tdTomato* cell density. **(a, b)** Exemplary microphotographs of immunostainings of CUX1 and tdTomato combined with DAPI taken from S1 (sagittal slices of Bregma 1.32 and 1.44, six-month-old male mice). The CUX1^+^ supragranular layers II-IV can be distinguished from infragranular layers V and VI. Scale bars: 100 µm. **(c, d)** Quantification of the tdTomato^+^ cell density normalized to the given area of the respective layers (**c**), the cortical thickness, and the proportional thickness of the deep and upper layers (**d**). Nested two-way ANOVA with *n =* 24 slices for control and *n =* 23 for KO from *N =* 3 brains for both genotypes. *p* < 0.05 *, *p* < 0.01 **, *p* < 0.001 ***, n.s.: not significant. Data points depicted in (c and d) with the same symbol arise from the same mouse. (e) Left: Neuropixels recordings in S1 of head-fixed control and *Dnmt1* KO mice, running on a wheel. Right: Example brain slice from S1, showing red dye fluorescence from the probe. The scheme visualizes the probe position. **(f)** Average current source density (CSD) across the whole cortical depth in S1, following a 20 ms tactile stimulus (white line) to the distal whisker pad. Blue colors indicate excitatory and yellow colors inhibitory currents. The earliest responses in control mice (left) occurred in the granular layer at ∼400 µm cortical depth and then spread to layer II/III (100-300 µm) and layer V (500-900 µm). In contrast, responses were much weaker and less spatially defined in KO mice. Matching the anatomical results, response differences to controls were largest in the supragranular layers. **(g)** Quantification of CSD response peaks across sessions for each group. Supragranular responses above 400 µm were significantly lower and delayed in *Dnmt1* KO versus control mice (peak amplitude*_control_* = -4.74 ± 1.22 µA/mm^3^, peak amplitude*_KO_* = -1.41 ± 0.18 µA/mm^3^, *p* = 0. 0037; peak time*_control_* = 27.20 ± 0.84 ms, peak time*_KO_* = 31.80 ± 1.03 ms, *p* = 0.0041; *n* = 8 sessions from 2 mice per group). **(h)** CSD traces in S1, averaged over tactile responses in either supragranular (top) or infragranular layers (bottom). Clearly visible are reduced and delayed responses in supragranular layers for *Dnmt1* KO mice (orange). Same data as in panels (f) and (g). **(i)** Peristimulus time histogram (PSTH) for the spiking response of S1 neurons in supra- or infragranular layers to tactile stimulation in KO (orange) and control mice (gray). **(j)** Spectral power density (PSD) of LFP recordings across all layers in either S1 (left) or primary visual cortex (V1, right). In both cortical areas, PSDs fall off with frequency but remain significantly higher in the high-frequency gamma range from 30-90 Hz for control versus KO mice (S1: gamma power*_control_* = -23.69 ± 21.32 dBmV, gamma power*_KO_* = -61.95 ± 3.19 dBmV, *p* = 0.0378; V1: gamma power*_control_* = -25.65 ± 8.65 dBmV, gamma power*_KO_* = -68.27 ± 4.38 dBmV, *p* = 0.0037; *n* = 8 sessions from 2 mice per group). **(k)** Stimulus-induced change in PSD in V1, computed as the percent difference relative to baseline. Full-field visual stimulation induced a strong increase in V1 gamma power (> 30 Hz), especially around 70 Hz, in control mice. In contrast, stimulus-induced modulation was much weaker in KO mice (relative gamma*_control_* = 104.25 ± 24.42%, relative gamma*_KO_* = 28.71 ± 6.31%, *p* = 0.0018; *n* = 8 sessions from 2 mice per group). **(l)** Stimulus-induced change in PSD gamma power across cortical depth. Changes in control mice were largest between 0.3- and 0.7-mm but largely absent in KO mice. Significance for panels g, j, and k was based on a Wilcoxon *rank-sum* test, values are mean ± SEM. Shading in panels h, i, and k show the SEM.

The shift in layer proportions aligns with increased TBR1^+^ neurons and IPCs at E14.5 and E16.5, as well as snRNA-seq and MERFISH findings (Fig. 3, 4). These results indicate that *Dnmt1* deletion in SST^+^ interneurons non-cell autonomously alters cortical progenitor dynamics, impacting adult cortical architecture and SST^+^ cIN densities in superficial layers.

### *Sst-Cre/tdTomato/Dnmt1 loxP^2^* mice display functional abnormalities

To determine whether the observed structural alterations in the adult cortex also translate into functional changes, we conducted electrophysiological recordings in the primary somatosensory barrel cortex (S1) of awake head-fixed *Dnmt1* KO and control mice on a running wheel (**Fig. 5e left**, **Extended Data Fig. E11**). To simultaneously capture population and single-cell neural activity across the entire cortical depth, we recorded with high-density Neuropixels probes (**Fig. 5e right; Extended Data Fig. E11a, b**) while mice were passively stimulated with short air puffs to their whisker pad^74^. Tactile stimulation induced a clear transient response in average local field potentials (LFPs) in S1 of control mice, whereas LFP responses in conditional *Dnmt1* KO mice were much weaker and temporally delayed (**Extended Data Fig. E11c**). To dissect the spatiotemporal structure of neural responses across all cortical layers, we employed a current source density (CSD) analysis based on our LFP recordings^75^. Consistent with the structural alterations in *Dnmt1* KO cortices, we found clear differences in responses across layers between the genotypes. As expected from the literature^76^, the earliest responses for control mice (**Fig. 5f left**) occurred in the granular layer (∼400 *μ*m cortical depth), then spreading to layer II/III (100-300 *μ*m) and layer V (500-900 µm), which indicates intact spatiotemporal processing of sensory signals. In contrast, the CSD profile of *Dnmt1* KO mice was severely disrupted, with tactile responses being largely confined to the deeper infragranular layers and only weak responses in the superficial layers (**Fig. 5f right**). Quantification of CSD peak responses across the entire depth confirmed that the most pronounced differences between *Dnmt1* KO and control mice occurred in the upper cortical layers (**Fig. 5g, h**). Notably, while control mice exhibited a positive rebound following the initial negative CSD deflection, the responses in *Dnmt1* KO mice lacked this feature, possibly due to disrupted feed-forward inhibition (**Fig. 5h**).

To quantify the tactile responses of cortical S1 neurons, we also used spike-sorting to isolate the spiking activity of sorted clusters. Consistent with our earlier results, the average spiking activity of S1 neurons showed that tactile responses were much weaker in *Dnmt1* KO mice, especially in the superficial layers (**Fig. 5i**, **Extended Data Fig. E11d**). Moreover, spiking responses of upper-layer neurons were much longer lasting compared to controls, with neural activity extending up to 400 ms after stimulation. This suggests reduced temporal precision in sensory processing in *Dnmt1* KO mice, potentially due to impaired feedback inhibition.

We also investigated whether functional alterations manifest at the level of cortical network oscillations. Gamma (γ) oscillations from 30-120 Hz, which strongly rely on the accurate function of SST^+^ cINs^77,78^, are essential for integrating neural networks within and across brain structures during cognitive processes, and abnormalities are a feature of cognitive diseases such as schizophrenia, Alzheimer’s disease, and Fragile X syndrome^79^. Thus, changes in γ-oscillations represent a useful marker of function and dysfunction in cortical circuit operations^80^. Indeed, the high-frequency LFP power in S1 was significantly reduced in *Dnmt1* KO versus control animals (**Fig. 5j left**).

Similar disruptions in oscillatory network dynamics were observed in the primary visual cortex (V1; **Fig. 5j right**). Visual stimulation (5-s-long visual grating stimuli) reliably induced γ-oscillations in V1, with control mice showing a clear 70 Hz peak, consistent with stimulus-induced γ activity (Fig. 5k, l). This effect was markedly reduced in *Dnmt1* KO mice, which instead exhibited increased low-frequency activity in superficial cortical layers (**Fig. 5k, l)**. These oscillatory changes suggest that *Dnmt1* deletion disrupts neural networks, likely due to structural alterations in cortical architecture, particularly in superficial layers.

To distinguish whether these functional deficits stem from altered cortical structure or impaired SST^+^ cIN function, we optogenetically activated SST^+^ cINs and assessed their spontaneous firing, action potential waveform, and inhibitory capacity **(Extended Data Fig. E11d-g)**. No significant differences emerged between control and *Dnmt1* KO neurons, indicating that SST^+^ cIN functionality remained intact. Thus, the observed cortical network disruptions likely result from structural changes, such as superficial layer thinning and increased SST^+^ cIN density, rather than a generalized impairment of SST^+^ cIN function.

### *Sst-Cre/tdTomato/Dnmt1* KO mice display behavioral abnormalities

To investigate whether functional impairments in *Dnmt1* KO mice translated into impaired behavior we tested the animals’ sensory perception. Therefore, we employed a visuo-tactile evidence accumulation task and trained head-fixed *Dnmt1* KO and control mice to detect tactile and visual stimuli. Both genotypes learned the stimulus detection at similar rates, performed both uni- and multisensory task conditions with equal behavioral performances and showed multisensory enhancement (**Extended Data Fig. E12a-d**), indicating intact sensory perception in *Dnmt1*-deficient mice. Moreover, we tested learning and memory capabilities using the Morris water maze, which also showed no differences between genotypes (**Extended Data Fig. E12e-g**).

Notably, we observed abnormal eye movements in head-fixed *Dnmt1* KO mice during the tests on the running wheel, characterized by brief episodes of significantly dilated pupils (more than double their normal size) lasting 30-60 seconds, accompanied by facial spasms or salivation (**Fig. 6a**). These episodes of pupil dilation correlated with a marked increase in low-frequency activity (1–4 Hz), indicating epileptiform activity in KO mice (**Fig. 6b**). Further analysis of the propagation of this activity across cortical depth revealed that these events were particularly prominent in the superficial layers (**Fig. 6c**). In line with this, we detected prolonged seizure durations for *Dnmt1* KO ² mice in response to administration of PTZ (pentylenetetrazol), which is widely utilized to examine epileptic events in animal models and to assess changes in seizure responses^81^ (**Figure 6d-f**, **Supplementary Videos 3-9**). The latencies to the first occurrence of various epileptic events were comparable between genotypes (**Extended Data Fig. 12h, i**). However, *Dnmt1*-deficient mice exhibited higher Racine scale scores^82^, indicating increased seizure severity (**Figure 6d-f)**. This aligns with studies reporting that alterations in SST^+^ cIN activity and function can lead to epileptic events^83^ and γ oscillations have also been proposed to play a role in seizure generation^84^.

**Figure 6:**
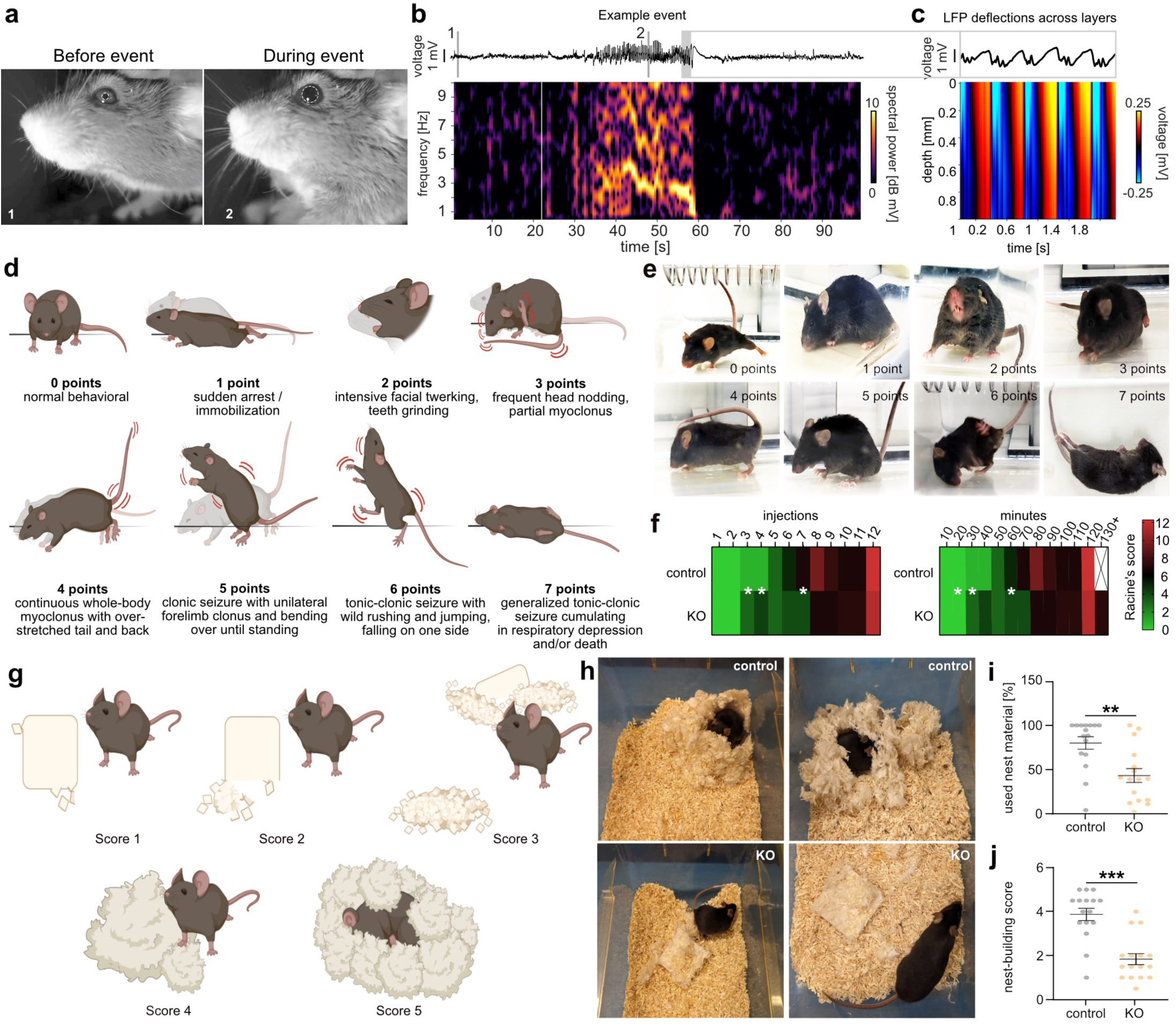
Adult *Dnmt1*-deficient mice show behavioral abnormalities. **(a)** Example images of a head-fixed *Dnmt1* KO mouse before and during an epileptiform event with dilated pupil and postural changes (white circles). **(b)** Example trace and spectrogram from cortical LFP recording during the event show a strong increase in theta oscillations for ∼30 seconds. Times of video images 1 and 2 (white numbers in (a)) are shown as gray lines in the top trace. **(c)** Quantification of LFP deflections across depth for a 2-second time window during the event. **(d-f)** Pentylenetetrazol (PTZ)-injections every 10 min in three months old *Sst-Cre/tdTomato* (control) and *Sst-Cre/tdTomato/Dnmt1 loxP^2^* males (*Dnmt1* KO) point to defects in cortical inhibition in KO mice. **(d)** Schematic illustrations defining PTZ-induced seizures using Racine’s scoring: 1: immobilization by sudden arrest and lying flat on the ground for several seconds up to minutes. 2: facial twerking, intensive mouth movement, and/or teeth grinding without convulsions. 3: frequent head nodding and/or partial myoclonic events (mostly unilateral limb- or tail clonus). 4: continuous whole-body myoclonus indicated by a stretched back, as well as extensive and dynamic stiffening of the tail with overhead pointing towards the rostrum. 5: clonic seizure characterized by unilateral forelimb clonus and bending over until reaching a sitting/standing position. 6: clonic-tonic seizure with falling on one side, wild rushing, and jumping. 7: Generalized tonic-clonic seizure is indicated by wild rushing, jumping, and tonic extension cumulating in overstretched limbs and tail. This epileptiform event results in respiratory depression and/or death (end of the experiment). (**e**) Footages captured from video monitoring of PTZ-induced convulsions for respective Racine’s scores, that are depicted in (d). **(f)** Heat maps and quantification of Racine’s scale events associated with the given number of PTZ-injections or passed time. Within every column (illustrating the number of applied injections (left) or minutes after the first injection of PTZ (right)), all detected epileptiform events were correlated to their respective severity score (0-7), added up, and averaged for each genotype (rows). *Dnmt1*-KO mice show significantly higher Racine’s scale points (=severity levels) indicated by red colors. Wilcoxon *rank-sum* test, *p* < 0.05 *, *p* < 0.01 **, *p* < 0.001 ***. *N* = 10 animals per genotype. **(g-j)** Nest building test reveals a significant deterioration in using given material and forming proper nests in adult *Sst-Cre/tdTomato/Dnmt1* loxP^2^ (KO) animals compared to *Sst-Cre/tdTomato* (control) mice. With the help of the scoring index depicted in (**g**) and formerly described^94^, all mice were ranked based on nest quality and the amount of used material (**h-j**). Representative photographs are shown in (**h**) for control and KO mice. Tests were performed overnight and twice within one week. Respective values were averaged for both trials (**i, j**). Unpaired, two-tailed Student’s *t*-test and additional unpaired Welch’s *t*-test. *N* = 16 male adult mice for both genotypes.

Moreover, we observed frequent repetitive motor behavior of *Dnmt1* KO mice in their home cages (**Supplementary Video 10**). Such behavioral abnormalities are often linked to neurological and neuropsychiatric diseases^85^. Notably, seizures frequently co-occur with comorbid symptoms, including autism-like stereotypies (e.g., repetitive motor behavior) and mood disorder-related phenotypes^86^. Thus, we next assessed nest-building performance, as this test assessed behavior associated with neuropsychiatric disorders^87^. In line with our functional results, we found that *Dnmt1* KO mice showed a reduced performance in nest-building and diminished interest in the provided nest-building material (**Fig. 6g-j**).

In summary, our cellular, functional, and behavioral results demonstrate that *Dnmt1* deletion in postmitotic SST^+^ interneurons leads to altered cortical architecture and network function in adult mice, manifesting in disease-related behavioral abnormalities. This strongly suggests that DNMT1 plays a crucial role in the development of SST^+^ interneurons and has non-cell autonomous functions essential for the proper formation of the cerebral cortex.

## Discussion

Two novel and main features of cortical development arise from our experiments: (i) dispersion of SST^+^ interneurons along the superficial migratory stream within the cortical MZ is regulated by DNMT1, and (ii) invading cINs influence RGCs and thereby proper cortical layering.

*Dnmt1* deletion in SST-expressing interneurons caused a premature exit from the MZ at E14.5, accompanied by increased numbers of EOMES^+^ IPCs at E14.5, and more TBR1-positive postmitotic neurons at E14.5 and E16.5, likely resulting from an altered division mode of RGCs. This led to an increased generation of deep-layer neurons and enlarged deep layers at adult stages, while upper layers were reduced in radial extension. These changes in cortical architecture together with altered densities of SST^+^ interneurons in adult mice were associated with prominent functional and behavioral deficits, demonstrating the importance of DNMT1 for establishing accurate network function in the adult cortex.

While DNMT1 is well known as a maintenance methyltransferase, primarily responsible for preserving DNA methylation during replication, emerging evidence—including our findings— suggests a broader role for DNMT1 in postmitotic neurons^35,36,43^. DNMT1 is expressed in non-dividing POA- and MGE-derived cINs, where it is proposed to regulate neuronal development through both canonical and non-canonical mechanisms^28,36^. Based on results in the crystal phase, Song et al^33^ have suggested weak binding of DNMT1’s catalytic domain to UMDNA. Solution-based affinity measurements challenge this hypothesis, demonstrating a high-affinity interaction for both HDNA/DNMT1 and UMDNA/DNMT1 complexes^34^, that is in similar scale compared to other DNA-protein complexes. Our MD simulations resolve this contradiction by showing that UMDNA undergoes a significant conformational rearrangement in solution, increasing the interactions of its CpG sites with DNMT1’s catalytic domain, as it was observed for the HDNA/DNMT1 complex^32^. Given that our findings in solution are valid also for *in vivo* conditions, they support the hypothesis that DNMT1 has a functional role beyond maintenance methylation. In line with this, DNMT1 has been shown to interact with CFP1 (CysxxCys finger protein 1), which presents a high affinity for unmethylated DNA^88^. Moreover, *de novo* methylation activity at certain repetitive elements and single copy sequences has already been shown for DNMT1^89^. Functional cooperation of DNMT1 during *de novo* methylation of DNA has further been described^90^, and gene-specific *de novo* methylation can be initiated by reintroduction of DNMT1 in cells lacking DNMT1 and DNMT3b, that present with nearly absent genomic methylation^91^. Taken together, these observations strongly propose DNMT1’s function beyond maintenance DNA methylation, likely contributing to (*de novo*) DNA methylation-dependent gene regulation in postmitotic neurons. This provides an additional layer of epigenetic control during neuronal differentiation and function. In combination with its described non-canonical actions through crosstalk with histone-modifying mechanisms^28,92^, our findings render DNMT1 as an active participant in epigenetic programming in postmitotic cINs and putatively also other neuronal subsets^35,36,43^.

*Dnmt1* deletion in SST^+^ cINs impairs their migration within the MZ. Martinotti cells, constituting the majority of SST^+^ interneurons^70^, migrate primarily along the MZ during corticogenesis^31^, while non-Martinotti SST^+^ cells disperse rather via the SVZ/IZ^31^. Given the selective effects of *Dnmt1* deletion on MZ migrating SST^+^ interneurons, DNMT1 seems to be relevant for the proper migration of Martinotti cells. Our global transcriptome and methylome analyses propose that DNMT1 controls essential key transcription factors of MGE cIN development such as *Lhx6*, *Arx*, *Zeb2*, *Satb1*, *Sox6*, and *Dlx5*^13,41,47^. LHX6, ARX, MAF, and MAFB are proposed to regulate the migration along the MZ, as *Lhx6*, *Maf*, *Mafb*, and *Arx* mutant mouse brains display defects in MZ migration of cINs^24,30,31^. The altered expression of these genes in *Sst-Cre/tdTomato/Dnmt1-loxP^2^* cells is in line with their premature exit from the MZ. For *Arx,* the most prominent reduction in DNA methylation at the TSS was observed (**Fig. 1i, n**), which is in line with its increased expression in *Sst-Cre/tdTomato/Dnmt1-loxP^2^* samples from the basal telencephalon and the cortex. DNA methylation at promoter regions is well known for its repressive power^51^. Thus, our data propose DNMT1 to keep proper *Arx* expression levels in migrating SST^+^ cINs, and through this regulate the migration along the MZ.

ARX has been reported to directly and indirectly modulate other genes controlling different stages of cIN development, including *Maf*, *Mafb*, *Mef2c*, *Nkx2.1*, *Lmo1*, *Nrg1*, *Erbb4*, *Npr1*, and *EphA4*^24^, many of which were altered in expression upon *Dnmt1* deletion in SST^+^ interneurons. For *Maf* and *Mafb*, we detected increased expression and reduced DNA methylation levels in *Dnmt1* KO samples (**Fig. 1h, I, n**), pointing to a DNMT1-dependent regulation as well. However, a transcriptional dysregulation of these and other ARX target genes such as *Mef2c* or *Erbb4* as a secondary effect of changed *Arx* expression cannot be excluded.

ERBB4 promotes the switch from tangential to radial migration and the distribution of interneurons within the CP, by interacting with NRG3 being expressed in the CP^26^. As *Nrg3* overexpression in the CP results in premature invasion of migrating interneurons into the CP^26^, the augmented *Erbb4* expression observed in *Sst-Cre/tdTomato/Dnmt1-loxP^2^* might cause the premature exit from the MZ. Moreover, increased *efnB2* expression in *Sst-Cre/tdTomato/Dnmt1-loxP^2^* cells could contribute to the precocious invasion of the CP. EPHB1 is expressed in the MZ^20^, and EPHB1-triggered reverse signaling has already been shown to elicit a repulsive response in migrating interneurons^58^. Of note, DNA methylation dependent regulation of *efnB2* expression was already proposed in neuronal stem cells^93^. Thus, DNMT1-mediated repression of *Erbb4* and *Efnb2* could regulate the proper migration of SST^+^ cINs along the superficial migratory stream and the timed invasion of the CP.

Apart from migration regulation, some of the dysregulated genes are involved in cell fate determination of cINs. For example, MEF2C and DLX5 drive the PV fate^57,72,73^, and both were increasingly expressed in E14.5 *Sst-Cre/tdTomato/Dnmt1-loxP^2^* cells. MEF2C initiates distinct transcriptional programs in immature PV^+^ interneurons upon settling within the cortex by opening the chromatin landscapes at PV-specific loci^72,73^. In addition to ARX, MAF, and MAFB, promote *Mef2c* expression^56^, which also increased in expression upon *Dnmt1* deletion. This aligns with the rise in PV-positive SST^+^ tdTomato cINs in adult *Dnmt1* KO mice. Thus, alongside migration regulation, DNMT1-dependent transcriptional remodeling seems to promote the SST^+^ cIN cell fate. Thus, although bulk RNA and methyl sequencing of FAC-sorted cells comes with limitations in matters of cell population heterogeneity, we obtained transcriptional changes that align with the observed phenotype of disturbed migration along the MZ, some of which were also confirmed by snRNA sequencing.

However, migration and final differentiation are interconnected processes. For example, the migration of Martinotti cells is functionally related to the morphological maturation and thus, the adaption of their final fate, as the proper formation of their axon collaterals in layer I depends on their migration along the MZ^31^. These axon collaterals extend horizontally for up to 2000 µm, making up approximately three-quarters (76%) of the GABAergic axons in this layer^70^. This enables Martinotti cells to influence the activity of numerous pyramidal neurons, which are their primary postsynaptic targets^70^. Hence, DNMT1 might also indirectly influence the proper formation of these powerful axon collaterals in layer I, by promoting the migration of SST^+^ cINs along the MZ. Disturbed formation of axon collaterals could contribute to the strong functional and behavioral impairments that we observed for adult *Sst-Cre/Dnmt1* KO mice, despite the overall number of SST^+^ cells not having been affected. This is consistent with data from *Mafb* KO mice, which also display a premature exit from the MZ and no changes in interneuron numbers and distribution in adults. Yet, functional impairments were likewise observed^31^.

The described functional deficits might also arise from the altered cortical architecture, which seem to arise from the non-cell autonomous effect of *Dnmt1* deletion SST^+^ cINs on cortical progenitors, resulting in altered SST^+^ cIN densities in the adult cortex, that link the effects on migration to the adult phenotype.

SST+ cINs regulate cortical excitability and synchronization^70^, for which their altered density in the superficial layers of *Dnmt1* KO mice likely contributes to observed functional and behavioral deficits. Consistently, functional impairments were most pronounced in these layers. LFP recordings showed weaker, delayed tactile responses with disrupted spatiotemporal processing, reduced activation, and prolonged spiking activity, particularly in the upper layers. Alongside diminished γ-oscillations, low-frequency activity was increased. Additionally, abnormal eye movements with severe pupil dilations correlated with heightened low-frequency activity (1–4 Hz) in superficial layers, reinforcing the link between structural changes and functional deficits.

As optogenetic stimulation revealed no intrinsic deficits in SST^+^ cIN firing properties upon *Dnmt1* deletion, our findings suggest that functional disruptions are primarily driven by structural cortical alterations. However, we cannot exclude additional DNMT1-dependent functional impairments in adult SST^+^ cINs, akin to DNMT1’s role in regulating GABAergic transmission in PV interneurons^36^. Future studies using inducible *Sst-Cre/tdTomato/Dnmt1 loxP^2^* mice to analyze methylation and gene expression signatures in adult SST^+^ cINs could clarify embryonic versus adult effects, though this is beyond the scope of the present study.

The expansion of deep cortical layers and reduction of upper layers observed in the adult cortex in adult *Sst-Cre/tdTomato/Dnmt1 loxP^2^* mice can be traced back to embryonic stages, where enhanced deep-layer neuron production and altered IPC numbers were detected. This strongly implies, that the migration defects in *Sst-Cre/tdTomato/Dnmt1 loxP^2^* embryos non-cell-autonomously affect cortical progenitors and hence the timed generation of cortical neurons.

As evidenced by Sessa et al. (2010)^59^, the development of cINs and excitatory neurons must have been effectively integrated throughout evolution, since an increase in the production of excitatory neurons also necessitates a rise in interneuron generation to maintain the proper balance of excitation and inhibition. Moreover, invading cINs may influence cortical progenitors, as they have been shown to modulate EOMES^+^ IPC proliferation, thereby affecting the timing of excitatory neuron generation^10^. Our findings extend this hypothesis by providing evidence that cINs also impact RGCs, which in turn regulate IPC and neuronal production.

Specifically, we implicate EFNB2-EPHA4 signaling in RGC division, similar to the reported modulation of EPHA4-expressing RGCs by thalamic afferents importing EFNA5^5^, which shifts apical progenitors between proliferative and neurogenic divisions. Both EFNA5 and EphA4 deficiency enhances deep-layer neuron production by altering RGC neurogenic potential^5^.

Similarly, cINs may provide external cues via membrane-bound or secreted signals. *Sst-Cre/tdTomato/Dnmt1-loxP^2^* interneurons showed increased *Efnb2* expression, encoding EFNB2, a regulator of cortical neurogenesis. While progenitor-restricted *Efnb2* deletion delays neurogenesis, EFNB2-driven EPHB signaling transiently boosts neuronal output^60^, mirroring our pair cell assay results. This ephrinB2-dependent neurogenic shift occurred within a specific temporal window of corticogenesis^60^, aligning with prior studies showing that temporally restricted neurogenic changes shape cortical layer thickness^5,93^.

In sum, the altered expression of secreted and membrane-bound signaling molecules with known potential to regulate cortical progenitor division and fate, in concert with the altered migration pattern of *Sst-Cre/tdTomato/Dnmt1 loxP^2^* interneurons could lead to the defects in cortical layer formation. This suggests an essential role for cINs in modulating the temporal production of distinct subtypes of excitatory cortical neurons, and a high level of crosstalk between immature cIN precursors and cortical progenitors to optimize the formation of neural circuits. These findings also establish a key role for DNMT1 in orchestrating cortical development through cell-autonomous and non-cell-autonomous actions.

## Methods

### Molecular Dynamics Simulations

The AMBER 22 software suite was employed. Details on the DNMT1/UMDNA/SAM complex and force field parameters are given in the **Extended data Figure E1 and Extended Data Table E3-E5**. Long-range electrostatic interactions were calculated using the Particle Mesh Ewald method^97^. Non-bonded interactions were treated with a 10 Å cutoff.

The system was energy-minimized *in vacuo* with 10,000 steps of steepest descent, followed by 10,000 steps of conjugate gradient minimization in order to eliminate atomic collision. The heavy-atom RMSD of the optimized configurations with respect to the initial geometry was 1.4 Å. Then, the complex was solvated in a truncated octahedral water box with a minimum distance of 12 Å from the solute to the box edge (**Extended Data Table E5**). K^+^, Na^+^, and Cl^−^ ions were added to neutralize the system. The concentration of K^+^ was similar to that in the nucleus (**Extended Data Table E5**). Periodic boundary conditions were applied. The whole system was then subjected to (i) 10,000 steps of steepest descent followed by 10,000 steps of conjugate gradient minimization with a 100 kcal/(mol·Å²) restraint on the whole solute. (ii) the same as (i), but with restraints only on heavy atoms of the solute; (iii) the same as (i), but without restraints. Successively, the system was heated from 0 K to 100 K over 5 ps then relaxed at 100 K for 5 ps, then from 100 K to 310 K in 0.5 ns, and finally relaxed at 310 K for 4.5 ns by using Langevin dynamics^98^, with a restraint of 100 kcal/(mol·Å²) was applied to the heavy atoms of the solute. The time integration step was set to 1 fs. Finally, three independent 500 ns isobaric-isothermal simulations with different initial velocities were performed without restraints and with a time step of 2 fs. Langevin dynamics^98^ and the Monte Carlo barostat^99^ were used to maintain a constant temperature (310 K) and pressure (1 atm), respectively. CPPTRAJ^100^ was used for trajectory analyses.

### Animals

In the recent study transgenic mouse strains with a genetic C57BL/6J background (initially obtained from the University Hospital UKA Aachen, Germany) were used. *Sst^tm2.l(cre)Zjh^*/J x *B6.CgGt(ROSA)26Sor^tml 4(CAG-tdTomato )Hze^* (*Sst^+/-^-Cre/tdTomato)* served as control animals whereas *Sst^tm2.l(cre)Zjh^/J* x B6.CgGt(ROSA)26Sor^tm14(CAG-tdTomato)Hze^ x *B6; 129S-Dnmt1tm2Jae*/J (*Sst-Cre^+/-^/tdTomato/Dnmt1 loxP^2^)* were used as *Dnmt1* knockout (KO) model. For more details see supplementary information.

### Isolation of Embryonic and Adult Brains

Individuals of embryonic stage 14.5, 16.5 or 18.5 were isolated as described in Symmank et al. (2018)^101^. Briefly, pregnant females were anesthetized by intraperitoneal administration of ketamine/xylazine (200/25 mg/kg living weight per injection up to a maximum application of 600/75 mg ketamine/xylazine per kg living weight in total). Upon reaching surgical tolerance, the abdominal cavity was opened to expose the uterine horns that were removed. Embryos were isolated from the uteri and decapitated. For histological analyses of E14.5 and E16.5 brains, the heads were directly transferred to 4% paraformaldehyde (PFA)/1x phosphate-buffered saline (PBS). For histological analyses at E18.5, brains were removed from the skull before transferring them to 4% PFA/1x PBS. E14.5 heads were fixed for 5 hours, while E16.5 heads and E18.5 brains were fixed overnight at 4 °C on an orbital shaker (50 rpm). Afterwards, stepwise cryopreservation was performed by overnight incubation first in 10% Sucrose/1x PBS, followed by 30% sucrose/1x PBS at 4 °C on an orbital shaker (50 rpm). Tissue was frozen in liquid nitrogen and stored at -80 °C.

For the isolation of adult brains, male mice were sacrificed with an overdose of 5% (v/v) isoflurane. For histological analyses and identification of Neuropixels recording sites, transcardial perfusion was conducted with 1x PBS (pH 7.4) followed by 4% PFA/1x PBS (pH 7.4) with a pump^102^. After brain preparation, post-fixation was conducted in 4% PFA/1x PBS for 24 h at 4°C on a roller mixer with constant rotation (approx. 50 rpm). Then, cryopreservation was performed in 10% sucrose/1x PBS and 30% sucrose/1x PBS for 24 h each at 4°C on a roller mixer with constant rotation (approx. 50 rpm) before brains were frozen in liquid nitrogen and stored at -80 °C.

### Organotypic brain slices and single-cell preparations

MGE- and cortical single cells were prepared as described previously^20,36^. Briefly, embryonic heads were collected and dissected in Gey’s Balanced Salt Solution (GBSS, pH 7.4)/0.65% D-glucose on ice. MGEs and cortices were cut out and collected in Hanks’ Balanced Salt Solution (HBSS, w/ phenol red, w/o calcium, w/o magnesium)/0.65% D-glucose on ice. The tissue was treated with 0.04% of trypsin/EDTA (Thermo Fisher Scientific, U.S.A.) for 17 min at 37°C, before MGE cells were dissociated in cold Dulbecco‘s Modified Eagle Medium (DMEM) with additional L-glutamine and 4.5 g/L D-glucose (Thermo Fisher Scientific, U.S.A.), 10% fetal bovine serum (FBS; Biowest, U.S.A.), and 1% penicillin/streptomycin (P/S) (Thermo Fisher Scientific, U.S.A.) by trituration with glass Pasteur pipettes and by subsequent filtering through a nylon gauze (pore size 140 µm; Merck, U.S.A.). Dissociated single cells were seeded on laminin (19 µg/ml; Sigma-Aldrich, U.S.A.)/poly-L-lysine-(10 µg/ml; Sigma-Aldrich, U.S.A.) coated glass coverslips. For MGE cells, we used a density of 455 cells/mm^2^. Cortical single cells were seeded in clonal densities of 150 cells/mm^2^. Transfection of MGE cells with siRNA oligos occurred after 5–6 h of incubation (37°C, 5% CO_2_ and 95% relative humidity) in Neurobasal medium with phenol red, 1x B27™, and 0.25x GlutaMAX (Gibco, U.S.A.) for 24 h. For this, the medium was supplemented with Lipofectamine^TM^ 3000 in accordance with the manufacturer’s instructions (Thermo Fisher, U.S.A.) together with *Dnmt1* siRNA (30 nM, #sc-35203, Santa Cruz, U.S.A.) and a scrambled control siRNA construct (15 nM, BLOCK-iT^TM^ Fluorescent Oligo, #2013, Thermo Fisher, U.S.A.). After 24 h MGE cells were fixated with 4% PFA/1x PBS for 10 min.

Incubation of cortical cells plated at clonal density occurred at 37°C, 5% CO_2_ and 95% relative humidity in culture medium (Neurobasal medium with phenol red, 1x B27™, 0.25x GlutaMAX (Gibco, U.S.A.), and 0.4% methylcellulose (Sigma, U.S.A.)) for 24 h. Treatment of cortical single cells was conducted 5–6 h after plating. For this, the medium was supplemented with 10 µM 5-ethynyl-2′-deoxyuridine (5-EdU, baseclick^TM^, baseclick GmbH, Germany) and with either 5 µg/mL of a recombinant human Fc control protein (Rockland Immunochemicals, U.S.A.) or 5 µg/mL of a recombinant human efnB2-Fc (R&D Systems, U.S.A.). Both recombinant proteins were pre-clustered with 10 µg/mL of an anti-human IgG antibody (Thermo Fisher, U.S.A.). Cortical single cells were fixated with 4% PFA/1x PBS for 10 min.

For organotypic brain slice preparations^28^, brains of E14.5 embryos were embedded in 4% low melt agarose (37°C) in Krebs buffer (126 mM NaCl, 2.5 mM KCl, 1.2 mM NaH^2^PO_4_, 1.2 mM MgCl_2_ * 6H_2_O, 2.1 mM CaCl_2_; pH 7.4; supplemented with 10 mM D-glucose and 12.5 mM NaHCO_3_, sterile-filtered). For sectioning of 350 µM coronal slices, a 5100mz-Plus vibrating microtome (speed: 0.7–0.8 mm/s and blade oscillation frequency: 5 Hz; Camden Instruments, United Kingdom) was used. Slices were transferred to ice-cold post-holding buffer (1x Krebs buffer, 10 mM HEPES, 1% P/S, 0.2% gentamycin, pH 7.4). Slices were imaged in µ-slide 4 Well imaging plates (ibidi GmbH, Germany) coated with 19 µg/mL laminin (Sigma-Aldrich, U.S.A.) and 10 µg/mL poly-L-lysine (Sigma-Aldrich, U.S.A.) in Neurobasal without phenol red, 1x B27™, 1% P/S, 0.5% D-glucose, 10 mM HEPES, using a confocal-like Leica DMi8 fluorescent microscope in combination with a THUNDER® imager unit (Leica, Germany) and the corresponding software *LASX* (Leica, Germany) equipped with an incubation chamber (37°C, 5% CO_2_ and 95% relative humidity). Imaging was conducted with a wavelength of 544 nm (TRITC-channel) in tile scans including z-stacks of 8–10 µm step size every 15 min. Post-processing of the respective footages was executed in the *LASX* software using the “*Mosaic Merge*”- and “*Thunder Lightning (Large Volume)*”-tools. Finally, a maximum intensity projection of the merged tile scans was processed using the *Fiji* software^103^ and LASX. Analysis was performed blindly using the manual track plugin from ImageJ (NIH, USA) to track cells with a migration time of at least 6 h.

### Histology and immunocytochemistry

For histology on embryonic brains, we used 50-µm coronal brain slices, cryosectioned with a CM3050 S Cryostate (Leica, Germany), and collected on Superfrost®Plus (Avantor, U.S.A.) object slides. On-slide immunohistochemistry was performed as described in Pensold et al. (2017)^28^. Briefly, all samples were washed for 5x 20 min in 1x PBS /0.5% Triton X-100/0.5% Tween®20 with constant horizontal shaking (50 rpm). After that, all samples were treated with blocking solution (4% bovine serum albumin (BSA) and 10% normal goat serum in 1x PBS/0.5% Triton X-100/0.5% Tween®20) was applied for 2 h at room temperature (RT). Primary antibody incubation diluted in blocking solution was conducted overnight at RT in a humid chamber. After washing for 5x 20 min in 1x PBS/0.5% Triton X-100/0.5% Tween®20, secondary antibodies were applied in blocking solution for 2h at RT. After washing for 3 x 20 min, the samples were stained with 4’,6-diamidino-2-phenylindol dihydrochloride (DAPI, 1:10000 in PBS, Carl Roth, Germany) for 15 min at RT, followed by washing twice for 5 min with PBS, sections were embedded in Mowiol or Fluoromount (Thermo Fisher, U.S.A.).

For adult brains, immunohistochemistry was performed in free-floating sagittal and coronal sections (30 µm), prepared using the CM3050 S Cryostate (Leica, Germany). Slices were transferred to 1x PBS and stored at 4°C. For sagittal sections, Bregma 1.32 and 1.44 were used, while Bregma 1.18, 1.10, 0.14, -0.22, -3.08, and -3.28 were taken for coronal sections. The same procedure as for embryonic sections was applied, except that an antigen retrieval with heated HistoVT One® (1x HistoVT One®/H_2_O bidest., Nacalai, Japan) at 70°C for 20 min was performed for staining against DNMT1. Sections that were stained for somatostatin, calretinin, and NPY underwent an antigen retrieval with citrate buffer (95°C; 10 mM, pH 6.0, supplemented with 0.5% Tween®20) for 15 min and subsequent cool-down for 30 min before washing. Finally, all slices were transferred onto glass slides in 0.5% (w/v) gelatine/0.05% (w/v) chromium(III) potassium sulfate dodecahydrate (KCrS_2_O_8_)/H_2_O solution prior to embedding in Mowiol.

Immunostaining on dissociated MGE and cortical single cells was conducted by washing coverslips with 1x PBS/0.1% Triton X-100 for 3x 5 min and blocking with 1x PBS/0.1% Triton X-100/4% BSA for 30 min at room temperature. Primary antibodies were applied for 2 h at room temperature. After washing for 3x 5 min with 1x PBS/0.1% Triton X-100, the secondary antibody was incubated for 1 h at room temperature. Phalloidin-647 diluted 1:1000 in 1x PBS (#ab176759, Abcam, U.S.A.) was applied for 20 min after washing. Then a final washing step with 1x PBS for 10 min and a DAPI staining (1:10000/1x PBS; Carl Roth, Germany) for 5 min were performed and coverslips were mounted in Mowiol.

The following primary antibodies were used: mouse anti-Calretinin (1:500; Swant, Switzerland, #6B3), rabbit anti-NPY (1:2500; Immunostar, U.S.A., #2940), rabbit anti-TBR1 (1:200; Abcam, U.S.A., #ab31940); rabbit anti-EOMES (1:500; Abcam, U.S.A., #ab23345); rat anti-Somatostatin (1:100; Millipore, U.S.A., #MAB354), mouse anti-Parvalbumin (1:2000; Swant, Switzerland, #235); rabbit anti-DNMT1 (1:100; Santa Cruz, U.S.A., # sc20701); rabbit anti-CUX1 (CDP; 1:100, Santa Cruz, U.S.A., #sc13024); mouse anti-Nestin (1:100; Merck U.S.A., #MAB353), rabbit anti-ß-Tubulin III (1:500; Sigma Aldrich, U.S.A., #T2200), and rabbit anti-ErB4 (1:1000; Proteintech, U.S.A., #22387-1-AP).

The following secondary antibodies conjugated with respective fluorophores were used at 1:1000 dilutions: Alexa488-goat anti-Rat IgG (Invitrogen, U.S.A., #A11006) Cy5-Goat anti-Rabbit IgG (Life Technologies, U.S.A., #A10523), A488-Donkey anti-Mouse IgG (Jackson, U.S.A., #15454150), Cy5-Goat anti-Mouse IgG (Jackson, U.S.A., #115175146), A488-Goat anti-Rabbit IgG (Life Technologies, U.S.A., #A11008).

### Transcriptome data on neurons overexpressing DNMT1

*Dnmt1^tet/tet^* mouse embryonic stem cells have been previously described^104^. The levels of DNMT1 in these cells are four times higher than the normal wild-type *R1* ESCs. Upon differentiation, the induced *Dnmt1^tet/tet^* neurons produced contain twice the DNMT1 levels than in *R1* neurons^104^. Differential expression gene data on *Tet/Tet* neurons was taken from Singh *et al*. (2023)^44^ and genes with significantly altered transcript levels were compared with the transcriptome data on the tissues lacking DNMT1.

### Enrichment of E14.5 tdTomato cells for RNA and methyl-sequencing

For FACS-mediated enrichment of *Sst-Cre*/*tdTomato* cells, telencephalons were prepared from E14.5 embryos and subjected to cell dissociation. Nuclease-free reaction tubes were used during isolation and long-term storage of the resulting material. The telencephalons were collected in cold HBSS (w/ phenol red, w/o calcium, w/o magnesium)/0.65% D-glucose; 4 µg/µL (600 U) of DNAse I (AppliChem GmbH, Germany). After treatment with 0.04% trypsin for 17 min at 37 °C, HBSS was replaced by DMEM with additional L-glutamine and 4.5 g/L D-glucose, 10% FBS and 1% P/S to stop the trypsinization. Subsequently, the cells were pelletized, resuspended and triturated in cold HBSS (w/o phenol red, w/o calcium, w/o magnesium)/0.65% D-glucose, before being filtered through a nylon gauze (pore size 140 µm, Merck, U.S.A.) for FACS.

FACS was performed by the Flow Cytometry Facility (FCF, University Hospital RWTH Aachen, Germany). Respective parameters for the procedure using a BD FACS Aria Fusion (BD Biosciences, U.S.A) were defined as follows: 5-laser (FCS, SSC, PE, BV421), 18-color (3-6-2-4-3). TdTomato-positive cells were either collected in 100 µL of cold TRIzol™ (Thermo Fisher Scientific, U.S.A.) for subsequent RNA sequencing or in 100 µL of cold HBSS (w/o phenol red, w/o calcium, w/o magnesium)/0.65% D-glucose for further processing for DNA methylation analysis. Finally, all samples were stored at -80 °C.

### Total RNA seq

TdTomato-positive E14.5 *Sst-Cre/tdTomato or Sst-Cre/tdTomato/ Dnmt1 loxP^2^* cells from the basal telencephalon and the cortex were FAC sorted into 100 µL TRIzol™ reagent (Thermo Fisher Scientific, U.S.A.). Samples were mixed by inversion and stored at -80 °C until processing. RNA was isolated from 50,000 cells pooled per genotype. The samples were filled up to 1 ml with TRIzol™ and dounced 25 times in a glass homogenizer using the small-clearance pestle. Samples were incubated for 5 min at room temperature, supplemented with 200 µL chloroform, and mixed by shaking vigorously until a homogenous milky solution appeared. Following another incubation at room temperature (until a visible phase separation appeared) samples were centrifuged at 13,000x *g* and 4 °C for 15 min in a tabletop centrifuge with cooling function. For precipitation, the clear upper phase was collected and mixed with 70% ethanol. Samples were bound onto RNeasy mini columns (Qiagen, Germany) and washed according to the instructions in the manual. RNA was eluted with 30 µL nuclease-free water.

RNA concentrations were determined using a Qubit™ 4 flourometer (Thermo Fisher Scientific, U.S.A.) with RNA HS reagents (Thermo Fisher Scientific, U.S.A.). RNA integrity was checked on a Tape Station 4200 using the HS RNA kit (Agilent Technologies, U.S.A.).

Prior to library preparation, samples were digested with HL dsDNAseI (ArcticZymes Technologies ASA, Norway) for 10 min at 37 °C to remove traces of gDNA. To avoid digestion of the newly generated cDNA during library preparation, the dsDNAseI was heat-inactivated for 5 min at 58 °C. Thereafter, samples were subjected to library preparation using the TAKARA SMARTer Stranded Total RNA-Seq Kit v3 - Pico Input Mammalian for Illumina (Takara Bio, Japan). Fragmentation was done at 94°C for 4 min. Illumina adaptors and indices were ligated to cDNA in a PCR reaction with 5 cycles (98 °C for 15 s, 55 °C for 15 s and 68 °C for 30 s). RRNA depletion was performed on the amplified libraries. Final libraries were subjected to an upscale PCR with 13 cycles (98 °C for 15 s, 55 °C for 15 s and 68 °C for 30 s). Amplified libraries were cleaned with magnetic AMPure XP beads (Beckman Coulter, U.S.A.). Library concentration was determined with a Qubit™ 4 fluorometer using the 1x dsDNA HS reagent (Thermo Fisher Scientific, U.S.A.). Library size was assessed on a Tape Station 4200 using the DNA D1000 kit (Agilent Technologies, U.S.A.).

Libraries were diluted, equimolarly pooled, denatured, and loaded for clustering onto an Illumina NovaSeq 6000 SP (200 cycles) flow cell (Illumina, U.S.A.) in conjunction with 1% PhiX control library, and run on an Illumina NovaSeq machine (Illumina, U.S.A.) in 75 bp paired-end mode. The 1% PhiX control library was spiked in to improve base calling accuracy.

*FASTQ* files were generated using *bcl2fastq* (Illumina, U.S.A.). To facilitate reproducible analysis, samples were processed using the publicly available *nf-core/rnaseq* pipeline version 3.12^105^ implemented in Nextflow 23.10.0^106^ with minimal command. In brief, lane-level reads were trimmed using *Trim Galore* 0.6.7^107^ and aligned to the mouse genome (GRCm39) using *STAR 2.7.9a*^108^. Gene-level and transcript-level quantification was done by *Salmon v1.10.1*^109^. All analysis was performed using custom scripts in *R versi*on 4.3.2 using the *DESeq2 v.1.32.0* framework^110^.

### MethylSeq

TdTomato-positive E14.5 *Sst-Cre/tdTomato* and *Sst-Cre/tdTomato/ Dnmt1 loxP^2^* cells were FAC-sorted into 100 µL HBSS. Samples were mixed by inversion and stored at -80°C until processing.

DNA was isolated from at least 50.000 pooled cells per sample using the PureLink genomic DNA Mini Kit (Thermo Fisher Scientic, U.S.A.). Per genotype, the samples were initially pooled by volume (200 uL each), supplemented with lysis buffer provided in the kit and incubated with 2 µL proteinase K (Thermo Fisher Scientific, U.S.A.) at 55 °C for 10 min. DNA was precipitated with 100% ethanol. Samples were bound to one column per genotype each, washed as instructed, briefly dried and eluted with 35 µL Tris-EDTA (TE) buffer. DNA concentrations were determined using the Qubit^TM^ 4 flourometer with the 1x dsDNA HS reagent (Thermo Fisher Scientic, U.S.A.). Quality/integrity was checked on a Tape Station 4200 using the Genomic DNA kit (Agilent Technologies, U.S.A.).

Libraries were generated using the Enzymatic Methyl-seq kit for Illumina (#E7120, New England Biolabs, U.S.A.). After enzymatic shearing for 25 min at 37 °C with the NEBNext UltraShear enzyme and buffer, followed by an inactivation at 65 °C for 15 min, 5-methylcytosines were oxidized by TET2 after adaptor ligation. Samples were denatured with formamide and deaminated with APOBEC. Libraries were subjected to 7 cycles of upscale PCR (98 °C for 10 s, 62 °C for 30 s, and 65 °C for 60 s). Library concentrations were determined with a Qubit^TM^ 4 fluorometer and the 1x dsDNA HS reagent (Thermo Fisher Scientific, U.S.A.). All recommended clean-up steps were performed with magnetic AMPure XP beads (Beckman Coulter, U.S.A.). Library size was assessed on a Tape Station 4200 using the DNA D5000 kit (Agilent Technologies, U.S.A.). Concentrations were determined with a Qubit^TM^ 4 fluorometer and the 1x dsDNA HS reagent (Thermo Fisher Scientific, U.S.A.).

Libraries were diluted, equimolarly pooled, denatured, and clustered onto an Illumina NextSeq 500/550 Mid Output Kit v2.5 (300 cycles) flow cell (Illumina, U.S.A.). The 1% PhiX control library was spiked in to improve base calling accuracy. Paired-end sequencing was performed with 151 cycles.

FASTQ files were generated using bcl2fastq (Illumina, U.S.A.). To facilitate reproducible analysis, samples were processed using the publicly available nf-core/methylseq pipeline version 2.6.0^105^ implemented in Nextflow 23.10.0^106^ with the minimal command. Briefly, FastQC (http://www.bioinformatics.babraham.ac.uk/projects/fastqc) and Trimmomatic^111^ were utilized to perform quality control, and clean reads were aligned to the reference genome using Bismark^112^ to account for bisulfite conversion and identify methylated cytosines. Differentially methylated sites and regions were called by the R package DSS^113^ and annotated using the GENCODE VM23 dataset. Regions were similarly annotated with a margin of 1 kb in front of the TSS. Methylation tracks were converted from mm10 to mm39 using the USCS LiftOver tool^114^ for integration with RNA seq data and visualized in IGV^115^. The sequence of the promoter regions (obtained from Ensembl release 112^116^) within these limits was further analyzed for the enrichment of previously identified DNMT1-binding motifs using the AME algorithm provided within the MEME suite^117^. Gene sets of interest were analyzed for the enrichment of gene ontology terms using ShinyGO 0.80 with the biological processes’ dataset^118^.

### Single-cell RNA-sequencing

The dorsal telencephalons from E14.5 C57BL/6J mice were dissected. Single-cell suspensions were prepared using a papain dissociation system (#LK003150; Worthington Biochemical Corporation, U.S.A) and all centrifugation steps were done at 4°C. Cells were manually counted using Trypan Blue exclusion, which consistently showed a viability greater than 95%.

ScRNA-seq libraries were prepared using the 10x Genomics 3’ Gene Expression Kit v3.1 (10x Genomics, U.S.A). Each sample had a target of 10’000 cells and was independently loaded in the Chromium Controller (10x Genomics, U.S.A). Quality control of the cDNA and final libraries was conducted using the High Sensitivity RNA D5000 and D1000 ScreenTape assays (Agilent Technologies, U.S.A.), respectively. Libraries were pooled and sequenced on a NovaSeq 6000 using the S2 Reagent Kit v1.5 with 100 cycles (Illumina, U.S.A).

*FASTQ* files were processed using *CellRanger v7.0.0*. All bioinformatics analysis was performed in *R 4.4.0* running under *Ubuntu 20.04.6 LTS*. The *Seurat v5* package^119^ was used for filtering, quality control, clustering, and annotation of the data. *CellChat v2.1.2*^61^ was used to compute the cell communication probabilities among different clusters.

### Single-nucleus RNA-sequencing

The medial neocortices from E16.5 *Sst-Cre/tdTomato* and *Sst-Cre/tdTomato/Dnmt1 loxP^2^* mice were dissected. The samples were snap-frozen on dry ice and stored at -80 °C until library preparation to ensure RNA integrity was maintained while the genotype and the sex of each sample could be verified for a bias-free transcriptomic analysis. Consequently, single-nucleus RNA sequencing (snRNA-seq) was selected over scRNA-seq to accommodate the processing of frozen samples. The nuclei were extracted from the frozen tissue by discontinuous sucrose gradient ultracentrifugation. Briefly, the tissue was homogenized in 1 ml of lysis buffer (0.32 M sucrose, 10 mM Tris-HCl (pH 8.0), 5 mM CaCl2, 3 mM Mg acetate, 1 mM dithiothreitol (DTT), 0.1 mM EDTA, 0.1% Triton X-100, 50 U/ml RNase inhibitors (#M0314S, New England Biolabs, U.S.A.)) using a 2-ml glass Dounce homogenizer on ice and stroking 25 times with pestle A followed by 100 strokes with pestle B. The homogenates were transferred to 5-ml ultracentrifuge tubes (#344057, Beckman Coulter) and under-layered with 3 ml of sucrose cushion (1.8 M sucrose, 10 mM Tris-HCl (pH 8.0), 3 mM Mg acetate, 1 mM DTT, 50 U/ml RNase inhibitors). The samples were loaded onto a Sw-55 Ti rotor and centrifuged for 1 h at 4 °C and 28,300 rpm (97,500x *g*). The nuclei pellets were resuspended in 150 µl of cold 1% BSA in PBS and immediately fixed using the Evercode™ Nuclei Fixation kit (Parse Biosciences, U.S.A.) following the manufacturer’s guidelines. The fixed nuclei were stored at -80 °C until library preparation.

The snRNA-seq libraries were generated with the Evercode™ WT kit (Parse Biosciences, U.S.A.). The quality check of the cDNA and the final libraries was performed using the High Sensitivity D5000 and D1000 ScreenTape kits (Agilent Technologies, U.S.A.). Libraries were equimolarly pooled and sequenced in conjunction with 5% PhiX control library on an Illumina NovaSeq machine in paired-end mode using the S2 Reagent Kit v1.5 (200 cycles) (Illumina, U.S.A.).

The sequencing dataset was processed using the TrailmakerTM (https://app.trailmaker.parsebiosciences.com/; Parse Biosciences, 2024). Unfiltered count matrices were further processed using the Seurat package for R (R 4.4.1, Seurat 5.0.1.9001)^119^. Cells were filtered based on counts, discarding cells below 1500 and above 60000 reads per cell. Dead or dying cells were removed by filtering droplets with high mitochondrial content (10% cut-off).

Subsequently data was normalized prior to principal-component analysis (PCA) and Leiden-clustering. A Uniform Manifold Approximation and Projection (UMAP) embedding was calculated to visualize the results. Cluster-specific marker genes were identified by comparing cells of each cluster to all other cells using the presto package implementation of the Wilcoxon rank-sum test. The top marker genes for each cluster were cross-referenced with known cell type-specific markers from the literature and publicly available databases.

The Annotated Clusters were subsequently used to infer cell-cell communication probabilities using the CellChat Database and corresponding R package (CellChat version 2.1.2)^61^.

### MERSCOPE Analysis

Spatial transcriptomic analysis was performed on 10 µm brain sections from fresh-frozen E16.5 *Sst-Cre/tdTomato* and *Sst-Cre/tdTomato/Dnmt1 loxP^2^* embryos using the MERSCOPE™ system (Vizgen) according to the manufacturer’s instructions. Briefly, tissue samples were mounted onto specialized slides and fixed to preserve RNA integrity. Multiplexed Error-Robust Fluorescence In Situ Hybridization (MERFISH) was then applied, where gene-specific oligonucleotide probes hybridized to target RNA molecules. We used the commercially available Pan Neuro panel covering 500 transcripts used for the identification of cell types in the mouse Brain.

Following hybridization, high-resolution fluorescence imaging was conducted on the MERSCOPE platform to capture the spatial distribution of individual transcripts at subcellular resolution. The imaging data was preprocessed on the instrument, which decodes RNA signals, assigns molecular identities, segments the tissue into cells and quantifies transcript abundance across individual cells within the tissue.

Two brain tissue sections from comparable anatomical planes were analyzed using Seurat in R *(R 4.4.1, Seurat 5.0.1.9001)*. After normalization and identification of highly variable features for each dataset, the datasets were integrated using reciprocal PCA to account for shared sources of variation. Clustering was performed using the Louvain algorithm at a resolution of 0.8, and clusters were manually annotated based on marker genes identified through differential expression analysis. Heterogeneous clusters were further resolved through subclustering, with iterative marker analysis and annotation revealing additional distinct cell types and states within the brain tissue.

### Neuropixels recordings and behavioral experiments Surgical Procedures

Mice received systemic analgesia prior to surgical procedures in form of subcutaneous injections of carprofen (4 mg/kg, Rimadyl, Zoetis GmbH, Germany) and buprenorphine (0.1 mg/kg, Buprenovet sine, Bayer Vital GMBH, Germany) and were then anesthetized using (1%–2.5%) isoflurane in oxygen. The eyes were covered with eye ointment (Bepanthen, Bayer Vital GmbH, Germany) to prevent them from drying out. Following a local injection of bupivacaine (0.08 ml of 0.25% Bucain 7.5mg/ml, Puren Pharma GmbH, Germany), the scalp was incised and pushed outward to fix it in place using tissue adhesive (Vetbond, 3M). A circular headbar was attached on top using dental cement (C&B Metabond, Parkell U.S.A.; Ortho-Jet, Lang Dental). For electrophysiological recordings, craniotomies were performed to inject a viral vector into the cortex (AAV1.shortCAG.dlox.hChR2(H134R).WPRE.hGHp, Zurich Vector Core, viral titer = ∼8x10^12^ vg/ml). We targeted primary somatosensory (S1) and visual (V1) cortex (stereotactic coordinates; S1: -1.5 mm anteroposterior and -3.5 mm mediolateral from Bregma; V1: -4 mm anteroposterior and -2 mm mediolateral from Bregma) and injected a volume of 200 nl in bouts of 10 nl with a 14-s delay between bouts (Nanoject III, Drummond Scientific, U.S.A.). For each coordinate, injections were performed at two cortical depths (300 and 600 µm), adding up to a total volume of 400 nl per injection site. The glass pipette was removed at least 5 min after the full volume was injected to reduce the risk of backflow. After viral injections a 3-mm wide round coverslip was placed inside the craniotomy and fixed with light-curable dental cement (DE Flowable composite, Nordenta, Germany). The coverslips contained a small opening (∼0.2 mm) that was covered with silicon to allow access of the Neuropixels probes to the cortex. After the surgery, all animals received the same injections of carpofen and buprenorphine as in the beginning. They also received buprenorphine (0.009 mg/ml, Buprenovet sine, Bayer Vital GmbH, Germany) and enrofloxacin (0.0227 mg/ml, Baytril 5%, Bayer Vital GMBH, Germany) in their drinking water for three days following the surgery. Behavioral training started after at least two weeks of recovery

### Neuropixels recordings

Electrophysiological recordings were done with Neuropixels 1.0 Probes in head-fixed mice, freely running on a wheel. For each mouse, we performed four recordings on consecutive days. To later recover the probe position in each recording, the probes were painted with DiD cell labeling solution (Invitrogen V22887, U.S.A.) before each recording. To record neural responses to tactile stimulation across all cortical layers, the probes were inserted at an orthogonal angle to the brain surface (∼10°). The stimulation protocol was started 5–10 min after insertion, we recorded high pass-filtered data above 300 Hz at 30 kHz and low pass-filtered signals between 0.5–100 Hz at 2.5 kHz from the bottom 384 channels of the Neuropixels probe (∼3.8 mm active recording area). Signals were acquired with an external Neuropixels PXIe card (IMEC, Belgium) used in a PXIe-Chassis (PXIe-1071, National Instruments, USA). Triggers and control signals for different stimuli and wheel movements were separately recorded as analog and digital signals using the SpikeGLX software (Janelia Farm Research Campus, U.S.A.; Bill Karsh).

Visual stimulation to induce gamma oscillations consisted of 5-s long full-field drifting square wave gratings at horizontal orientation and a spatial frequency of 0.04 cycles per degree. All visual stimuli were presented at a 17 cm distance from the right eye on a gamma-corrected LED-backlit LCD monitor (Viewsonic VX3276-2K-MHD-2, 32”, 60Hz refresh rate). Tactile stimulation was controlled by a microcontroller-based finite state machine (Bpod r2, Sanworks, U.S.A.) using custom Matlab code (2020b, MathWorks). Tactile stimuli consisted of 20-ms long air puffs at 0.03 bar pressure directed at the distal parts of the whiskers. In each trial, two air puffs were presented at an inter-stimulus interval of 500 ms. The overall stimulation protocol consisted of 50 tactile trials and 50 visual trials in randomized order. Inter-trial intervals were also randomized between 3–5 seconds.

To extract spiking activity, channels that were broken or outside of the brain were detected and removed from the recording using the SpikeInterface analysis toolbox. The remaining channels were time-shifted and median-subtracted across channels and time. Corrected channels were then spike-sorted with the Kilosort 2.5 software and manually curated using phy. All sorted spiking and local field potential data were analyzed using custom Matlab code (2020b, MathWorks).

Peri-event time histograms (PETHs) for sensory responses in each cluster were computed with a bin size of 10 ms and baseline-subtracted by the mean activity within 1 second before the laser onset (between 2 to 1 seconds before the first tactile stimulus). Trial-averaged local field potentials (LFPs) in response to sensory stimulation were similarly baseline-corrected. To compute current source densities (CSDs) from LFP signals, we used the inverse CSD method by Pettersen et al. (2006)^75^. We applied the spline iCSD method, assuming a smoothly varying CSD between electrode contacts, based on polynomial interpolation. We assumed a homogeneous, isotropic conductivity of σ = 0.3 S/m within and directly above the cortex^75^. To reduce spatial noise, the estimated CSD was subsequently convolved with a Gaussian spatial filter with a standard deviation of 0.1 mm. For LFP and CSD analyses, the signals from neighboring contacts at the same depth were averaged together to improve signal quality.

### Evidence accumulation task

Head-fixed, water-deprived mice were stepwise introduced to performing a series of uni- or multisensory evidence accumulation tasks in a custom-built setup. The stimulus presentation, lick detection and reward delivery were driven by a microcontroller-based finite state machine running on a *Teensy 3.2* (PJRC) and using custom *Python* code (Python 3.7). For all trials, the general scheme of events was similar: for 3 seconds, randomized sequences of up to 6 visual, tactile or visuotactile stimuli were presented on one or both sides of the mouse. After a 0.5 s-long delay period, the animal had to respond within 2 seconds to obtain a reward. The ITI between trials was 3.5 seconds. Mice were first trained on reliably detecting visual stimuli, consisting of 6 visual gratings (spatial frequency = 0.018 cycles per degree) that were moving over the screen at a temporal frequency of 2 Hz. After mice learned to accurately respond on the side, where visual stimuli were presented, tactile stimuli were introduced in the form of 6 air puffs, as described above (see Neuropixels Recordings). Air puffs occurred at regular intervals of 0.5 seconds during the 3 second stimulus period. After mice also accurately detected the side of tactile stimulation, visuotactile stimuli were introduced where coherent visual and tactile stimuli were presented. Once mice achieved stable detection performances in all modalities, distractor stimuli of the same modality were introduced on the opposing side. While the number of stimuli in these trials varied to confront animals with different levels of difficulty, the general goal was for the mice to indicate the side with the higher number of presented stimuli.

### Morris water maze (MWM)

Visual navigation and learning capacities were investigated using a MWM. Mice were trained to search a translucent platform (Ø 10 cm) in a circular pool (Ø 1 m) containing tepid water. The pool was divided into four virtual quadrants, each of them equipped with distinct visual cues. Mice were tested on 6 consecutive days. Start positions for each trial were chosen at random. Mice remained on the platform for at least 5 s after finding it, to enable memorization of spatial cues. If a mouse did not find the platform within the given time limit, it was manually guided to the platform. In between trials, mice were kept underneath a red lamp for at least 10 minutes. Animals were habituated to the task on their first day of experiments by performing four trials for a maximum duration of 90 seconds each. The translucent platform slightly protruded from the water surface to facilitate quick locating. On the following 5 training days, the platform was submerged 1-2 cm below the water surface. All animals conducted 12 trials per day with three starts per quadrant, respectively. The maximum duration per trial was set to 60 seconds. Mice were recorded from above and behavioral data were processed using ANY-maze (Stoelting Co., U.S.A).

### Nest-building procedure

Nest-shredding and -building tests were performed with male mice between 12 and 24 weeks of age. Both *Sst-Cre* strains were tested by placing respective animals in separate housing cages with ad libitum access to food and water but without further enrichment except for fresh bedding and nest-building material (standardized autoclaved nestlets with a weight of 2–3 g (5 cm x 5 cm, Lexx, Germany). For each nestlet the weight was determined before the experiment which took place during the murine active phase (5 p.m. to 9 a.m.). After testing, all mice were transferred to their home cages and the percentual shredded material as well as the corresponding nest building score^95^ were recorded. Mice were tested twice within one week with two days between both trials. The resulting data were averaged for each mouse.

### Pentylenetetrazole (PTZ)-induced epileptic seizure protocol

Male *Sst-Cre/tdTomato* and *Sst-Cre/tdTomato/Dnmt1 LoxP^2^* mice (13- to 17-week-old) underwent an intraperitoneal administration of pentylenetetrazole (PTZ, Sigma-Aldrich, U.S.A.) with 10 mg/kg living weight. PTZ was diluted in sterile Ringer’s solution (pH 7.3–7.4) to a concentration of 5 µg/µL beforehand. All mice received an intraperitoneal injection of PTZ (40 µL PTZ-solution per 20 g living weight) every ten minutes. All tested animals were monitored in transparent cages with no additional enrichment. Within every period of ten minutes, the occurrence of different epileptic events ranked from severity level 0 (no abnormalities) to most severe score 7 (final tonic-clonic seizure with overstretched limbs and tail) were detected. For each period of ten minutes, all detected severity scores were added up and compared between *Sst-Cre/tdTomato* and *Sst-Cre/tdTomato/Dnmt1 LoxP^2^* mice. Animals that depicted a final tonic-clonic seizure before the ending of the experiment (maximum 12 injections and spending 150 min in the setup) were not included in the calculations after their final epileptic event (score 7). Respective severity levels and corresponding events are depicted in Figure 6 following Racine’s scoring. After PTZ-protocol, all mice were sacrificed with an overdose of 5 vol% isoflurane followed by cervical dislocation after reaching surgical tolerance, verified by loss of withdrawal response to toe pinch.

### Statistics and Figure illustration

Statistical tests were performed using Matlab, Phyton, and GraphPad Prism. Unless otherwise stated, normally distributed data was tested applying unpaired and two-tailed Student’s *t*-test and an unpaired Welch’s *t*-test, nested two-way ANOVA, and two-way ANOVA followed by Bonferroni correction. For non-normally distributed data we used a Wilcoxon *rank-sum* test. Respective statistical tests can be found in the main text and individual captions.

Unless otherwise stated, all depicted error bars represent the standard error of the mean (SEM). Violin plots show the distribution of the numeric data using density curves, with the respective frequency of data points represented by the width. Straight lines represent the median and dotted lines the corresponding interquartile range. Single-value plots (XY-graphs) display all detected data points together with their mean and SEM used to illustrate analyses including two-way nested ANOVA tests. Data derived from one specimen (embryo, adult mouse) are illustrated by different shapes of the respective data points and can therefore be distinguished from values derived from other animals. For boxplots, the central box represents the interquartile range (IQR), with the median denoted as a horizontal line within the box. Moreover, the 25th percentile (Q1) and the 75th percentile (Q3) of the data are shown. The whiskers extend to the data’s minimum and maximum values.

## Data Availability

The sequencing datasets generated and/or analyzed for this study are available in the Gene Expression Omnibus repository with the following accession numbers: GSE276510, GSE276512, and GSE276516. The MERESCOPE and sc/snRNA datasets are currently deposited at GEO and accession numbers will be available upon acceptance. MD simulation data are uploaded here: https://pubmed.ncbi.nlm.nih.gov/21070939/. Neuropixels recording data are uploaded here: https://doi.org/10.6084/m9.figshare.28282838.

## Code Availability

Not applicable.

## Acknowledgements

This work was supported by the Flow Cytometry Facility and the Sequencing Facility, two core facilities of the Interdisciplinary Center for Clinical Research (IZKF) Aachen within the Faculty of Medicine at RWTH Aachen University. We thank Dr. Mira Jakovcevski, Hendro Langecker, Dorothee Hoffmann, Pia Döring and Ananya Sangeetha for their experimental support, and Sandra Brill for her support with mouse care and genotyping. We further thank Prof. Dr. Christoph Kuppe, Emilia Scheidereit and Vanessa Künstler for the access to and support with MERESCOPE.

## Author contributions

J.R. performed experiments and data analysis, figure illustration, manuscript editing; P.W. performed experiments and data analysis, figure illustration, manuscript editing; S.X. performed the *in silico* experiments, data analysis, and figure illustration; K.Z. performed the *in silico* experiments and data analysis; C.L.F. performed experiments and data analysis; J.D. performed experiments and data analysis; S.G. performed experiments and data analysis; J.L. performed experiments and data analysis; C.B.Y. performed experiments and data analysis; G.P. performed experiments and data analysis; G.N. methods support and data analysis; L.D. performed experiments and data analysis; J.V. performed experiments and data analysis; L.B. performed experiments and data analysis; S.K. performed experiments and data analysis; M.S. performed experiments and data analysis, K.N.M. Data analysis, data discussion; C.C.K. data analysis; T.V. data discussion and analysis, manuscript editing; P.C. data interpretation and discussion, wrote the *in silico* analysis; S.M. data analysis and discussion, figure illustration, manuscript preparation; G.Z.B. conceptual design, administration and supervision, experimental design, wrote the manuscript, figure illustration, data analysis, interpretation and discussion.

