## Extended data information for "DNMT1-Mediated Regulation of Somatostatin-positive Interneuron Migration Impacts Cortical Architecture and Function"

### Extended Data Figure Legends

#### Extended Data Figure E1:

**Molecular dynamics simulations of DNMT1 binding to unmethylated DNA (UMDNA).**

**(a-f)** Inter-unit DNMT1 proteins interacting with the DNMT1/UMDNA/SAH complex unit in the X-ray structure (PDB ID: 3PTA). **(a)** Intra-unit DNMT1 regions shown including the CXXC domain (yellow), which interacts most with UMDNA (blue), autoinhibitory linker BAH1/BAH2 (white surface), catalytic domain (orange), and SAH (red spheres). The neighboring inter-unit DNMT1 proteins are illustrated in rose-pink. **(b)** Electrostatic surface representation of the complex observed in the X-ray structure. Surfaces are colored as in **(a)**. The additional red and blue surfaces represent negative and positive potential regions, respectively. The inter-unit DNMT1 regions the UMDNA interacts with, are mostly positively charged. **(c-f)** Number of contacts of the 5' and 3' regions with their intra-unit and inter-unit DNMT1 with different distance cutoffs in the X-ray structure. **(g, h)** Simulations of DNMT1/UMDNA interactions in aqueous solution. The complexes shown are at the end of replica 2 and replica 3 with the same coloring scheme as **(a)**. Number of contacts between DNMT1 with UMDNA **(g´, h´)**, 5' regions **(g´´, h´´)**, and 3' regions **(g´´´, h´´´)** as well as between 5' region and the catalytic domain of DNMT1 **(g´´´´, h´´´´)** during last 100 ns simulations.

**(i-k)** The heavy-atom root-mean-square deviation (RMSD) values for each component were monitored across three independent MD simulations with **(i)** depicting replica 1, **(j)** depicting replica 2, and **(k)** depicting replica 3. UMDNA exhibited significant conformational changes in all replicas, with RMSD values exceeding 10 Å.

#### Extended Data Figure E2:

**Validation of SST expression and reduced DNMT1 levels in tdTomato-positive cells. (a)** Mating strategy to obtain *Sst-Cre/tdTomato* (control) and *Sst-Cre/tdTomato/Dnmt1 loxP*$^{2}$ (KO) mice. *Sst*$^{tm2.l(cre)Zjh}$/J mice were mated with *tdTomato* reporter mice (*B6.CgGt(ROSA)26Sor*$^{tml\ 4(CAG\text{-}tdTomato\ )Hze}$) resulting in *Sst-Cre/tdTomato* mice with an internal ribosomal entry site (*IRES*), a *Cre-recombinase* sequence, a *polyA* sequence, and a *frt*-flanked neo cassette located in the 3' untranslated region (UTR) of the *Somatostatin* (*Sst*) locus on chromosome 16, limiting the respective *Cre* expression to *Sst*-positive neurons. The *tdTomato* sequence and a loxP-flanked stop cassette are located in the *Rosa26* locus. Triple transgenic *Sst-Cre/tdTomato/Dnmt1 loxP*$^{2}$ mice were obtained by crossing control mice with a *Dnmt1 loxP*$^{2}$ strain (*B6; 129Sv-Dnmt1tm4Jae/J*; exon 4 and 5 of the *Dnmt1* gene are *loxP*-flanked, resulting in a null allele of these loci and a subsequent DNMT1 deficiency. **(b)** Exemplary microphotographs depicting SST expression in tdTomato-cells in sagittal brain sections of a four-month-old male *Sst-Cre/tdTomato* mouse (Bregma 2.76). **(c)** Magnified images of the areas outlined in (**b**). Scale bars: 100 $\mu$m in (**b**) and 20 $\mu$m (**c**). **(d-g)** Quantification of DNMT1 expression by immunohistochemistry in *Dnmt1* control **(d, e)** and *Dnmt1* KO **(f, g)** male mice (six-month-old individuals) in 30 $\mu$m, fixated coronal cryosections. Scale bars **(d, f)** 100 $\mu$m and 10 $\mu$m **(e, g)**. **(h)** Quantified ratio of the normalized mean grey value of the DNMT1 immunofluorescence signal in *tdTomato*-positive cells to the normalized mean grey value of

DNMT1 immunofluorescence of all cortical nuclei (identified by DAPI, shown in blue). Bregma 1.18, 1.10, 0.14, -0.22, -2.92. -3.08, and -3.28 were used for quantification. Statistical testing was conducted by using a nested two-way ANOVA ($p < 0.001$ ***, $N = 3$ for *Sst-Cre/tdTomato* ($n = 18$ slices) and *Sst-Cre/tdTomato/Dnmt1 loxP*$^2$ ($n = 17$ slices)). Data points represented using identical symbols in the plot originate from the same mouse. **(i)** *Dnmt1* transcript counts determined by RNA sequencing of FAC-sorted *Sst-Cre/tdTomato* and *Sst-Cre/tdTomato/Dnmt1 loxP*$^2$ cells from the E14.5 cortex. 3 libraries were analyzed for both genotypes from pooled material of $n = 7$ *Sst-Cre/tdTomato* embryos, and $n = 8$ *Sst-Cre/tdTomato/Dnmt1 loxP*$^2$ embryos; $\log_2$FC = -0.26482506, $p_{adj}$: 0.0121078). **(j-n)** Quantification of DNMT1 expression by immunohistochemistry in *Dnmt1* control **(j, k)** and *Dnmt1* KO **(l, m)** E14.5 embryos in 50 $\mu$m, fixated coronal cryosections. Scale bars 50 $\mu$m in **(j, l)** and 25 $\mu$m **(k, m)**. **(n)** Quantified ratio of the normalized mean grey value of the DNMT1 immunofluorescence signal in tdTomato-positive cells to the normalized mean grey value of DNMT1 immunofluorescence of all cortical nuclei (identified by DAPI, shown in blue). Statistical testing was conducted by using a nested two-way ANOVA ($p < 0.01$ **, $N = 3$ for *Sst-Cre/tdTomato* ($n = 22$ slices) and *Sst-Cre/tdTomato/Dnmt1 loxP*$^2$ ($n = 18$ slices)). Data points originating from the same mouse are plotted using the same symbol. FC = foldchange

### Extended Data Figure E3:

**RNA Sequencing and methyl-sequencing analysis of FAC-sorted *Sst-Cre/tdTomato/Dnmt1 loxP*$^2$ neurons compared to *Sst-Cre/tdTomato* control cells from the E14.5 basal telencephlaon**

**(a-c)** Gene ontology analysis and characteristics of genes that were upregulated in expression and differentially methylated in FAC-sorted *Sst-Cre/tdTomato/Dnmt1 loxP*$^2$ neurons compared to *Sst-Cre/tdTomato* control cells prepared from the E14.5 basal telencephalon. Background is defined as all detected transcripts in both genotypes; Chi-squared and Student's *t*-tests were run to analyze if our gene set has special characteristics when compared with all the other genes (ShinyGO 0.80; http://bioinformatics.sdstate.edu/go/). **(a)** Gene ontology (GO) analysis – Biological Process, **(b)** bar plots (tested by Chi-squared test), and **(c)** density plots (Student's *t*-test). **(d-f)** Gene ontology analysis and characteristics of genes that were upregulated in expression in FAC-sorted *Sst-Cre/tdTomato/Dnmt1 loxP*$^2$ neurons compared to *Sst-Cre/tdTomato* cells prepared from the E14.5 basal telencephalon. Background is defined as all detected transcripts in both genotypes; Chi-squared and Student's *t*-tests were run to analyze if our gene set has special characteristics when compared with all the other genes (ShinyGO 0.80; http://bioinformatics.sdstate.edu/go/). **(d)** Gene ontology (GO) analysis - Biological Process, **(e)** bar plots (tested by Chi-squared test) and **(f)** density plots (Student's *t*-test) that depict the characteristics of this gene set compared with the entire set of detected transcripts.

### Extended Data Figure E4:

**Analysis of genes overlapping between upregulated genes in *Sst-Cre/tdTomato/Dnmt1-loxP*$^2$ cells and downregulated after *Dnmt1* overexpression in neurons obtained from ESCs**

**(a-c)** Gene ontology analysis and characteristics of genes that upregulated in *Sst* expressing interneurons of *Sst-Cre/tdTomato/Dnmt1-loxP²* mice and overlap with genes that are downregulated after *Dnmt1* overexpression (OE) in neurons obtained from murine embryonic stem cells (ESCs)[1]. **(a)** Gene ontology (GO) analysis for the overlap. **(b)** Further characteristics are shown in bar plots (tested by Chi-squared test) and density plots (Student's *t*-test) **(c)**, depicting the characteristics of this gene set compared with the entire set of detected transcripts.

### Extended Data Figure E5:

**The cortical size and dimensions of its different regions do not differ between *Sst-Cre/tdTomato/Dnmt1-loxP²* and *Sst-Cre/tdTomato* brains at E14.5, E16.5, and E18.5. (a)** Schematic representation of cortical regions and their extensions in a single hemisphere of 50 μm-thick coronal brain sections from E14.5 embryos. Quantifications of different parameters are depicted in **(b)**. Quantification was conducted by using a nested two-way ANOVA with $N = 3$ embryos for both genotypes, $n = 13$ slices for *Sst-Cre/tdTomato,* and $n = 9$ slices for *Sst-Cre/tdTomato/Dnmt1 loxP²* samples. **(c)** and **(e)** Schematic representation of cortical regions and their extensions in a single hemisphere of 50 μm-thick coronal brain sections from E16.5 and E18.5 embryos, respectively. The cortical size and dimensions of its different regions are not different in E16.5 and E18.5 *Sst-Cre/tdTomato/Dnmt1 loxP²* embryos compared to age-matched *Sst-Cre/tdTomato* individuals, as quantified in **(d)** for E16.5 and in **(f)** for E18.5. Quantification was conducted by using a nested two-way ANOVA with $N = 4$ embryos for both genotypes, $n = 12$ slices for *Sst-Cre/tdTomato,* and $n = 15$ slices for *Sst-Cre/tdTomato/Dnmt1 loxP²* for E16.5 **(d),** and $N = 4$ embryos for both genotypes, $n = 12$ slices for *Sst-Cre/tdTomato,* and $n = 9$ slices for *Sst-Cre/tdTomato/Dnmt1 loxP²* at 18.5 **(f).** MZ: marginal zone, CP: cortical plate, IZ: intermediate zone, SVZ+VZ: subventricular zone, n.s.: not significant. Data points with the same symbol belong to one embryo.

### Extended Data Figure E6:

**DNMT1 regulates ERBB4 expression and proper migration of MGE-derived cells but does not affect morphology. (a-c)** Live-cell imaging analysis of *Sst-Cre/tdTomato* and *Sst-Cre/tdTomato/Dnmt1 loxP²* cells in E14.5 organotypic brain slices (350 μm coronal sections, time-period of imaging: 20 h). Analysis of the radial migration velocity and migrated path length are shown in **(a)** and **(b)**, respectively. **(c)** DNMT1 does not impact the migratory morphology of tdTomato⁺ cells investigated in organotypic brain slices (350 μm coronal sections). Nested two-way ANOVA ($p < 0.01$ **) with $N = 3$ embryos and $n = 4$ slices for control and KO animals, respectively, $n = 185$ cells for control, and $n = 219$ KO cells. **(d-g)** siRNA-mediated depletion of *Dnmt1* in MGE-derived single cells *in vitro* (E14.5 + 1DIV) does not affect the cellular morphology compared to control siRNA-transfected cells. Scale bars: 10 μm. Quantification is shown in **(e-g)**. Unpaired two-tailed Student's *t*-test and unpaired Welch's *t*-test, $n = 60$ cells each for control siRNA and *Dnmt1* siRNA. siRNA: small-interfering RNA; LP: leading process, n.s.: not significant. **(h, i)** ERBB4 level is increased after *Dnmt1* knockdown (KD) in MGE cells (E14.5 + 1DIV). Exemplary microphotographs showing control siRNA-treated (upper panel) and *Dnmt1*-KD cells (lower panel) stained for Phalloidin647 and ERBB4. On the very right of each example, thermal-color-coded panels represent the respective fluorescent integrated density (IntDens) of ERBB4 (therm. (thermal) LUT). A dark blue color indicates 0 and a red color represents 5000 fluorescent units of the IntDens. Scale bars: 10 $\mu$m. **(i)** Quantification comparing the integrated density (IntDens) of ERBB4 fluorescence signals of control and *Dnmt1*-KD MGE cells. Unpaired two-tailed Student's *t*-test and unpaired Welch's *t*-test, $p < 0.001$ *** ($n = 88$ cells for control siRNA and 78 cells for *Dnmt1* siRNA, $N = 4$ experiments). Data points depicted in (a) and (c) with the same symbol derive from one embryo.

### Extended Data Figure E7:

#### *Dnmt1* deletion in *Sst-Cre/tdTomato* interneurons does not affect their cortical distribution in E16.5 and E18.5 embryos.

**(a)** Exemplary microphotographs of tdTomato⁺ interneurons in coronally sectioned (50 $\mu$m) hemispheres of E16.5 *Som-Cre/tdTomato* (control) and *Som-Cre/tdTomato/Dnmt1 loxP*$^2$ (KO) embryos; scale bars: 500 $\mu$m. TdTomato⁺ cells are labeled in red and DAPI in blue. **(b)** Quantitative analysis of the proportional distribution of tdTomato⁺ cells in the basal telencephalon and the cerebral cortex in E16.5 coronal sections (nested two-way ANOVA, $n = 11$ sections for control and $n = 15$ sections for KO, $N = 4$ embryos per genotype). **(c)** Magnified microphotographs of the cortices of control and KO embryos from E16.5 coronal brain sections (50 $\mu$m) to illustrate the distribution of tdTomato⁺ cells within the cortical zones. Scale bars: 100 $\mu$m. **(d)** Quantitative analysis of the proportional distribution of tdTomato⁺ cells within the E16.5 cortical zones normalized to the overall tdTomato-cell count within the cortex of control and KO embryos, analyzed in 50 $\mu$m coronal cryosections. Nested two-way ANOVA, $n = 13$ sections for control and $n = 14$ sections for KO, $N = 4$ embryos per genotype. **(e)** Quantitative analysis of tdTomato-cell density normalized to the given area of the respective cortical zones (E16.5, nested two-way ANOVA, $n = 13$ sections for control and $n = 14$ sections for KO, $N = 4$ embryos per genotype). **(f)** Exemplary microphotographs of tdTomato⁺ interneurons in the cortex of coronally sectioned (50 $\mu$m) brains of E18.5 *Som-Cre/tdTomato* (control) and *Som-Cre/tdTomato/Dnmt1 loxP*$^2$ (KO) embryos; scale bars: 100 $\mu$m. **(g)** Quantitative analysis of the proportional distribution of tdTomato⁺ cells within the E18.5 cortical zones normalized to the overall tdTomato-cell count within the cortex of control and KO embryos, analyzed in 50 $\mu$m coronal cryosections. Nested two-way ANOVA, $n = 11$ sections for control and $n = 9$ sections for KO, $N = 4$ embryos per genotype. **(h)** Quantitative analysis of tdTomato-cell density normalized to the given area of the respective cortical zones at E18.5 (nested two-way ANOVA, $n = 11$ sections for control and $n = 9$ sections for KO, $N = 4$ embryos per genotype). MZ: marginal zone, CP: cortical plate, IZ-VZ: intermediate zone to ventricular zone. $p < 0.05$ *, $p < 0.01$ **, $p < 0.001$ ***, n.s.: not significant. Error bars represent the standard error of the mean (SEM). Data points with the same symbol belong to one embryo.

### Extended Data Figure E8: scRNA Seq data.

**(a-e)** Single-cell RNA sequencing (scRNA-seq) of E14.5 dorsal telencephalons **(a-c)** and E16.5 cortices **(e, f)** from C57BL/6J mice. UMAP illustrating cell clusters at E14.5 are depicted in **(a)**, and *Dnmt1* and *Sst* expression across these cluster are shown in **(b)** and **(c)**, respectively. **(e, f)** Panels show significant ligand-receptor pairs involved in communication from SST$^+$ cINs to cortical progenitors, filtered by differentially expressed genes (DEG) from single-cell RNA-sequencing data. Communication probability is represented by dot color, and *p*-value by dot size, with *p*-values being computed using a one-sided permutation test. Panel (**e**) shows the communication with different apical progenitor (AP) populations at E16.5. Panel **(f)** illustrates communication with IPCs at E16.5.

**(f-i)** Single nuclear RNA (snRNA) sequencing of cortical cells prepared from E16.5 *Sst-Cre/tdTomato* and *Sst-Cre/tdTomato/Dnmt1 loxP*$^2$ embryos ($n = 2$ brains from $N = 2$ mice per genotype). **(f)** UMAP depicting the cell clusters determined by snRNA sequencing across all samples. **(g)** Volcano plot collecting the DEGs in *Sst*$^+$/*Gad2*$^+$ cells between E16.5 *Sst-Cre/tdTomato* and *Sst-Cre/tdTomato/Dnmt1 loxP*$^2$ samples. **(h)** Dot plot depicting marker gene expression in the different cell clusters. **(i)** UMAPs depicting the expression of relevant cluster determining genes.

**Extended Data Figure E9: Spatial transcriptomic analysis using MERFISH**

MERFISH was used to investigate the spatial distribution of different cell types in the developing mouse brain at E16.5 in *Sst-Cre/tdTomato* and *Sst-Cre/tdTomato/Dnmt1 loxP*$^2$ embryos. The 10 µm-thick coronal brain sections were processed and imaged with the MERSCOPE platform (Vizgen) and further analyzed on the single-cell level in R using Seurat. **(a)** Spatial distribution of identified clusters **(b)** Bar plot depicting the proportional distribution of cell types identified in cortical columns depicted in Figure 4. **(c)** Dot plot illustrating the expression of key marker genes across the identified cell clusters. **(d)** Distribution of select transcripts within control and knockout sections.

**Extended Data Figure E10: Phenotypic characterization of adult *Som-Cre/tdTomato* (control) and *Som-Cre/tdTomato/Dnmt1 loxP*$^2$ (KO) cortices.**

**(a-d)** TdTomato-expressing cells show immunoreactivity for neuropeptide Y and calretinin. Exemplary microphotographs of sagittal sections from a four-month-old *Sst-Cre/tdTomato* animal (Bregma 2.76). **(a** and **b)** Examples of immunohistochemical stainings using an anti-somatostatin antibody (Som, green) and an anti-neuropeptide Y antibody (NPY, magenta). TdTomato$^+$ cells are depicted in red. **(c** and **d)** Examples of immunohistochemical stainings using an anti-somatostatin antibody (Som, green) and an anti-calretinin antibody (Cal, magenta). TdTomato$^+$ cells are depicted in red. Scale bars in **(a)** and **(c)**: 100 µm. Scale bars in **(b)** and **(d)**: 20 µm (magnified views of sections outlined in **(a)** and **(c)**, respectively). **(e-h)** Immunostaining using an antibody directed against parvalbumin (PV) in 50 *µ*m brain sections of six-month-old **(e, f)** *Sst-Cre/tdTomato* (control) and **(g, h)** *Sst-Cre/tdTomato/Dnmt1 loxP*$^2$ (KO) mice. Exemplary microphotographs of immunostainings (tdTomato in red, PV in green, DAPI in blue) taken from the primary somatosensory cortex (S1) of coronal slices (Bregma 0.14, -0.22, and -0.34). Scale bars: 100 *µ*m (left panels) and 20 *µ*m (magnifications of marked selections). **(i)** Quantification of the proportion of tdTomato$^+$ cells expressing PV in *Sst-*

*Cre/tdTomato* and *Sst-Cre/tdTomato/Dnmt1 loxP*$^2$ mice across the cortical layers normalized to the total amount of all detected tdTomato$^+$ cells. Nested two-way ANOVA with $n = 11$ slices for control and $n = 10$ for KO from $N = 3$ brains for both genotypes. $p < 0.05$ *, $p < 0.01$ **, $p < 0.001$ ***, n.s.: not significant. Data points with the same symbol belong to one mouse.

### Extended Data Figure E11:

**Neuropixels recordings in adult *Sst-Cre/tdTomato/Dnmt1-loxP*$^2$ and *Sst-Cre/tdTomato* mice**

**(a, b)** Histological localization of Neuropixels probes in V1 and S1. For *post-hoc* verification of the accurate positioning of the Neuropixels probes in V1 and S1 of *Sst-Cre/tdTomato/Dnmt1-loxP*$^2$ (*Dnmt1* KO) and *Sst-Cre/tdTomato* (control) mice, we covered each electrode with the fluorescent cell labeling solution DiD (Invitrogen, #V22887) before each recording. After all electrophysiological recordings were complete, brains were cut into 50 $\mu$m-wide brain slices and stained with DAPI to obtain an anatomical labeling (visible in blue). Based on anatomical landmarks, the slices were then aligned to the Allen common coordinate framework, and we verified that the probes were correctly positioned in S1 and V1. The figure shows examples of brain slices from S1 (left) and V1 (right) of a control **(a)** and *Dnmt1* KO mouse **(b)**. The fluorescent traces in red were found in comparable regions of S1 and V1, confirming similar and accurate targeting of cortical regions in both groups. **(c-h)** Adult *Dnmt1*-deficient mice show functional abnormalities. **(c)** Left and center panel: Average local field potential (LFP) in S1 across the whole cortical depth, following a 20-ms air puff stimulus (onset at white line) to the distal whisker pad. Blue colors indicate negative, and yellow colors positive LFP responses. Right panel: Across all layers, LFP responses were stronger and more temporally precise in control mice (gray) compared to KO mice (orange). **(d)** Quantification of spiking responses to tactile stimulation for neurons in the supra- and infragranular layers of S1 (see also Fig. 5i). Early spiking responses within the first 30 ms after stimulus onset were significantly reduced in KO mice, especially in the supragranular layers (early $\text{response}_{\text{control}}$ = 11.99 ± 0.86 Hz, early $\text{response}_{\text{KO}}$ = 4.96 ± 0.64 Hz, $p < 8.5^{-35}$, $n_{\text{control}}$ = 183 neurons from 2 mice, $n_{\text{KO}}$= 60 neurons from 2 mice). Supragranular tactile responses were also longer lasting in KO mice, with significantly increased late sensory responses from 100 to 300 ms (late $\text{response}_{\text{control}}$ = 2.09 ± 0.48 hz, late $\text{response}_{\text{KO}}$ = 5.47 ± 1.16 Hz, $p$ = 0. 0002). **(e)** To isolate the activity of SST$^+$ cINs, we used optogenetic stimulation with channelrhodopsin-2 to stimulate SST$^+$ cINs in V1 and S1 with 20-ms-long flashes of blue light (10 mW, 488 nm). The action potential waveform of light-responsive neurons was largely similar for control and KO mice, suggesting that developmental changes did not strongly alter action potential generation in SST$^+$ cINs. Correspondingly, the spike width was not significantly different for the two groups (right panel: spike $\text{width}_{\text{control}}$ = 1.04 ± 0.01 ms, spike $\text{width}_{\text{KO}}$ = 1.05 ± 0.03 ms, $p$ = 0. 8271, $n_{\text{control}}$ = 73 neurons from 2 mice, $n_{\text{KO}}$= 42 neurons from 2 mice). **(f)** Comparison of the spontaneous firing rate of the same SST$^+$ cINs as in (**c**) for control and KO mice. There was no significant difference in the firing rate of SST$^+$ cINs across the groups (firing $\text{rate}_{\text{control}}$ = 2.56

± 0.7 Hz, firing $rate_{KO}$ = 1.79 ± 1.12 Hz, $p$ = 0. 2958). **(g)** Left: Peristimulus time histogram (PSTH) for S1 and V1 neurons that were suppressed by optogenetic stimulation of $SST^+$ cINs in the same area for control (gray) and KO (orange) mice. The optogenetic stimulus started at time 0, ramped up for 0.5 seconds, and remained on for 1 second. Right: Optogenetic stimulation of $SST^+$ cINs equally suppressed other cortical neurons in both groups, suggesting that *Dnmt1* KO did not strongly alter the synaptic release of GABA in $SST^+$ cINs (firing rate $change_{control}$ = -4.69 ± 0.15 Hz, firing rate $change_{KO}$ = -4.65 ± 0.20 Hz, $p$ = 0. 8352, $n_{control}$ = 266 neurons from 2 mice, $n_{KO}$= 123 neurons from 2 mice). **(h)** Spontaneous firing rate of all cortical neurons across depth for control versus KO mice. The spontaneous firing rate of non-$SST^+$ cINs was significantly reduced in KO mice, especially in the supragranular layers (firing $rate_{control}$ = 1.25 ± 0.48 Hz, firing $rate_{KO}$ = 0.43 ± 0.52 Hz, $p$ = 0. 0015, $n_{control}$ = 262 neurons from 2 mice, $n_{KO}$= 93 neurons from 2 mice). Since the activity of $SST^+$ cINs was not strongly altered, these results suggest that functional aberrations in *Dnmt1* KO are due to structural changes in cortical network architecture rather than disrupted $SST^+$ cIN function. Significance for panels d, e, f, g, and was based on a Wilcoxon *rank-sum* test, values are mean ± SEM. Shading in panels c and g shows the SEM.

### Extended Data Figure E12:

#### Behavioral analyses of adult *Sst-Cre/tdTomato* and *Sst-Cre/tdTomato/Dnmt1 loxP²* mice.

(**a-d**) Sensory-driven decision-making was tested using an evidence accumulation task. Mice were stepwise trained on performing a two-alternative forced choice task, where they were required to accumulate either unisensory (visual/tactile) or multisensory (visuotactile) evidence over multiple seconds before indicating which site displayed more stimuli. (**a**) Learning curves of *Som-Cre/tdTomato* (control) and *SomCre/tdTomato/Dnmt1 loxP²* (*Dnmt1 KO*) mice for the three consecutively trained modalities display the percentages of correctly performed trials per session. Repeated performance values above 75% (red dashed lines) were indicative of successful learning. Comparing a similar time window for learning in each modality, no differences were found between control and knockout mice (linear mixed effects model (LME), $n$ = 18 binned sessions from $N$ = 6 mice per modality, $p_{visual}$ = 0.89, $p_{tactile}$ = 0.64, $p_{multisensory}$ = 0.29). (**b**) Multisensory enhancement, i.e. increased performances in trials with multisensory stimulation compared to trials with the preferred unisensory stimulation, was similarly expressed in mice of both genotypes (LME, $n$ = 98 sessions from N= 6 mice, $p$ = 0.95). (**c, d**) Psychometric curves for all three modalities indicate similar performances of *Som-Cre/tdTomato* (control) and *SomCre/tdTomato/Dnmt1 loxP²* (KO) mice across varying difficulty levels, based on the difference in stimuli presented on the left and right side of the animal. Side-specific (**c**) and -independent (**d**) performances revealed no genotype-specific effects (LME, $n$ = 18 performance values from $N$ = 6 mice over 3 difficulties, $p_{visual}$ = 0.78, $p_{tactile}$ = 0.33, $p_{multisensory}$ = 0.26). (**e-g**) Learning and visual navigation were tested in a Morris water maze (MWM) over six days. Animals were introduced into the water from alternating quadrants with visual landmarks indicating the position of the platform ($N$ = 7 *SomCre/tdTomato*, $N$ = 6 *SomCre/tdTomato/Dnmt1 loxP²*). **(e)** Scheme of the MWM-task. **(f)** Successful escapes from the MWM are shown as the percentage of trials in which the animals found the platform (binomial test: $p_{visual}$ = 0.32). **(g)** Covered distances per session are displayed genotypedependently (Two-way ANOVA: $p_{Genotype}$ = 0.238, $p_{Session}$ = $3.4^{-10}$, $p_{Genotype\ x\ Session}$ = 0.889). **(h, i)** Average timepoint and applied dosage of PTZ for provoking first occurrence of different forms of epileptic seizures following Racine's score: 4: continuous whole-body myoclonus indicated by a stretched back and extensive and dynamic stiffening of the tail with overhead pointing towards the rostrum. 5: clonic seizure characterized by unilateral forelimb clonus and bending over until reaching a sitting/standing position. 6: clonic-tonic seizure with falling on one side, wild rushing, and jumping. 7: generalized tonic-clonic seizure is indicated by wild rushing, jumping, and tonic extension cumulating in overstretched limbs and tail. Results in respiratory depression and/or death (end of experiment). *N* = 10 for both *Dnmt1*-genotypes, whereby the following showed respective epileptic seizure types: severity level 4 with *N* = 10 *SomCre/tdTomato* and *N* = 9 *SomCre/tdTomato/Dnmt1 loxP2*; severity level 5 with *N* = 10 *SomCre/tdTomato* and *SomCre/tdTomato/Dnmt1 loxP2* each; severity level 6 with N = 4 *SomCre/tdTomato* and *N* = 5 *SomCre/tdTomato/Dnmt1 loxP2;* severity level 7 with N = 9 *SomCre/tdTomato* and *N* = 7 *SomCre/tdTomato/Dnmt1 loxP2* **(h)** After application of the first PTZ injection (t = 0 min) *SomCre/tdTomato* and *SomCre/tdTomato/Dnmt1 loxP*[2] show similar timepoints of first occurrence regarding different Racine's score severity levels, respectively. **(i)** After application of the first PTZ injection (concentration = 10 mg PTZ per kg living weight) *SomCre/tdTomato* and *SomCre/tdTomato/Dnmt1 loxP2* show similar cumulated PTZ dosages to induce the first occurrence of different Racine's score severity levels, respectively. Unpaired, two-tailed Student's *t*-test and additional unpaired Welch's *t*-test; n.s.: not significant.

### 2. Extended Data Tables and Legends

**Extended Data Table E1**.

The number of contacts for A^2 of interface area (calculated by dr-sasa web server[2]) of selected protein/DNA complexes with affinities in the nM range. Affinity data from the ProNAB database[3]. All selected structures in solution (either by NMR or by the end of our three MD simulations), except for the DNMT1/UMDNA/SAH complex X-ray structure (PDB ID: 3PTA[4]).

| PDB ID | Kd (nM) | Number of contacts with a cutoff at 4 Å | Protein/DNA interfacial area (A^2) | Ratio |
|---|---|---|---|---|
| 1IV6[5] | 200.0 | 194 | 961 | 0.20 |
| 1J5N[6] | 10.0 | 296 | 1740 | 0.20 |
| 1MSE[7] | 11.2 | 275 | 1377 | 0.17 |
| 1GCC[8] | 4.1 | 164 | 800 | 0.21 |

| | | | | |
|---|---|---|---|---|
| 3PTA[4] | 2.3 | 97 | 761 | 0.13 |
| MD | 2.3 | 184±17 | 1102±54 | 0.17 |

**Extended Data Table E2.**

Interaction list between the catalytic domain and the CpG site in DNMT1/HMDNA/SAH complex was determined by Cryo-EM (PDB ID: 7XI9[9]) and by the end of our three MD simulations (residues within 4 Å).

| | 7XI9[9] | Replica1 | Replica2 | Replica3 |
|---|---|---|---|---|
| **LYS1275** | × | √ | × | √ |
| **ARG1276** | × | × | √ | √ |
| **MET1232** | √ | × | × | × |
| **ASN1233** | √ | × | × | × |
| **ARG1234** | √ | √ | √ | √ |
| **ASN1236** | √ | √ | × | √ |

**Extended Data Table E3**

The atomistic partial charges of SAM (**Supplementary Figure 1**) in this study.

| Atom | Charge | Atom | Charge | Atom | Charge | Atom | Charge | Atom | Charge |
|---|---|---|---|---|---|---|---|---|---|
| N1 | -0.486 | O11 | -0.382 | C21 | 0.602 | H31 | 0.400 | H41 | 0.122 |
| C2 | 0.024 | C12 | 0.159 | C22 | -0.062 | H32 | 0.245 | H42 | 0.159 |
| C3 | 0.661 | O13 | -0.650 | C23 | 0.682 | H33 | 0.245 | H43 | 0.0889 |
| C4 | -0.689 | C14 | 0.152 | N24 | -0.807 | H34 | 0.164 | H44 | 0.096 |
| C5 | 0.032 | O15 | -0.618 | N25 | -0.546 | H35 | 0.164 | H45 | 0.112 |
| C6 | -0.249 | C16 | 0.139 | C26 | 0.154 | H36 | 0.193 | H46 | 0.057 |
| C7 | 0.337 | O17 | -0.689 | N27 | -0.186 | H37 | 0.193 | H47 | 0.134 |
| C8 | -0.355 | N18 | -0.770 | H28 | 0.451 | H38 | 0.193 | H48 | 0.347 |
| C9 | -0.450 | C19 | 0.548 | H29 | 0.424 | H39 | 0.067 | H49 | 0.347 |
| C10 | 0.219 | N20 | -0.785 | H30 | 0.400 | H40 | 0.067 | H50 | 0.347 |

**Extended Data Table E4.**

Parameters of Zn(II) ions (**Supplementary Figure 1**) in this study.

| | Type | LJ Radius (Å) | LJ Depth (kcal/mol) | Charge (e) | GB Radius (Å) | GB Screen |
|---|---|---|---|---|---|---|
| Zn1 | Cys1476-Cys1478-Cys1485-His1502 | 1.3730 | 0.0118 | 0.6157 | 1.5000 | 0.8000 |
| Zn2 | Cys653-Cys656-Cys659-Cys691 | 1.3730 | 0.0118 | 0.8827 | 1.5000 | 0.8000 |
| Zn3 | His793-Cys820-Cys893-Cys896 | 1.3730 | 0.0118 | 0.6757 | 1.5000 | 0.8000 |
| Zn4 | Cys664-Cys667-Cys670-Cys686 | 1.3730 | 0.0118 | 0.6654 | 1.5000 | 0.8000 |

**Extended Data Table E5**.

Simulated system in this study.

| System | Charge of complex | Number of $K^+$ ions | Number of $Na^+$ ions | Number of $Cl^-$ ions | Number of $H_2O$ molecules |
|---|---|---|---|---|---|
| DNMT1/UMDNA/SAM | -21 | 157 | 14 | 150 | 39,570 |
