## Extended data Figures for "DNMT1-Mediated Regulation of Somatostatin-positive Interneuron Migration Impacts Cortical Architecture and Function"

### Extended Data Figure E1

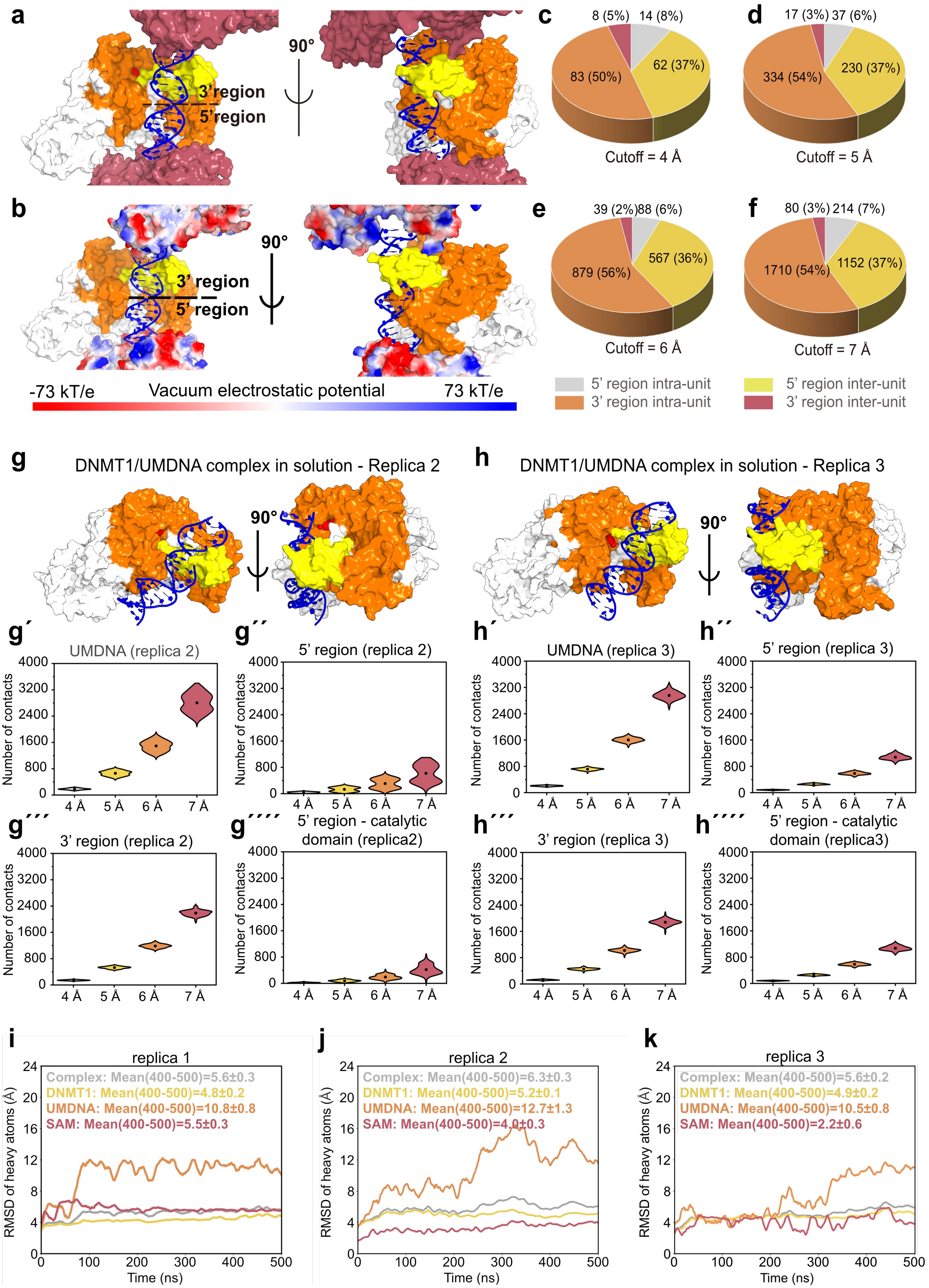

### Extended Data Figure E2

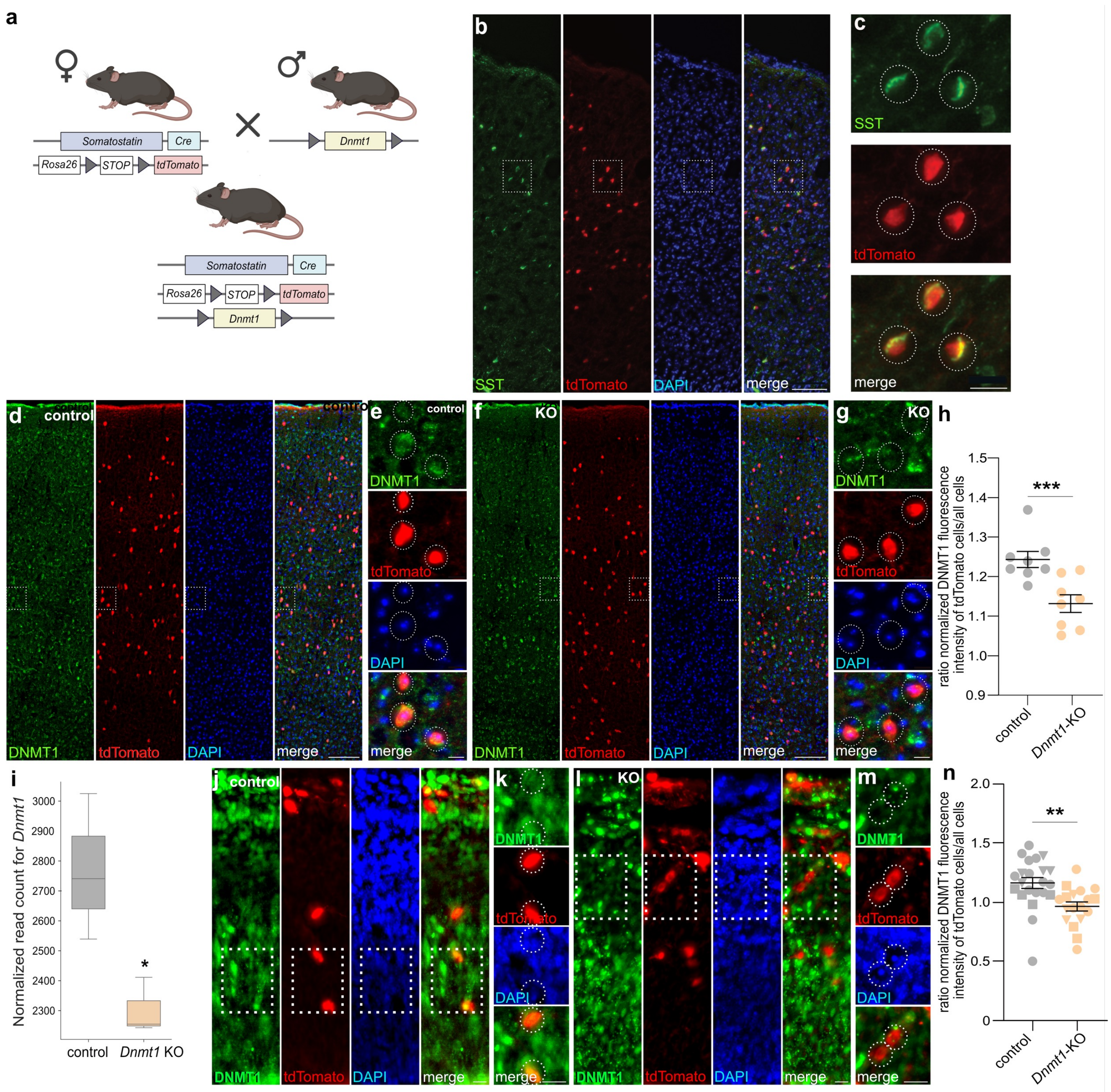

### Extended Data Figure E3

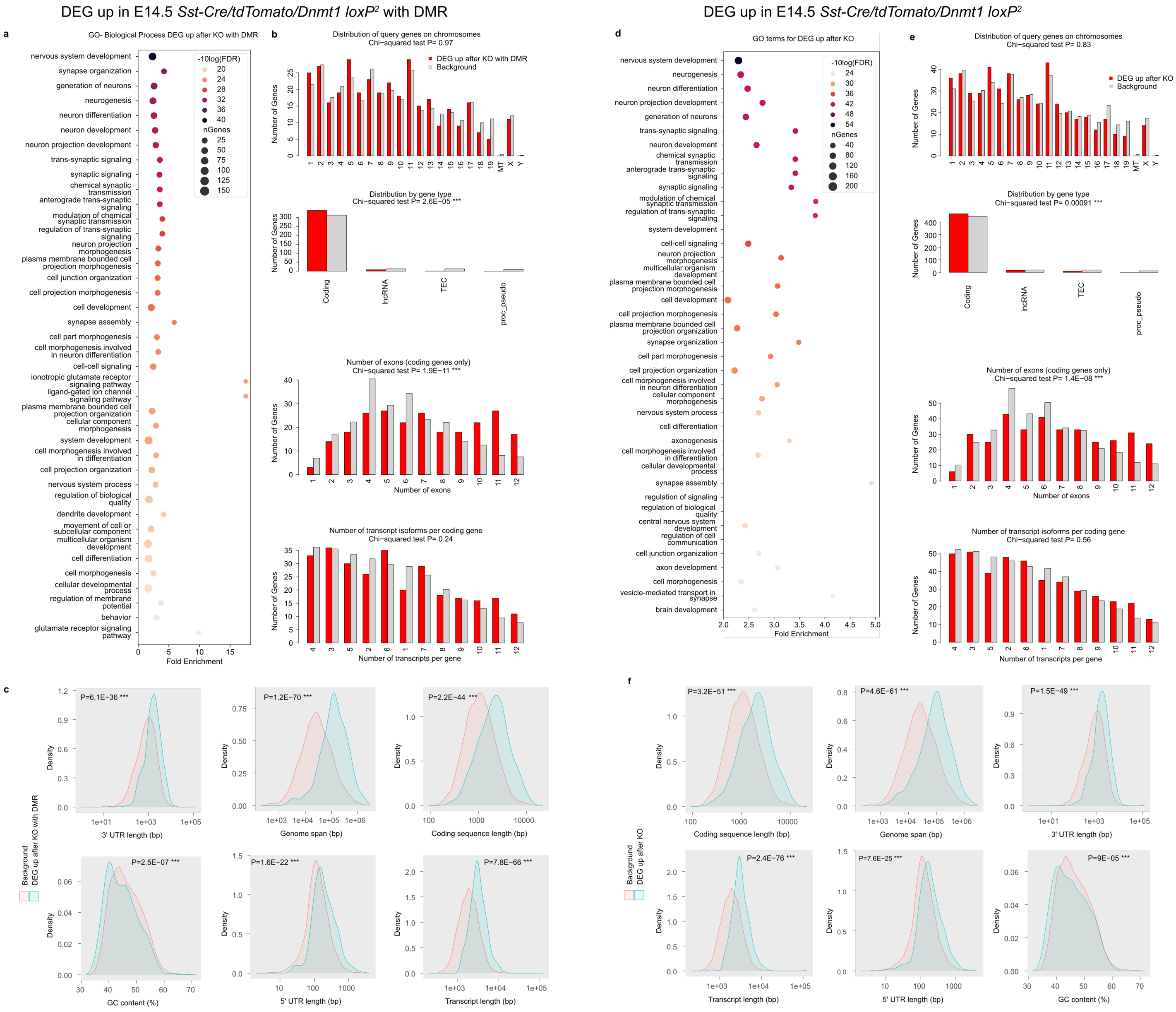

### Extended Data Figure E4

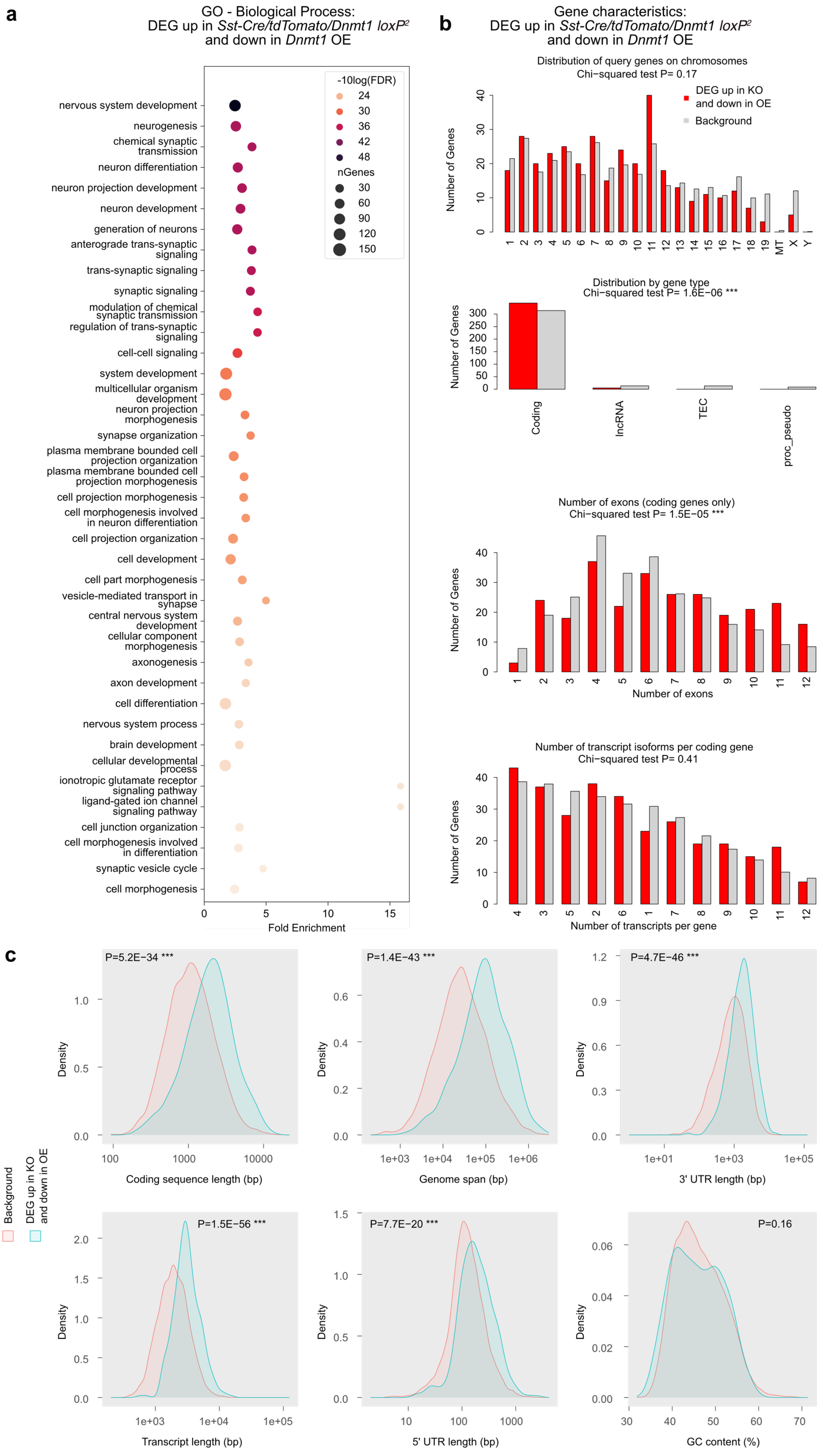

### Extended Data Figure E5

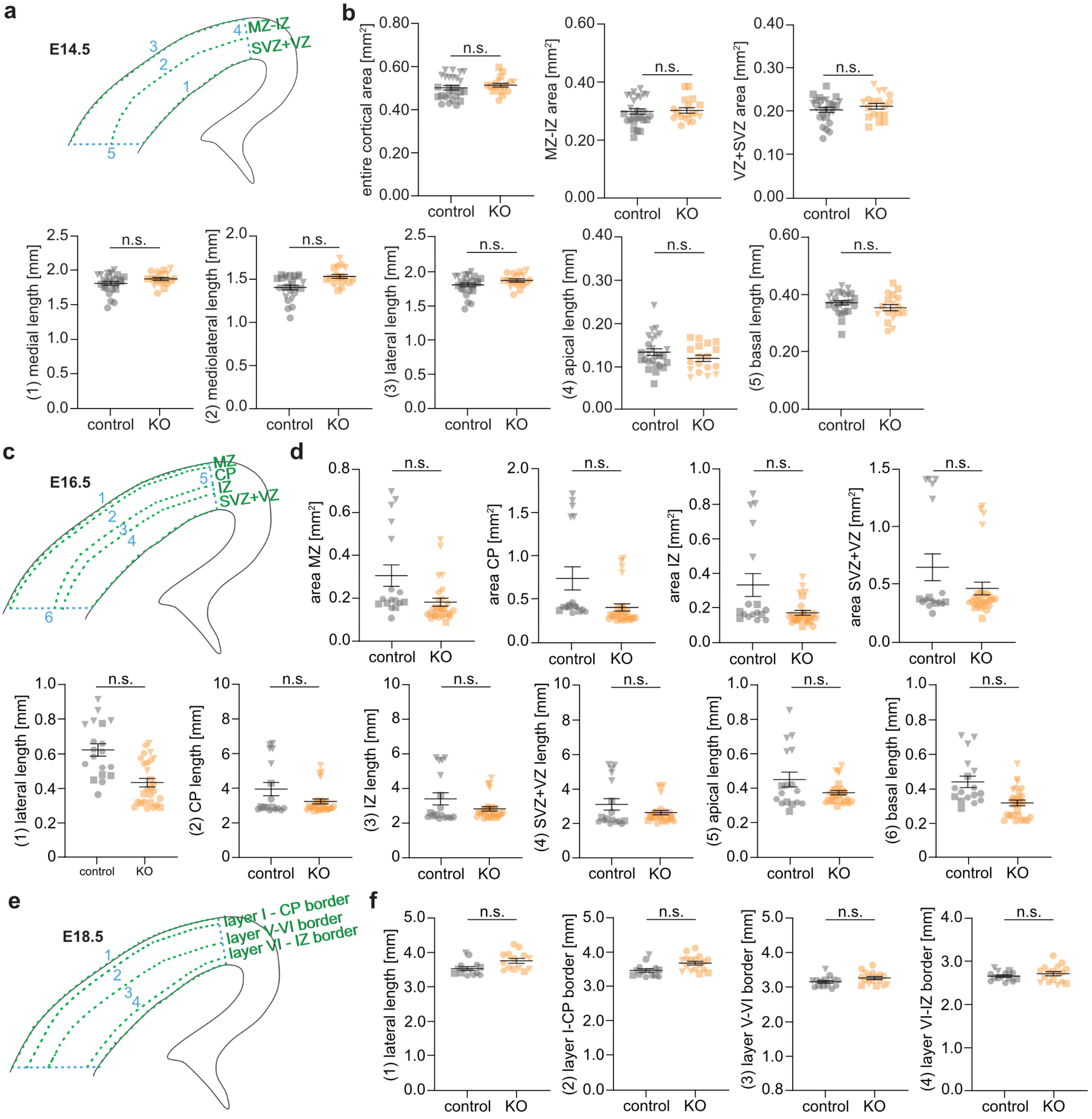

### Extended Data Figure E6

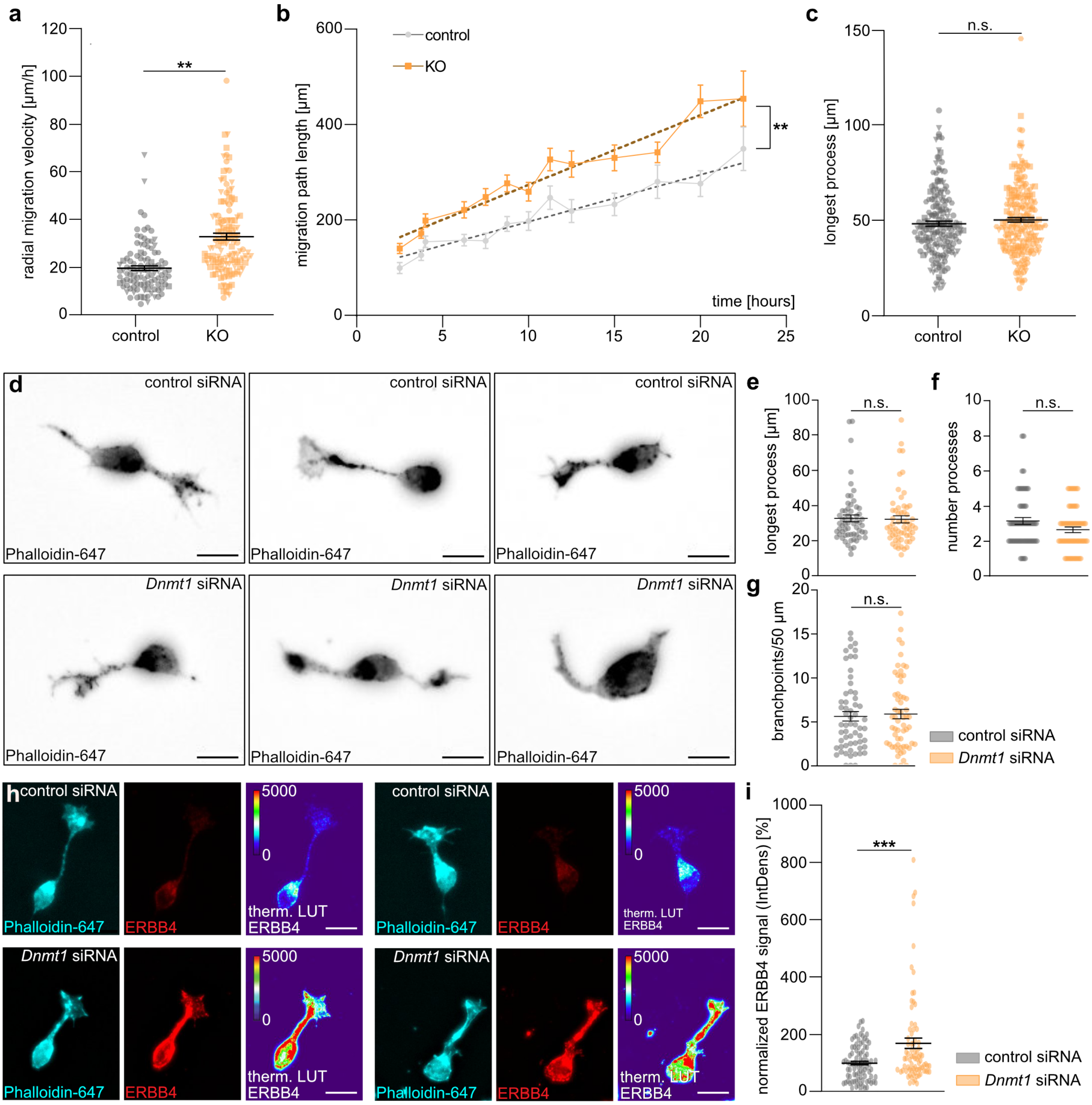

### Extended Data Figure E7

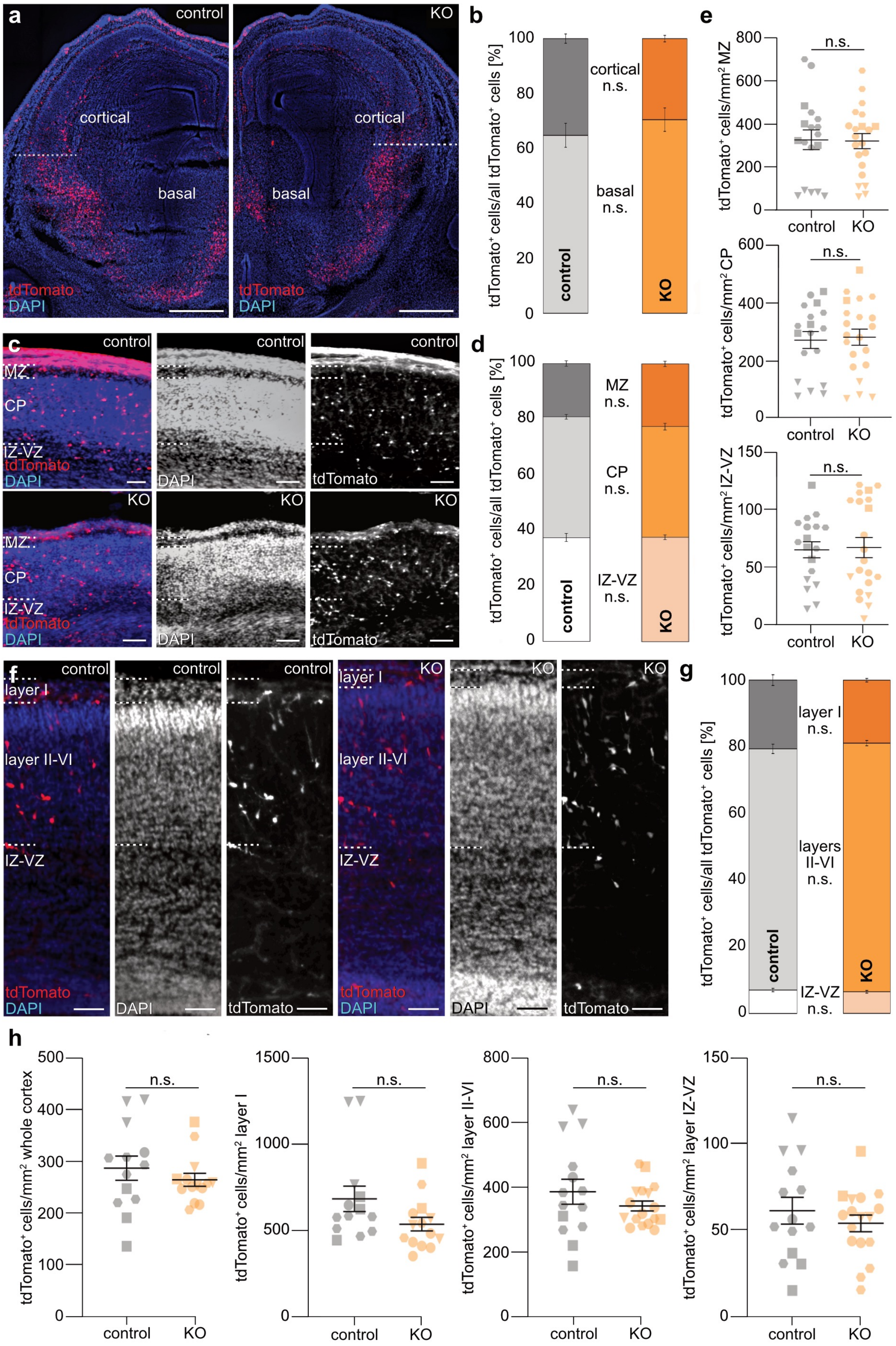

### Extended Data Figure E8

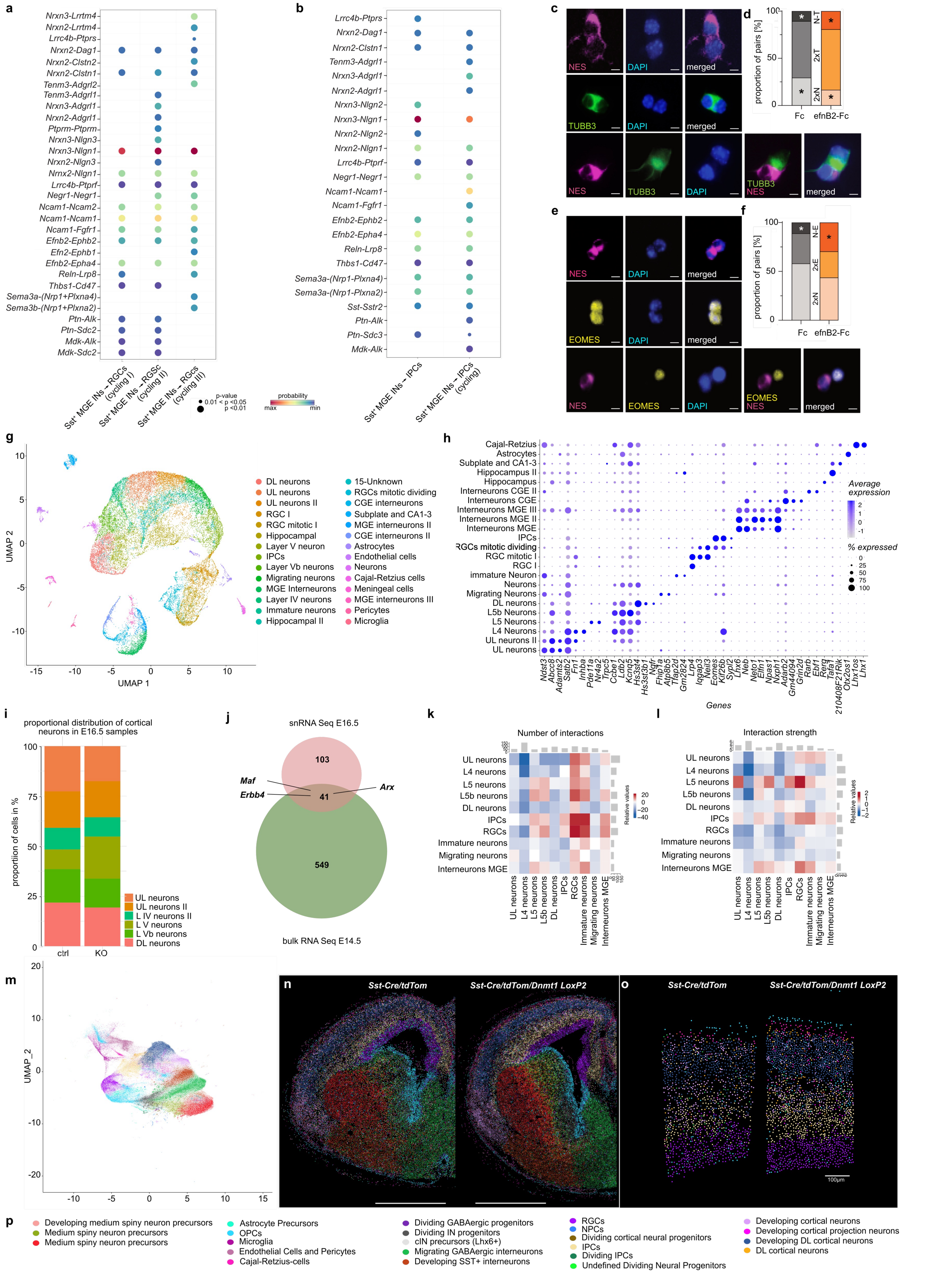

### Extended Data Figure E9

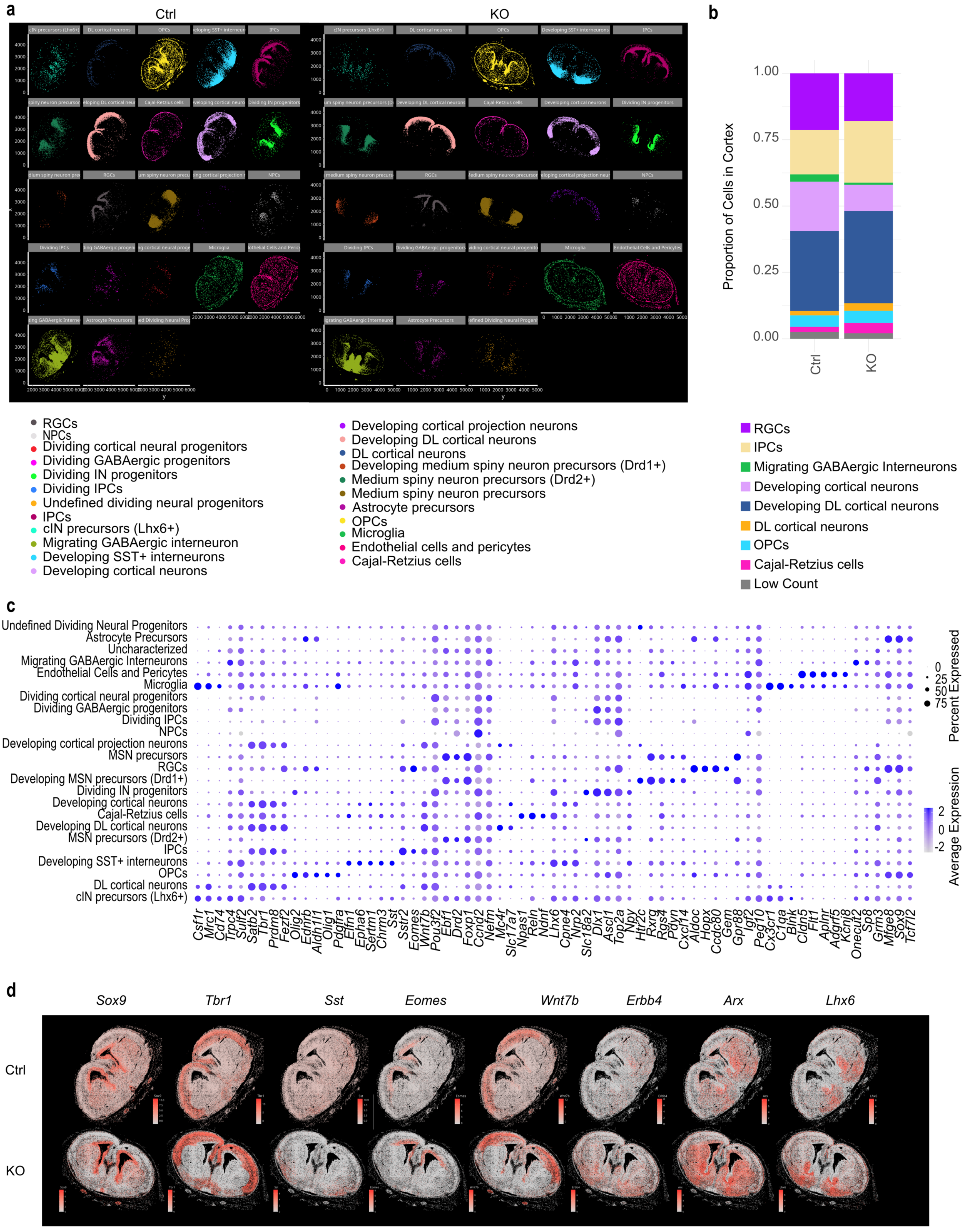

### Extended Data Figure E10

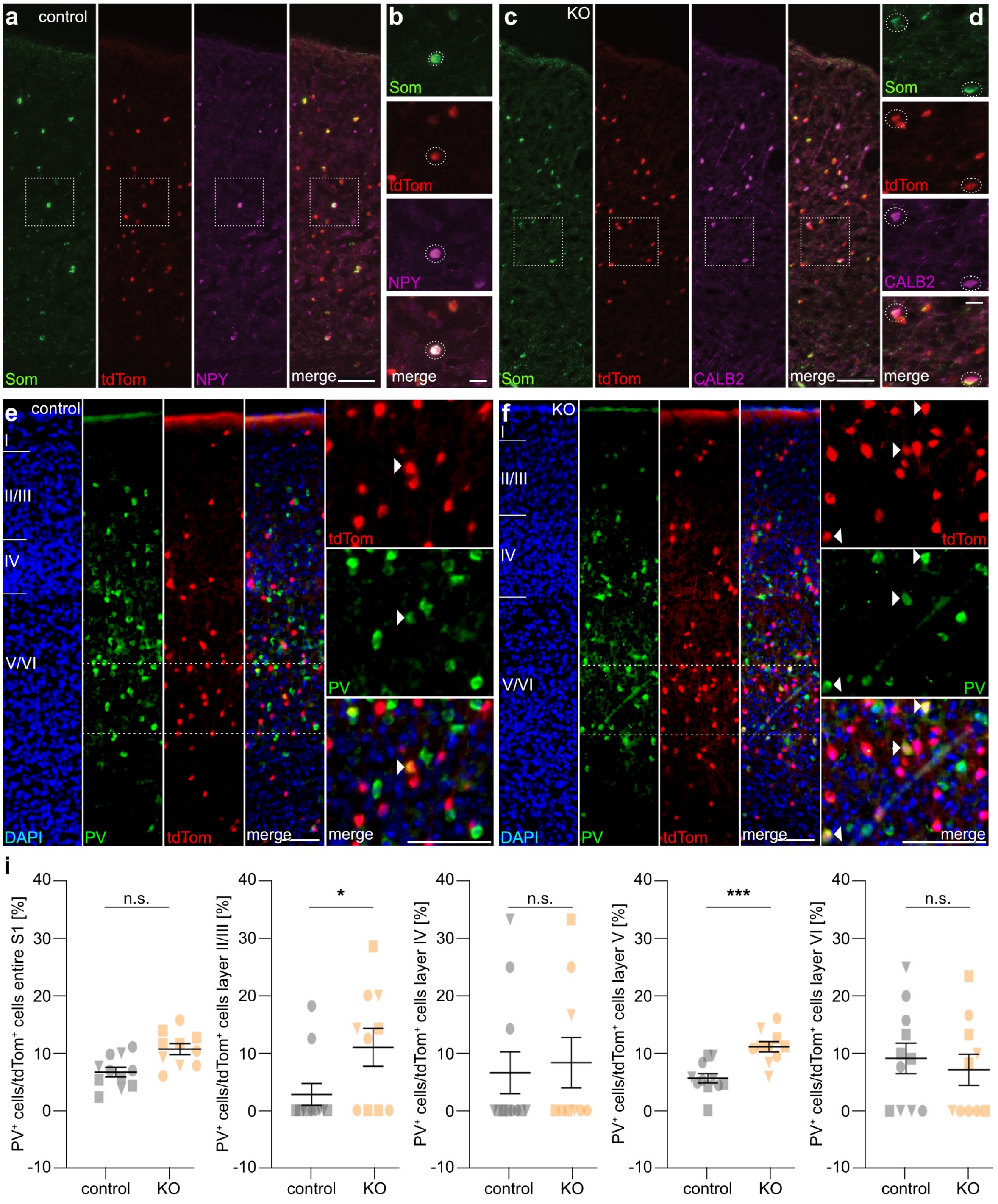

### Extended Data Figure E11

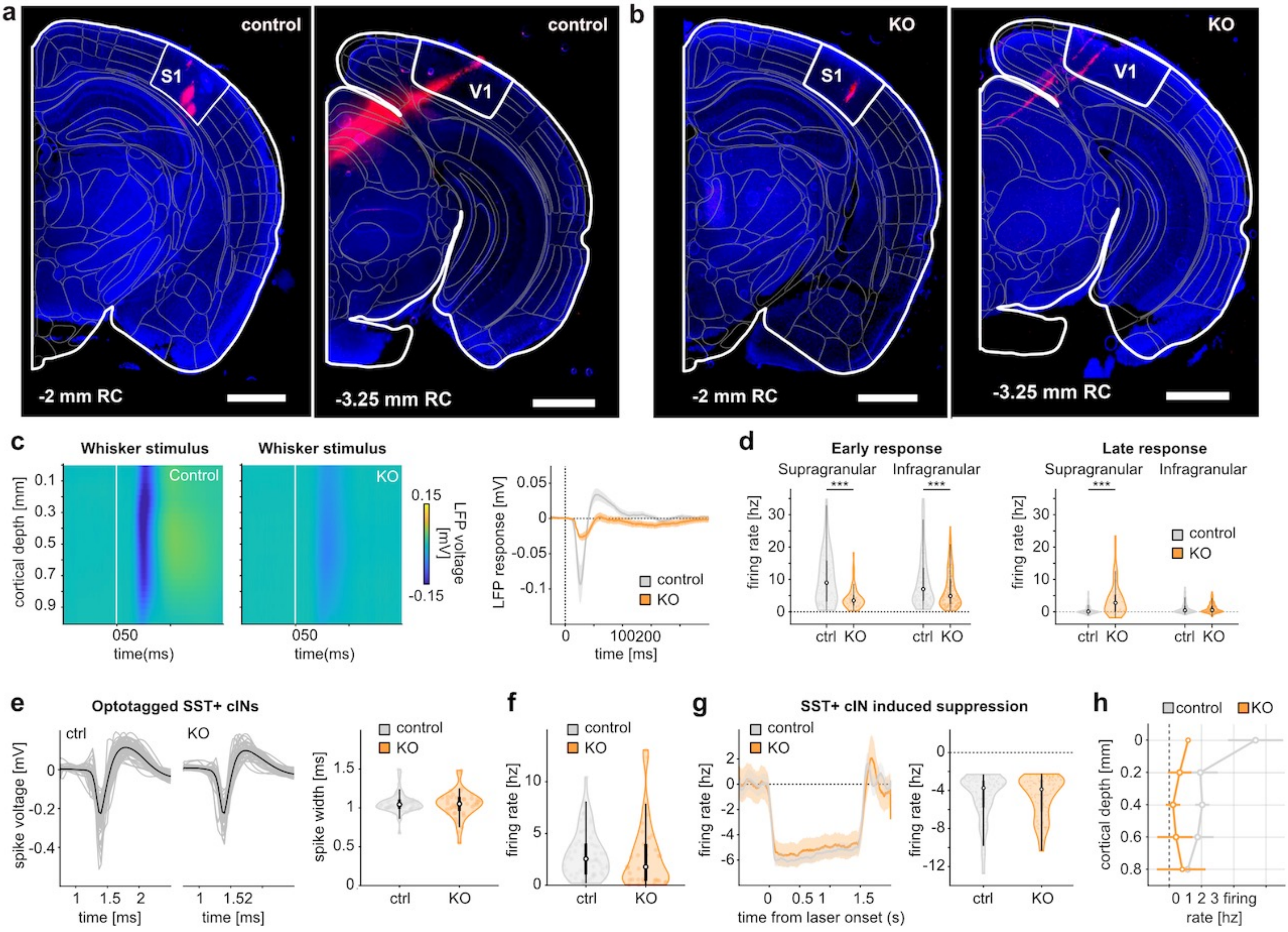

### Extended Data Figure E12

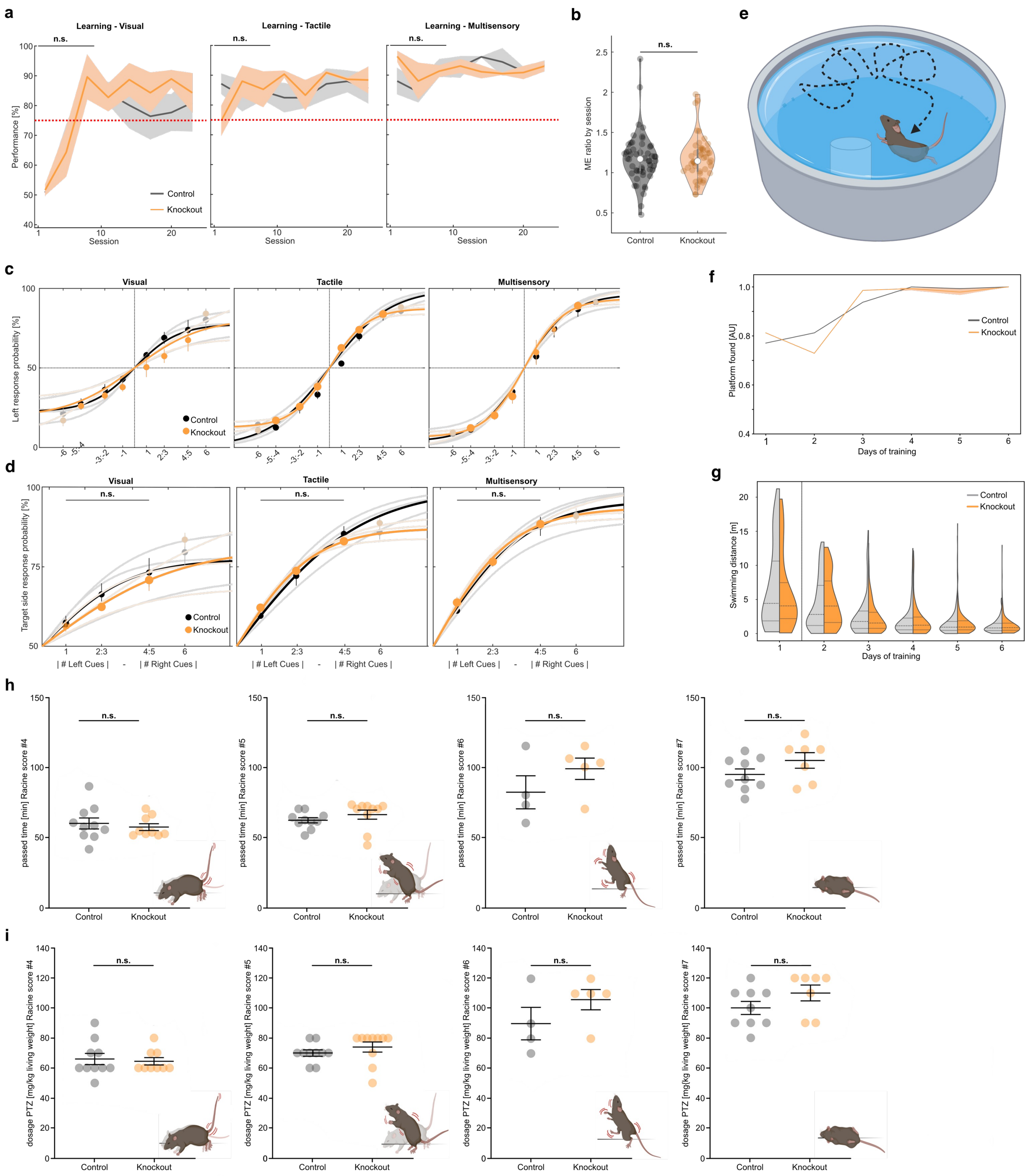
