## Supplementary Information for "DNMT1-Mediated Regulation of Somatostatin-positive Interneuron Migration Impacts Cortical Architecture and Function"

**Supplementary Table legends**

**Supplementary Table 1. (Supplementary table1-RNA Seq FACS SST-KO and ctrl E14.xlsx)**
Comparison of differential gene expression in tdTomato-positive interneurons of the basal telencephalon of E14.5 *Sst-Cre/tdTomato/Dnmt1 loxP*$^{2}$ and *Sst-Cre/tdTomato* control mice as revealed by RNA sequencing. Differentially expressed genes (DEG) are listed in sheet 1-1, genes with reduced expression in *Sst-Cre/tdTomato/Dnmt1 loxP*$^{2}$ are listed in sheet 1-2, while genes found upregulated in *Sst-Cre/tdTomato/Dnmt1 loxP*$^{2}$ are listed in sheet 1-3.

**Supplementary Table 2. (Supplementary table2-DMRs FACS SST-KO and ctrl E14.xlsx)**
Differentially methylated regions in tdTomato-positive interneurons of the basal telencephalon of E14.5 *Sst-Cre/tdTomato/Dnmt1 loxP*$^{2}$ and *Sst-Cre/tdTomato* control mice as revealed by methyl-sequencing. Differentially methylated regions (DMRs) are listed in sheet 1-1. Negative values indicate increased methylation in KO.

**Supplementary Table 3. (Supplementary table3-Overlap DEG-DMG SST-Dre-DNMT1 E14.5Venn_Data.xlsx)**
Collection of genes that were both differentially expressed and differentially methylated in tdTomato-positive interneurons of the basal telencephalon of E14.5 *Sst-Cre/tdTomato/Dnmt1 loxP*$^{2}$ and/or *Sst-Cre/tdTomato* control mice as revealed by RNA and methyl-sequencing, respectively, building the basis of the Venn diagrams depicted in Figure 1. DEG_up refers to upregulated genes in *Sst-Cre/tdTomato/Dnmt1 loxP*$^{2}$ samples.

**Supplementary Table 4. (Supplementary Table 4 – ChIP seq THOR peaks p1.4.xlsx)**
DNMT1 ChIP-seq peaks in immortalized cerebellar granule (CB) cells. The table lists the chromosomal locations of DNMT1-interacting chromatin segments with a *p* value cut-off of 1.4 (-log10) as well as the corresponding genes on the coding and the template strand, marked by (+) and (-), respectively. The peaks were generated by the THOR algorithm with the default parameters[1].

**Supplementary Table 5. (Supplementary table5-GO_enrichment_DEGup_and_DMR.csv)**
Gene ontology (GO) – biological process analysis of genes with increased expression and reduced methylation in E14.5 *Sst-Cre/tdTomato/Dnmt1 loxP*$^{2}$ samples compared to *Sst-Cre/tdTomato* cells.

**Supplementary Table 6. (Supplementary table6-GO_BiologicalP_DEG_up.csv)**
Gene ontology (GO) – biological process analysis of all genes with increased expression in E14.5 *Sst-Cre/tdTomato/Dnmt1 loxP*$^{2}$ samples compared to *Sst-Cre/tdTomato* cells.

**Supplementary Table 7. (Supplementary table7-Overlap DEG SST-Cre DNMT1 KO withDNMT1 OE)**
Collection of genes that were differentially expressed in both E14.5 *Sst-Cre/tdTomato/Dnmt1 loxP²* samples compared to *Sst-Cre/tdTomato* cells, as well as in *Dnmt1* overexpressing neurons transdifferentiated from murine ESCs compared to their controls. This table also provides the basis for the Venn Diagram depicted in Figure 1.

**Supplementary Table 8. (Supplementary table8DEG_up_DNMT1OE_down_enrichment.csv)**
Gene ontology (GO) – biological process analysis of all genes with increased expression in E14.5 *Sst-Cre/tdTomato/Dnmt1 loxP²* samples compared to *Sst-Cre/tdTomato* cells as well as reduced expression in *Dnmt1* overexpressing neurons transdifferentiated from murine ESCs compared to their controls.

**Supplementary Table 9. (Supplementary table9-DEG bTel and Ctx and overlap)**
Collection of genes that were differentially expressed between E14.5 *Sst-Cre/tdTomato/Dnmt1 loxP²* and *Sst-Cre/tdTomato* cells prepared from the cerebral cortex. Moreover, the overlap of significantly upregulated genes in *Sst-Cre/tdTomato/Dnmt1 loxP²* cells from the basal telencephalon and from the cortex are summarized. Moreover, gene ontology analysis (Biological process) details are depicted, performed with the set of genes that were commonly upregulated in FAC-sorted *Sst-Cre/tdTomato/Dnmt1 loxP²* neurons from both compartments (basal telencephalon and cortex). Background is defined as all detected transcripts in both datasets (ShinyGO 0.81; http://bioinformatics.sdstate.edu/go/).

**Supplementary movie legends**

**Supplementary Videos 1, 2**
Migratory behavior of interneurons depicted in organotypic brain slices (350 $\mu$m) of E14.5 *Sst-Cre/tdTomato* (Suppl. Video 1) and *Sst-Cre/tdTomato/Dnmt1 loxP²* Suppl. Video 2) embryos. Footages display an imaging period of 20 hours with 15 minutes per frame.

TdTomato+ cells are depicted in grey. Scale bar: 100 μm. MZ: marginal zone, CP: cortical plate, VZ: ventricular zone.

**Supplementary Videos 3-9**

Seizure events during PTZ-induced convulsions in three-month-old *Sst-Cre/tdTomato*) and *Sst-Cre/tdTomato/Dnmt1 loxP*[2] mice. All videos are depicted in real-time. Videos 6 and 7 display examples of small seizures of event I (whole body myoclonus until reaching a standing position with an upward stretched tail). In videos 8 and 7 mice are displaying small seizure events of type II (whole-body myoclonus with stretched body and stretched tail pointing towards the head). Event III of seizure types is depicted in video 10, characterized by several convulsions leading to short rearing episodes, falling over, and rolling with subsequent recovery of the individual. Videos 11 and 12 show examples of final tonic-clonic seizures (event IV) with whole-body convulsions culminating in heavy rearing and overstretched limbs.

**Supplementary Video 10**

Stereotypic behavior of an adult *Sst-Cre* strain mouse depicting repetitive movements characterized by mounting on the play tunnel, flipping over, and instant repetition of the procedure. Stereotypic behavior can indicate general vulnerability to stress or autism-like symptoms. Footages are depicted in real-time.

### Supplementary Methods

### Supplementary Methods

#### Model Construction

Our structure model of the DNMT1/UMDNA/SAM complex was based on the X-ray structures of DNMT1/UMDNA/SAH (PDB ID: 3PTA[2]) and of DNMT1/SAM (PDB ID: 3AV6) [3]. The missing loops in DNMT1 were modeled using the SWISS-MODEL web server[4]. The protonation states of DNMT1's ASP, GLU, ARG, LYS, and HIS residues, as well as those of the UMDNA strand (5'-TpCpCpCpGpTpGpApGpCpCpTpCpCpGpCpApGpGp-3'), were determined using the H++ web server[5] and tLEaP based on the Amber OL21 force field[6] assuming a pH 7.4 to mimic physiological conditions.

The force fields for the protein, methylated cytosine, UMDNA, water, and ions were AMBER ff19SB force fields[7], parameter set from Lankas et al.[8], Amber OL21 force field[6], OPC[9], and Åqvist potential[10], respectively. Those of SAM (**Extended data Figure E1l, and Extended Data Table E3**) and the Zn coordination polyhedron (**Extended data Figure E1m, and Extended Data Table E4**) were consistent with the used protein force field: The charges were derived from the geometry-optimized structure at the B3LYP level of theory (with Grimme's dispersion correction[11]). The basis sets were 6-311G(d,p) and 6-31G(d), respectively. We used the restrained electrostatic potential (RESP) fitting method[12], following the Merz-Kollman (MK) scheme[13]. The van der Waals parameters were taken from the previous work of Li et al.[14] .

The bonded parameters of the Zn coordination polyhedron were calculated at the same level of theory and obtained by the Seminario method[15]. Those for SAM were from GAFF2[16]. All the quantum mechanics (QM) calculations are performed by Gaussian09[17].

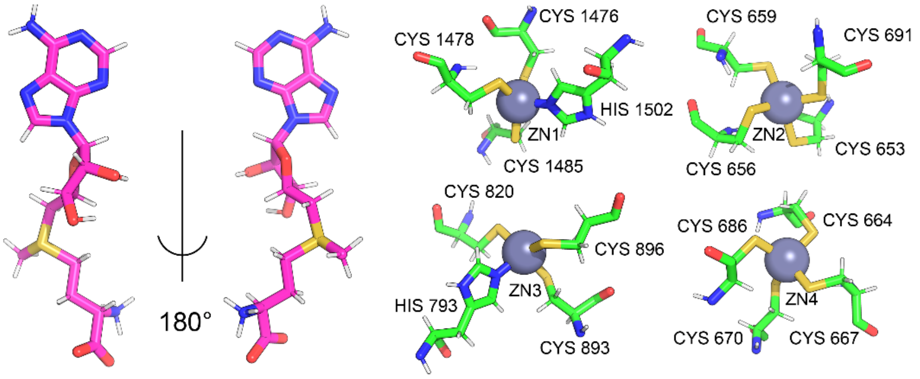


**Supplementary Figure 1:** Three-dimensional schematic diagram of SAM (left) and Zn coordination polyhedron (right). The parameters of SAM and Zn(II) ions are provided in **Extended Data Tables E3 and E4**.

**Details on generating transgenice mice:**

Generation of control animals was conducted by crossing a $Sst^{tm2.l(cre)Zjh}$/J strain (RRID:IMSR_JAX:013044, Jackson Laboratory, Bar Harbour, U.S.A.) with a t*dTomato* reporter mouse model (*B6.CgGt(ROSA)26Sor*$^{tml\ 4(\ CAG\text{-}tdTomato\ )Hze}$, obtained from Christian Hübner, University Hospital Jena, Germany). An *internal ribosomal entry site* (*IRES*), a *Cre-recombinase* sequence, a *polyA* sequence, and a *frt*-flanked neo cassette were inserted into the 3' untranslated region (UTR) of the *Somatostatin (Sst)* locus on chromosome 16, limiting the respective *Cre*-expression to *Sst*-positive cells. The t*dTomato* reporter strain harbors the *tdTomato* sequence combined with a *loxP*-flanked stop cassette in the *Rosa26* locus. Expression of *Cre-recombinase* dependent on *Sst*-promoter activity resulted in the deletion of the *loxP*-flanked stop cassette during recombination and an expression of *tdTomato* in *Sst-Cre*-positive cells. *Jackson Laboratory* recommends creating and breeding *Sst-Cre* heterozygous mice, which received the *Cre*-allele from the maternal side. This strategy avoids potential instability of the Cre-recombinase activity during paternal germline recombination[18]. To generate the triple transgenic conditional KO-model *Sst-Cre*$^{+/-}$*/tdTomato* females were bred with male individuals derived from crossing *tdTomato* mice with a *Dnmt1 loxP*$^{2}$ strain (*B6; 129Sv-Dnmt1tm4Jae/J,* obtained from Rudolf Jaenisch, Whitehead Institute for Biomedical Research Boston, U.S.A.) in which exon 4 and 5 of the *Dnmt1* gene are loxP-flanked, resulting in a null allele of these loci and a subsequent DNMT1 deficiency[19]. Corresponding programs and primers used for genotyping can be found in the supplementary information. An illustration of control- and KO mice is shown in Extended Data Figure E2a.

**DNA extraction and genotyping**

Ear biopsies as well as embryonic tissue and tail biopsies from mice killed in experiments were incubated in an alkaline lysis buffer (25 mM NaOH, 0.2 mM EDTA, volume adapted to tissue size) for 90 minutes at 96°C and 350 rpm using a ThermoMixer® (Eppendorf AG, Germany).

Subsequently, samples were cooled down on ice and neutralized using the same volume of 40 mM Tris-HCl. Next, amplification of genomic DNA was performed in a T100 thermal cycler (BioRad, U.S.A.) with the help of the corresponding primer sequences and PCR programs listed below. For this, 1 $\mu$L of isolated DNA was incubated together with 19 $\mu$L of PCR master mix containing 2x FastGene™ Optima reaction mix (Nippon Genetics, Japan), nuclease-free $H_2O$, and respective primers. Product size determination via electrophoresis was conducted using a 2% agarose gel (2% agarose/1x Tris-acetate-EDTA (TAE) buffer) containing MidoriGreen™ (Nippon Genetics, Japan). Additionally, a 1 kb DNA ladder was applied according to the manufacturer's guidelines (GeneRuler™, Thermo Fisher Scientific, U.S.A.). Gel electrophoresis was performed applying 190 V using a PowerPac power supply (BioRad, U.S.A.). Final detection of genotyping results was conducted via GelDoc™XR imaging system (BioRad, U.S.A.) and the corresponding software Image Lab (BioRad, U.S.A.).

**Primer sequences and polymerase chain reaction (PCR) programs used for genotyping**

*Somatostatin-Cre*

- Universal reverse primer: 5'-GGG CCA GGA GTT AAG GAA GA-3'
- Wildtype forward primer: 5'-TCT GAA AGA CTT GCG TTT GG-3'
- Mutant forward primer: 5'-TGG TTT GTC CAA ACT CAT CAA-3'

*TdTomato*

- Wildtype forward primer: 5'-AAG GGA GCT GCA GTG GAG TA-3'
- Wildtype reverse primer: 5'-CCG AAA ATC TGT GGG AAG TC-3'
- Mutant forward primer: 5'-GGC ATT AAA GCA GCG TAT CC-3'
- Mutant reverse primer: 5'-CTG TTC CTG TAC GGC ATG G-3'

*Dnmt1 loxP*[2]

- Forward primer: 5'-GGG CCA GTT GTG TGA CTT GG-3'
- Reverse primer: 5'-CCT GGG CCT GGA TCT TGG GGA-3'

| | |
|---|---|
| | 94 °C, 5 min |
| ⌈ | 94 °C, 30 s |
| 9x | 65 °C, 30 s |
| ⌊ | 72 °C, 30 s |
| ⌈ | 94 °C, 30 s |
| 40x | 60 °C, 30 s |
| ⌊ | 72 °C, 30 s |
| | 72 °C, 5 min |
| | 4 °C, ∞ |

| | |
|---|---|
| | 95 °C, 3 min |
| ⌈ | 95 °C, 15 s |
| 9x | 61 °C, 20 s |
| ⌊ | 72 °C, 30 s |
| | 72 °C, 3 min |
| | 12 °C, ∞ |

| | |
|---|---|
| | 95 °C, 3 min |
| ⌈ | 95 °C, 30 s |
| 37x | 58 °C, 25 s |
| ⌊ | 72 °C, 30 s |
| | 72 °C, 2 min |
| | 4 °C, ∞ |

**Chromatin immunoprecipitation (ChIP) and sequencing**

Cerebellar granule cells[20] were cultured and treated as previously described[21]. DNMT1-interacting chromatin was immunoprecipitated via native chromatin immunoprecipitation (ChIP)[21]. The sequencing libraries were prepared using the NEBNext® Ultra II DNA Library Prep kit (#E7410S, New England Biolabs, U.S.A.) as per manufacturer's instructions. The input controls were pooled across samples in equimolar proportions. The libraries were amplified via PCR on a T100 Thermal Cycler (Bio-Rad, U.S.A.) with the following conditions: initial denaturation at 98°C for 30 s, 15 cycles of denaturation at 98°C for 10 s and annealing/extension at 65°C for 75 s, followed by a final extension at 65°C for 5 min. To account for the laddering caused by the enzymatic digestion, the libraries were size selected on a 2% (v/v) agarose gel and purified using the QIAquick Gel Extraction Kit (#28704, Qiagen, Germany). The libraries were sequenced on the NextSeq platform using 75 bp single-end reads at the IZKF Genomics Facility (University Hospital Aachen).

FASTQ files generated from the ChIP-seq experiment were processed using the nf-core/chipseq pipeline (version 2.0.0) with the mm10 reference genome[22]. Differential peak calling between the samples was performed using THOR with default settings[23]. The identified peaks with -log10*p* > 1.4 in the control samples are listed in Supplementary Table 4. Sequences within a 500 bp window surrounding these peaks were analyzed for motif detection using MEME-ChIP[24].
